# Epigenetic and transcriptional regulation of ovarian development altered in *Erβ*^KO^ ovaries

**DOI:** 10.1101/2024.08.21.608994

**Authors:** Ryan Mohamadi, Kevin Vo, Yashica Sharma, Amelia Mohamadi, Elizabeth S. Bahadursingh, Patrick E. Fields, M. A. Karim Rumi

**Affiliations:** Department of Pathology and Laboratory Medicine, University of Kansas Medical Center, Kansas City, KS 66160, USA

**Keywords:** Ovarian follicles, postnatal development, transcriptome, epigenetic regulators, transcription factors, Estrogen receptor β, deletion mutants

## Abstract

We analyzed the transcriptome of wildtype and estrogen receptor β knockout (Erβ^KO^) rat ovaries during the early postnatal period and detected remarkable changes in epigenetic regulators and transcription factors. Compared to postnatal day (PD) 4.5 wildtype ovaries, 17 differentially expressed epigenetic regulators (DEERs), and 23 differentially expressed transcription factors (DETFs) were detected in PD6.5 wildtype ovaries. Subsequently, compared to PD 6.5 wildtype ovaries, 24 DEERs and 68 DETFs were detected in PD8.5 ovaries. Changes in DEERs and DEFTs resulted in 581 differentially expressed downstream genes (DEDGs) in PD6.5 and 920 DEDGs in wildtype PD8.5 ovaries. The DEERs, DETFs, and DEDGs in wildtype ovaries represented primordial follicle activation (PFA) and development of the first-wave follicles because the second-wave follicles remain dormant during this period. However, the changes in DEERs, DETFs, and DEDGs during this postnatal period were markedly different in Erβ^KO^ rat ovaries, which suffered from increased PFA in both waves. Compared to 17 DEERs and 23 DETFs in wildtype, 46 DEERs and 55 DETFs were identified in PD 6.5 Erβ^KO^ ovaries. The differences were more remarkable in PD 8.5 Erβ^KO^ ovaries; compared to 24 DEERS and 68 DETFs in wildtype, only 8 DEERs and 10 DETFs were detected in Erβ^KO^ ovaries. Such dysregulation resulted in altered DEDGs in PD 6.5 (581 vs. 744) and in PD8.5 (920 vs. 191) Erβ^KO^ ovaries. These findings also suggest that the number of DEDGs depends directly on the numbers of DEERS and DETFs. In addition to the quantitative differences in DEERs and DETFs between the wildtype and Erβ^KO^ ovaries, we detected distinct differences in the identities of the regulators. Our observations indicate that loss of ERβ dysregulates the epigenetic regulators and transcription factors in Erβ^KO^ ovaries, which disrupts the downstream genes in ovarian follicles and increases follicle activation.

## 1. Introduction

Ovarian follicles are the functional units of the ovary that accomplish steroidogenesis and oogenesis[1]. The first step in ovarian follicle development is oocyte nest breakdown coupled with primordial follicle assembly[2,3]. Primordial follicles in the mouse or rat ovary are formed in distinct waves in different ovary regions and show different developmental dynamics[3-6]. The first wave of primordial follicles is developed in the medulla at birth and activated rapidly[3-6]. In contrast, the second wave is formed during postnatal days (PD) 4.5 to 8.5 and remains dormant[3-7]. The dormant follicles of the second wave establish the ovarian reserve for the entire reproductive period and determine ovarian longevity[3,5]. In contrast, the first-wave follicles are activated immediately after assembly, and their roles remain largely unclear [4,5]. It has been shown that two distinct pathways of pregranulosa cell (pGC) formation support the two different waves of ovarian follicles [8]. The granulosa cells (GCs) of the FWFs express genes distinct from those of the SWFs[7].

Estrogen receptor β (ERβ) is the predominant transcriptional regulator in ovarian GCs, and the expression of ERβ is also very high in the ovaries compared to other tissues[9]. Studies have demonstrated that the loss of ERβ results in ovulation failure in mice and rats[10-12]. Previous studies have identified that ERβ regulates an important group of genes essential for preovulatory follicle maturation [11-13]. ERβ is essential for preovulatory follicle maturation from the antral stage, and ovaries of Erβ^KO^ mice or rats contained fewer large antral follicles [11,13,14]. The loss of ERβ reduced the expression of Cyp19a1, Lhcgr, and Ptgs2 and increased the expression of Ar in antral follicles [14,15]. In addition to the steroidogenic genes, the loss of ERβ signaling also affects oocyte maturation[16].

Although numerous studies have reported the role of ERβ in preovulatory follicle maturation [11,13,14], several studies have shown the role of ERβ in oocyte nest breakdown[17-19]. Activation of ERβ in neonatal mice with diethylstilbestrol (DES) and other agonists resulted in polyovular follicles in mice ovaries [20-22]. We observed an increased primordial follicle activation (PFA) due to the loss of ERβ in the mutant rats, which was evident as early as PD 4.5 [7,23]. A loss-of-function mutation in the ERβ gene in humans was also found to cause complete ovarian failure [24]. However, only a few studies have analyzed follicular genes that were altered due to the loss of ERβ, particularly during early ovarian development [7,15,21].

This study utilized wildtype and Erβ^KO^ ovaries collected on PD 4.5, 6.5, and 8.5, the crucial time points of oocyte nest breakdown, primordial follicle formation, and PFA. Total RNAs were purified, and transcriptome analyses were performed using RNA sequencing. We observed remarkable differential expressions in epigenetic regulators, transcription factors, and downstream genes. We analyzed each gene’s differentially expressed transcript variants to detect the actual mRNAs that undergo up or downregulation[25]. The results indicate that loss of ERβ dysregulates the epigenetic regulators and transcription factors in Erβ^KO^ ovaries, disrupting the downstream genes and increasing ovarian follicle activation.

## 2. Results

### 2.1. Changes in Epigenetic Regulators

All epigenetic regulators, transcription factors, and downstream genes are analyzed based on the TPM (transcript per million) values of transcript variants[25]. Compared to PD 4.5 wildtype ovaries, PD 6.5 wildtype ovaries showed upregulation of 8 out of 17 differentially expressed epigenetic regulators (DEERs) (≥2-absolute fold changes and FDR p≤0.05; TPM values ≥ 5) (**Figure 1A**). The upregulated DEERs included histone readers Phf14-208 and Morf4l1-201, histone writer Msl1-201, histone chaperone Chrac1-201 and histone erasers Phf2-201, and Mysm1-201 (**Supplementary Table A1-1**). In contrast, the downregulated DEERs included histone reader Glyr1-202, histone writers Setd2-201, Msl1-202, Kat7-204, and Ogt-203, and histone erasers Phf8-202 and Hdac6-203 (**Supplementary Table A1-1**). While chromatin remodeler Actb-204 was upregulated, Bptf-202 and Chd6-202 were downregulated in PD6.5 wildtype ovaries (**Supplementary Table A1-1**).

**Figure 1.**
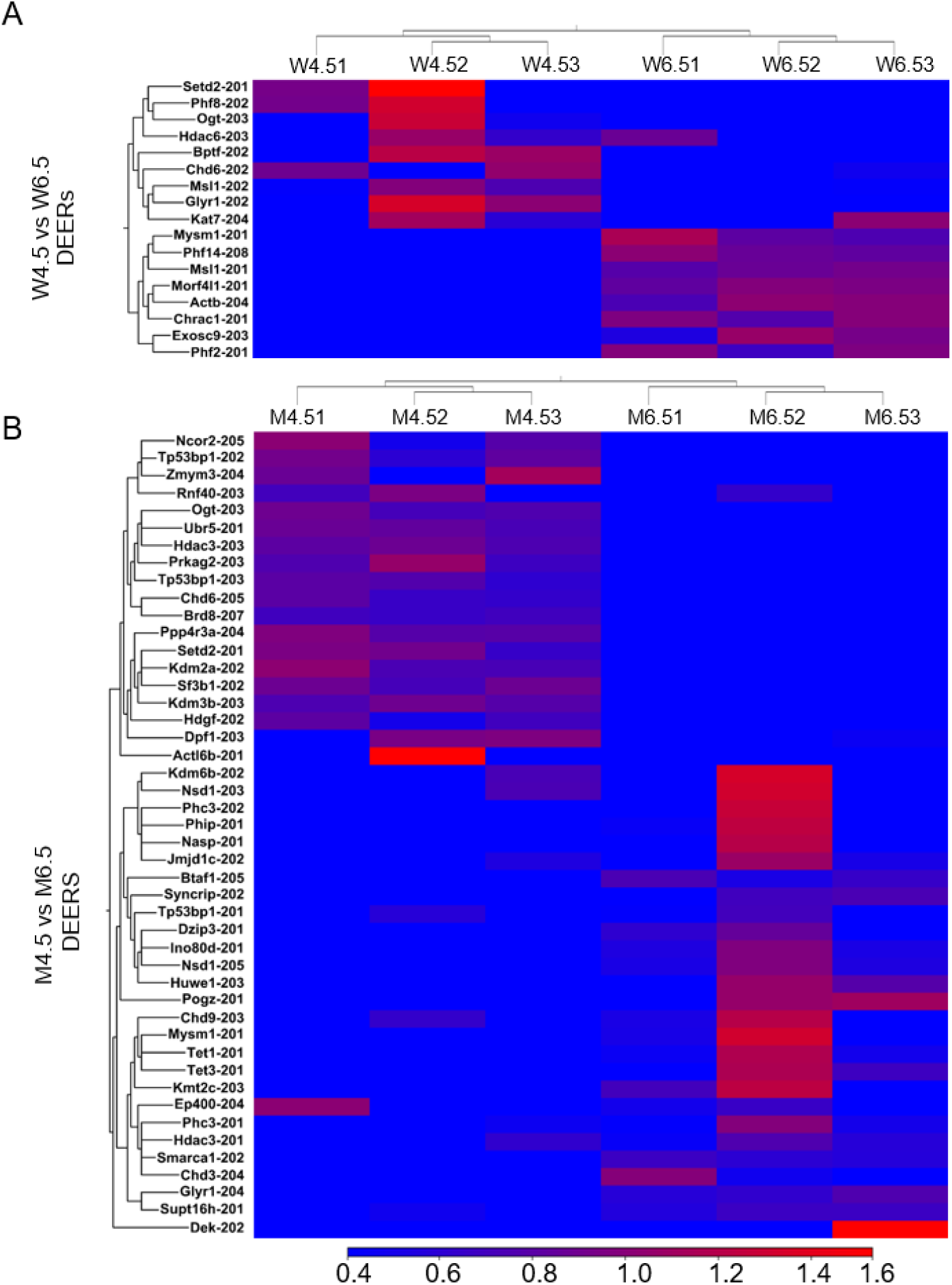
Heat Maps showing differentially expressed epigenetic regulators. RNA-Seq analysis was performed for postnatal days (PD) 4.5 and 6.5 wildtype and Erβ^KO^ rat ovaries. **A**) Heat Map shows differentially expressed epigenetic regulators (DEERs) (≥2-absolute fold changes; TPM values ≥ 5; FDR p≤0.05) between PD 4.5 and 6.5 wildtype ovaries. **B**) Heat Map shows DEERs (≥2-absolute fold changes and FDR p≤0.05; TPM values ≥ 5) between PD 4.5 and 6.5 of the Erβ^KO^ group ovaries. The three left columns represent PD 4.5 data, and the three right columns represent PD 6.5 data. All DEERs represent the relative TPM values of the transcript variants.

However, compared to PD 4.5 Erβ^KO^ ovaries, PD 6.5 Erβ_KO_ ovaries displayed the upregulation of 27 out of 46 DEERs (≥2-absolute fold changes and FDR p≤0.05; TPM values ≥ 5) (**Figure 1B**). The upregulated DEERs included histone readers Supt16h-201, Pogz-201 and Glyr1-204, histone writers Kmt2c-203, Huwe1-203, Dzip3-201 and Nsd1-205, and histone erasers Hdac3-201, Jmjd1c-202 and Mysm1-201 (**Supplementary Table A1-3**). In contrast, the downregulated DEERs included histone readers Tp53bp1-202, Tp53bp1-203, and Brd8-207, histone writers Setd2-201, Rnf40-203, Prkag2-203, and Ogt-203, and histone erasers Kdm2a-202, Kdm3b-203, Ncor2-205, Ppp4r3a-204, Hdac3-203 and Zmym3-204 (**Supplementary Table A1-3**). Chromatin remodelers Smarca1-202, Btaf1-205, and Ino80d-201 were upregulated in PD 6.5 ovaries, and Dpf1-203 and Ubr5-201 were downregulated (**Supplementary Table A1-3**).

We also detected that compared to PD 6.5 wildtype ovaries, PD 8.5 wildtype ovaries exhibited upregulation of 11 out of 24 DEERs (≥2-absolute fold changes and FDR p≤0.05; TPM values ≥ 5) (**Figure 2A**). The upregulated DEERs included histone readers Glyr1-202 and Cbx6-201, histone writer Kat7-204, and histone erasers Hdac6-203 and Kdm6a-205. The downregulated DEERs included histone readers, Morf4l1-201, and Phf14-208, histone writers Atxn7-201, Setdb1-203, Ube2b-202, and Ogt-202, histone eraser Morf4l2-207 (**Supplementary Table A1-2**). We also identified that chromatin remodelers Sfpq-201, Chd3-204, and Chd9-202 were upregulated, whereas Actb-204, Actl6a-202, and Hdgf-202 were downregulated in PD8.5 wildtype ovaries (**Supplementary Table A1-2**).

**Figure 2.**
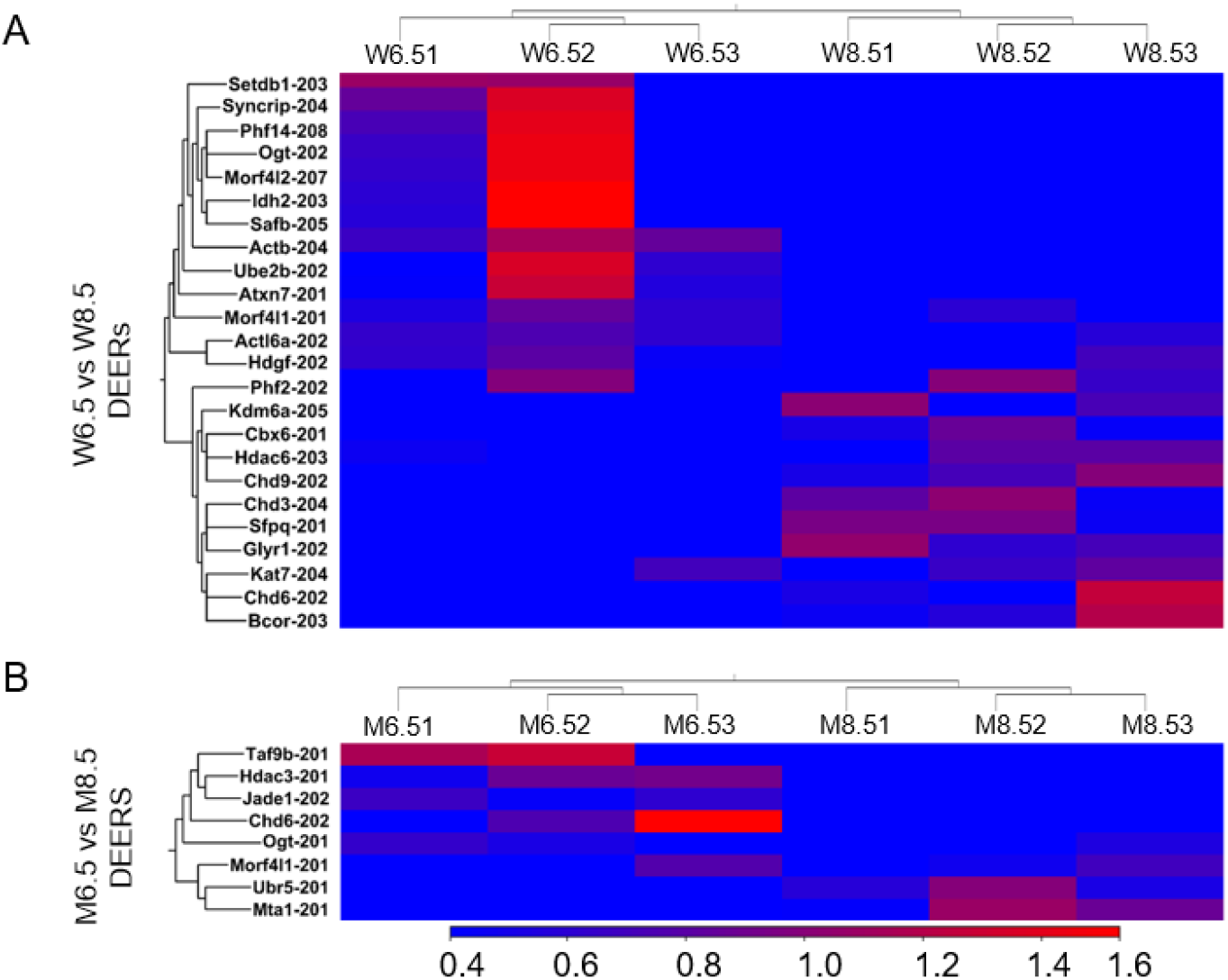
Heat Maps showing differentially expressed epigenetic regulators in wildtype and Erβ^KO^ ovaries on PD 6.5. RNA-Seq analysis was performed for postnatal days (PD) 6.5 and 8.5 wildtype and Erβ^KO^ rat ovaries. **A**) Heat Map shows differentially expressed epigenetic regulators (DEERs) (≥2-absolute fold changes; TPM values ≥ 5; FDR p≤0.05) between PD 6.5 and 8.5 wildtype ovaries. **B**) Heat Map shows DEERs (≥2-absolute fold changes and FDR p≤0.05; TPM values ≥ 5) between PD 6.5 and 8.5 of the Erβ^KO^ group ovaries. The three left columns represent PD 6.5 data, and the three right columns represent PD 8.5 data. All DEERs represent the relative TPM values of the transcript variants.

In contrast, compared to PD 6.5 Erβ^KO^ ovaries, PD 8.5 Erβ^KO^ ovaries showed upregulation of 3 out of only 8 DEERs (≥2-absolute fold changes and FDR p≤0.05; TPM values ≥ 5) (**Figure 2B**). Those included histone reader Morf4l1-201, chromatin remodelers Ubr5-201 and Mta1-201. The downregulated DEERs included histone writer Jade1-202, histone chaperone Taf9b-201, histone eraser Hdac3-20, and chromatin remodeler Chd6-202 (**Supplementary Table A1-4**).

### 2.2. Changes in Transcription Factors

We detected that 14 out of 23 differentially expressed transcription factors (DETFs) were upregulated in PD 6.5 wildtype ovaries compared to PD 4.5 ovaries (≥2-absolute fold changes and FDR p≤0.05; TPM values ≥ 5) (**Figure 3A**). The top 5 upregulated DETFs included Zfp518a-204, Foxk-203, Zfp711-202, Chchd3-204 and Zbed4-201 and the top 5 downregulated DETFs included Mlx-204, Kat7-204, Tgif1-202, Prr12-202 and Rbck1-203 (**Supplementary Table A2-1**).

**Figure 3.**
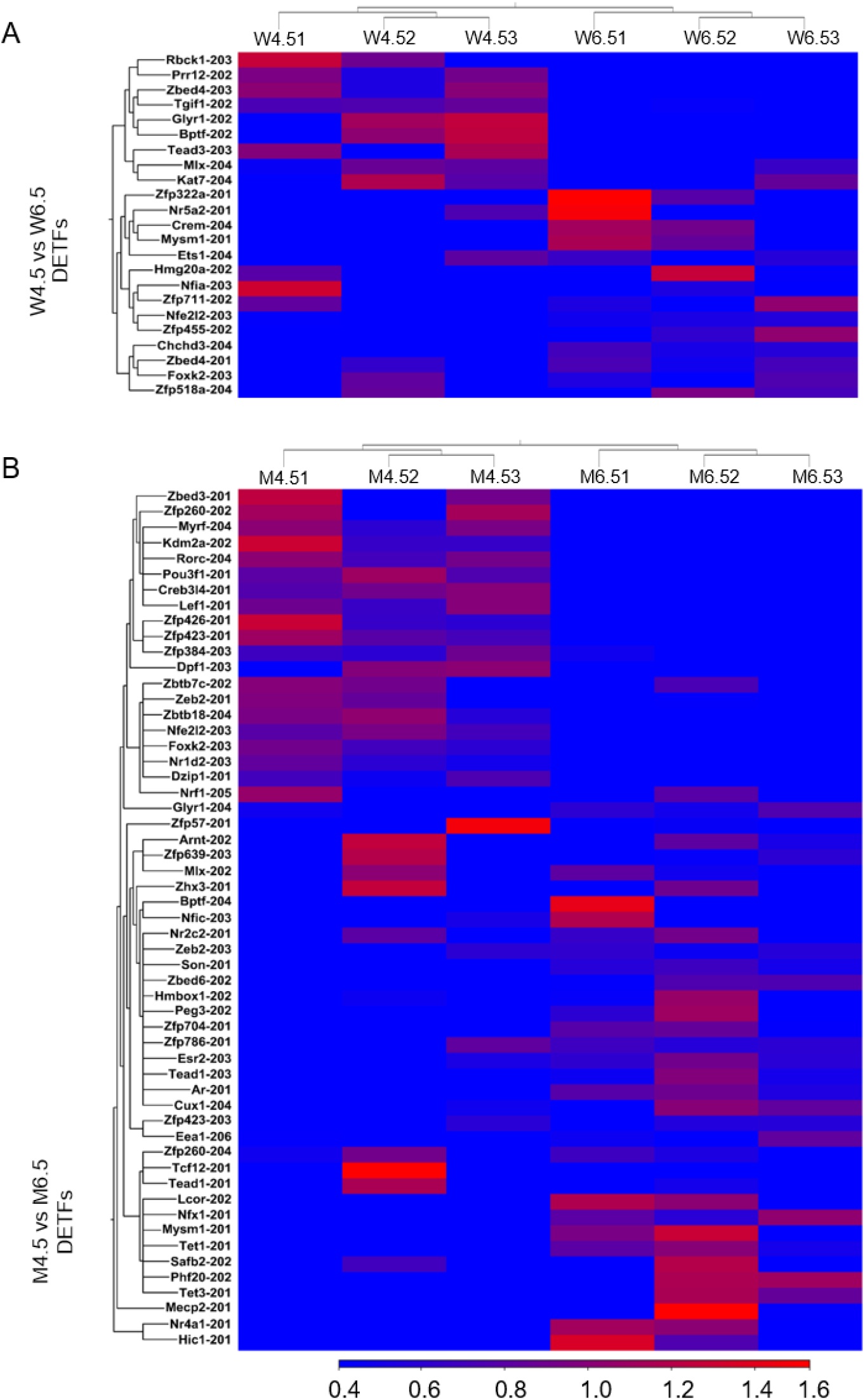
Heat Maps showing differentially expressed transcription factors on PD 6.5. RNA-Seq analysis was performed for postnatal days (PD) 4.5 and 6.5 wildtype and Erβ^KO^ rat ovaries. **A**) Heat Map shows differentially expressed transcription factors (DETFs) (≥2-absolute fold changes; TPM values ≥ 5; FDR p≤0.05) between PD 4.5 and 6.5 wildtype ovaries. **B**) Heat Map shows DETFs (≥2-absolute fold changes and FDR p≤0.05; TPM values ≥ 5) between PD 4.5 and 6.5 of the Erβ^KO^ group ovaries. The three left columns represent PD 4.5 data, and the three right columns represent PD 6.5 data. All DETFs represent the relative TPM values of the transcript variants.

However, compared to PD 4.5 Erβ^KO^ ovaries, PD 6.5 Erβ^KO^ ovaries showed upregulation of 36 out of 55 DETFs (≥2-absolute fold changes and FDR p≤0.05; TPM values ≥ 5) (**Figure 3B**). The top 5 upregulated DETFs included Zfp786-201, Mlx-202, Nfx1-201, Glyr1-204 and son-201 and the top 5 downregulated DETFs included Zfp384-203, Zfp260-202, Zbtb7c-202, Dpf1-203, and Nr1d2-203 (**Supplementary Table A2-3**).

We also identified that compared to PD 6.5 wildtype ovaries, 32 out of 68 DETFs (≥2-aboslute fold changes and FDR p≤0.05; TPM values ≥ 5) were upregulated in PD 8.5 wildtype ovaries (**Figure 4A**). The top 5 DETFs included Mlx-204, Kat7-204, Fosl2-201, Pparg-202, and Foxo1-201 and the top 5 downregulated DETFs included Crem-210, Nfe2l2-203, Ets1-204, Lhx9-203, and Lhx9-201 (**Supplementary Table A2-2**).

**Figure 4.**
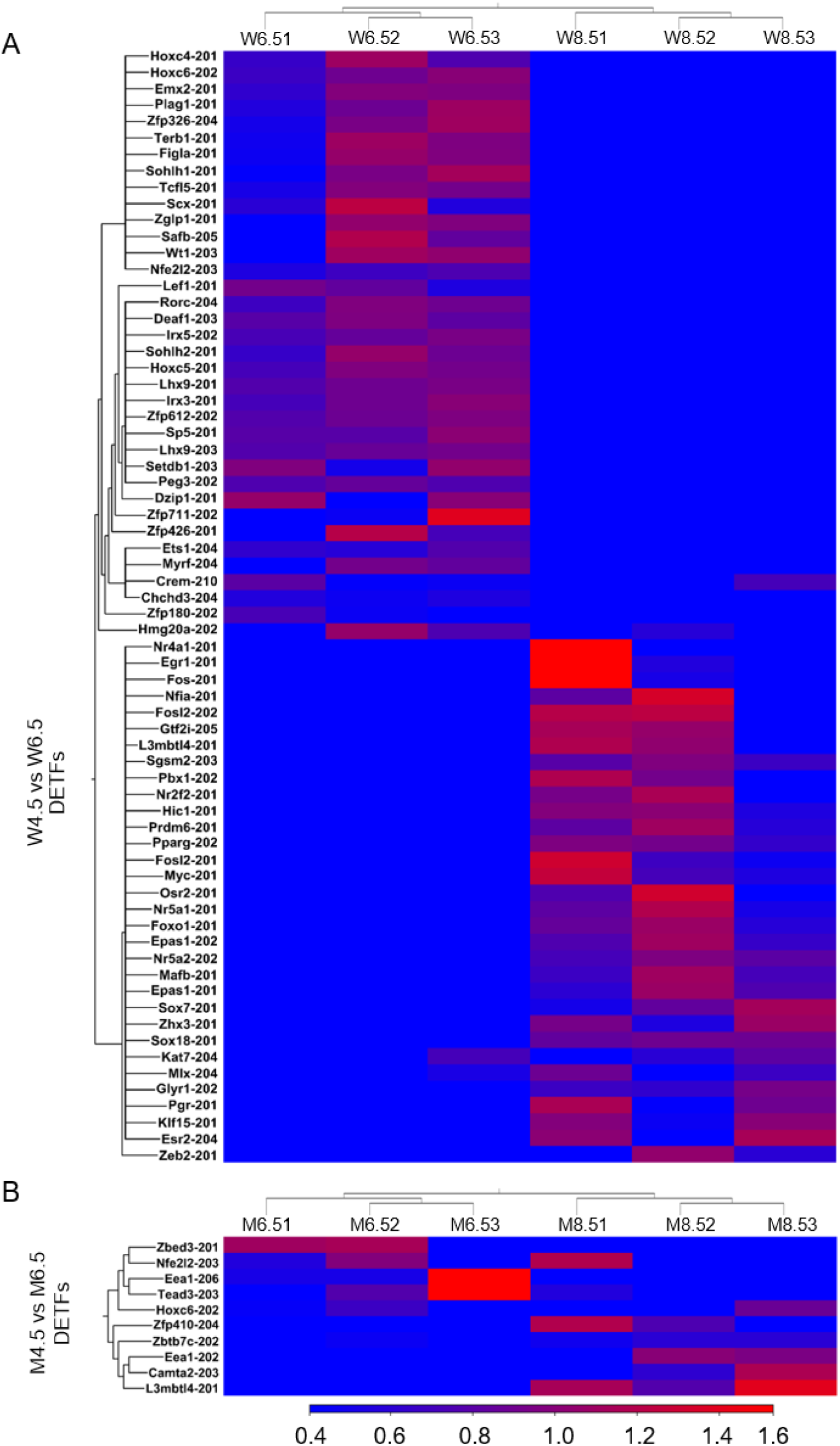
Heat Maps showing differentially expressed transcription factors. RNA-Seq analysis was performed for postnatal days (PD) 6.5 and 8.5 wildtype and Erβ^KO^ rat ovaries. **A**) Heat Map shows differentially expressed transcription factors (DETFs) (≥2-absolute fold changes; TPM values ≥ 5; FDR p≤0.05) between PD 6.5 and 8.5 wildtype ovaries. **B**) Heat Map shows DETFs (≥2-absolute fold changes and FDR p≤0.05; TPM values ≥ 5) between PD 6.5 and 8.5 of the Erβ^KO^ group ovaries. The three left columns represent PD 6.5 data, and the three right columns represent PD 8.5 data. All DETFs represent the relative TPM values of the transcript variants.

In contrast, compared to PD 6.5 Erβ^KO^ ovaries, PD 8.5 Erβ^KO^ ovaries showed upregulation of 5 out of 10 DETFs (≥2-absolute fold changes and FDR p≤0.05; TPM values ≥ 5) (**Figure 4B**). The upregulated DETFs included Zbtb7c-202, Zfp410-204, Eea1-202, Camta2-203 and L3mbtl4-201 and the downregulated DETFs were Nfe2l2-203, Zbed3-201, Tead3-203, Hoxc6-202 and Eea1-206 (**Supplementary Table A2-4**).

### 2.3. Changes in Downstream Genes

Changes in DEERs and DEFTs in PD 6.5 wildtype ovaries compared to PD 4.5 wildtype ovaries upregulated 368 out of 581 differentially expressed downstream genes (DEDGs) (**Figure 5A**). Similarly, the changes in DEERs and DETFs in PD 6.5 Erβ^KO^ ovaries compared to PD 4.5 Erβ^KO^ ovaries ovaries upregulated 419 out of 744 DEDGs (**Figure 5B**).

**Figure 5.**
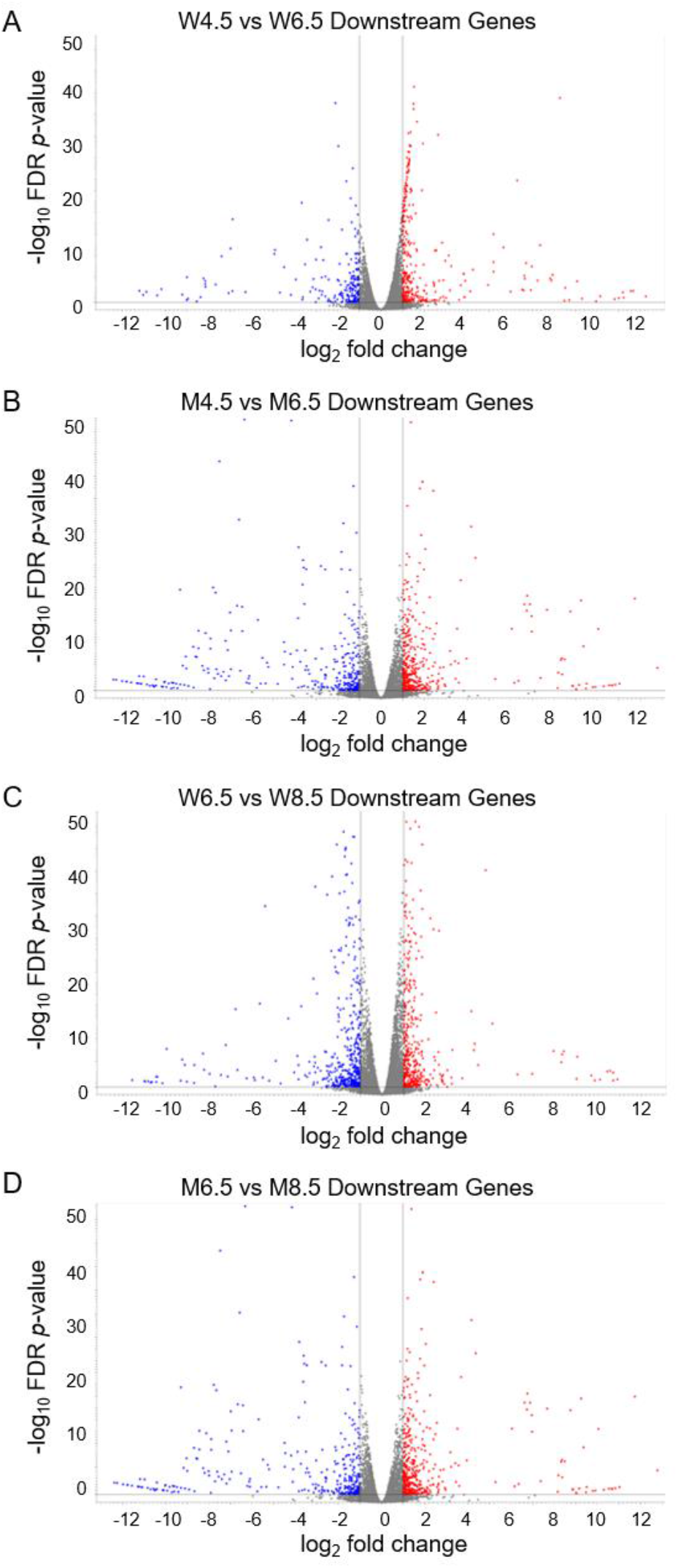
Volcano Plots display differentially expressed downstream genes. Volcano Plots show the differentially expressed downstream genes (DEDGs) between wildtype postnatal day (PD) 4.5 and 6.5 rat ovaries (**A**), between Erβ^KO^ PD 4.5 and 6.5 ovaries (**B**), between wildtype PD 6.5 and 8.5 ovaries (**C**), and between Erβ^KO^ PD 6.5 and 8.5 ovaries (**D**). All DEDGs (≥2-absolute fold changes; TPM values ≥ 5; and FDR p≤0.05) represent the relative TPM values of the transcript variants. While the red dots denote upregulation, the blue denotes downregulation of the DEDGs.

Changes in DEERs and DETFs in PD 8.5 wildtype ovaries compared to PD 6.5 wildtype ovaries also upregulated 424 of 920 DEDGs (**Figure 5C**). In contrast, PD 8.5 Erβ^KO^ ovaries had significantly fewer DEERs (24 vs. 8) and fewer DETFs (68 vs. 10) compared to wildtype ovaries, which were associated with markedly fewer number of DEDGs (920 vs. 191) (**Figure 5D**). In PD 8.5 Erβ^KO^ ovaries, only 111 of 191 DEDGs were upregulated (**Figure 5D**).

In addition to the quantitative differences in DEERs and DETFs between the wildtype and Erβ^KO^ ovaries, we detected distinct differences in the identities of the regulators of gene expression (Figure 6). Only 3 out of 46 DEERs were common between wildtype and Erβ^KO^ ovaries on PD 6.5, and 2 out of 8 DEERs were common on PD 8.5 (**Figure 6 A, 6B**). Similarly, only 3 out of 55 DETFs were common between wildtype and Erβ^KO^ ovaries on PD 6.5, and 3 out of 10 DETFs were common on PD 8.5 (**Figure 6C, 6D**). The DEDGs also showed a similar pattern; only 99 out of 744 DEDGs were common between wildtype and Erβ^KO^ ovaries on PD 6.5, and only 61 out of 191 DEDGs were common on PD 8.5 (**Figure 6E, 6F**).

**Figure 6.**
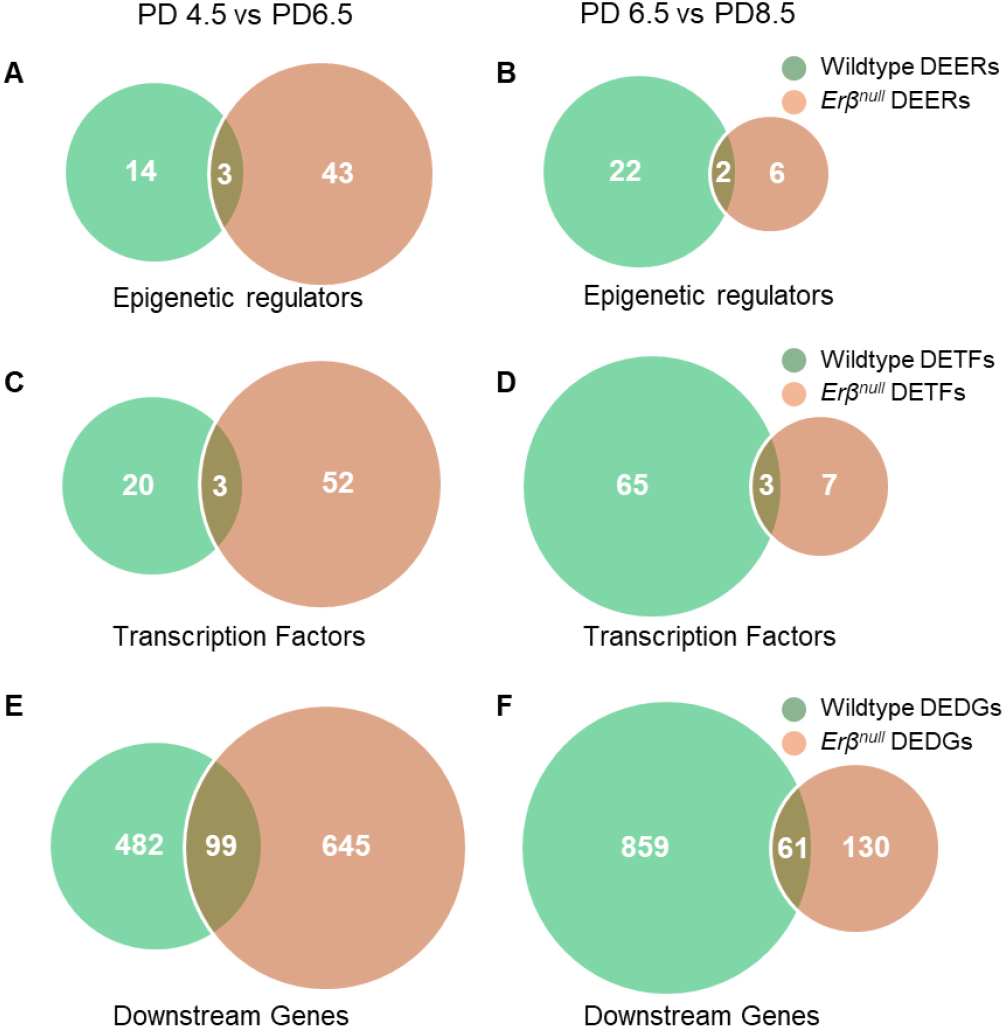
A comparison of differentially expressed epigenetic regulators (DEERs), transcription factors (DETFs), and downstream genes (DEDGs). Differential expression of DEERs, DETFs, and DEDGs (≥ 2-absolute fold change, TPM ≥5, and FDR p-value ≤0.05) in postnatal days (PD) 4.5 and 6.5 between wildtype and Erβ^KO^ rat ovaries are shown in the left panels (**A, C, E**). Differential expression of DEERs, DETFs, and DEDGs (absolute fold change ≥2, TPM ≥5, and FDR p-value≤0.05) in PD 6.5 and 8.5 between wildtype and Erβ^KO^ rat ovaries are shown in the right panels (**B, D, F**). All DEERs, DETFs, and DEDGs represent the relative TPM values of the transcript variants.

### 2.4 Changes in Genes Related to Follicle Assembly

Oocyte nest breakdown results in primordial follicle assembly during early ovarian development[2]. Studies have identified numerous transcription factors and downstream genes involved in this process[2,26,27]. We have analyzed the 40 transcript variants of 27 known genes to see if they exhibited any differential expression. In PD 6.5 wildtype ovaries, we detected upregulation of Irx3-201 and downregulation of Nobox-201 and Akt1-205. In contrast, Inhba-201, Elavl2-201, Elavl2-202, Notch2-201, and Akt1-204 were upregulated, and Notch2-204, Irx3-201, Irx5-202, Elavl2-203, Nobox-201, Figla-201, Sohlh1-201, were downregulated in PD 6.5 Erβ^KO^ovaries. In PD 8.5 wildtype ovaries, Inhba-201 and Ak1-204 were upregulated and Irx3-201, Irx5-202, Elavl2-203, elavl2-204, Nobox-201 Figla-201, and Sohlh1-201 were downregulated. In contrast, Inhba-201 was upregulated, and Elavl2-202, Elavl2-204, Figla-201, Irx3-201, and Sohlh1-201 were downregulated in PD 8.5 Erβ^KO^ovaries.

### 2.5. Changes in Genes Related to Primordial Follicle Activation

PFA is a complex process that activates the dormant primordial follicles into primary follicles[28]. Once activated, the follicles progress through successive developmental processes or undergo atresia[29,30]. We analyzed the 58 transcript variants of 34 genes known to be involved in PFA. In PD 6.5 wildtype ovaries, we detected upregulation of Nr5a2-201, Igf1-201, and Sirt1-204 and downregulation of Hdac6-203, Tsc2-201, Tsc2-203, Nobox-201 and Cdh1-201. In PD 6.5 Erβ^KO^ ovaries, Igf1-201, Igf1-204, Nr5a2-201, Nr5a2-202, and Foxl2-201 were upregulated, and Tsc2-201, Tsc2-202, Sohlh1-201, Sohlh2-201, Nobox-201, and Hdac6-202 were downregulated. In PD 8.5 wildtype ovaries, Nr5a1-201, Nr5a2-202, Hdac6-203, Esr2-204, Bmp15-201 and Tgfb1-201 were upregulated and Sohlh1-201, Sohlh2-201, Nobox-201, and Sirt1-204 were downregulated. Contrarily, Igf1-203 was upregulated, and Sholh2-202, Smad3-202, Tsc2-202, Nr5a2-201, and Sirt1-203 were downregulated in PD 8.5 Erβ^KO^ ovaries.

### 2.6. Changes in Genes Related to Steroidogenesis

Granulosa cells play a crucial role in steroidogenesis[31]. We analyzed the 36 transcript variants of 20 genes known to be involved in steroidogenesis. In PD 6.5 wildtype ovaries, we detected upregulation of Hsd17b2-201, Hsd3b3-201, Cyp11a1-201, Cyp11a1-202, Cyp17a1-201 Cyp19a1-201, and Star-201 and downregulation of Hsd11b2-201, Hsd17b7-201, Hsd17b4-203 and Hsd3b7-202. In contrast, Hsd3b3-201, Hsd17b1-201, Cyp17a1-201, Cyp19a1-201, Cyp11a1-201, Cyp11a1-202, and Star-201 were upregulated, and Hsd3b7-201 and Hsd11b2-201 were downregulated in PD 6.5 Erβ^KO^ ovaries. In PD 8.5 wildtype ovaries, Hsd3b3-201, Hsd17b1-201, Cyp11a1-201, Cyp11a1-202, Cyp17a1-201 and Cyp19a1-201 were upregulated and Hsd17b2-201 was downregulated. In contrast, Hsd3b3-201, Hsd3b7-201, Hsd17b1-201, Cyp11a1-201, Cyp11a1-202, and Cyp17a1-201 were upregulated, and Hsd3b2-201 and Hsd17b8-202 were downregulated in PD 8.5 Erβ^KO^ ovaries.

## 3. Discussion

We previously detected that ERβ plays a gatekeeping role in regulating PFA[23], and the loss of ERβ accelerates PFA in both first wave follicles as well as second-wave follicles[7]. Increased PFA of the first-wave primordial follicles in Erβ^KO^ ovaries becomes evident as early as PD 4.5, while increased PFA of SW primordial follicles becomes apparent as early as PD 8.5 [7,23]. Accordingly, in this study, we collected total RNAs from PD 4.5, PD 6.5, and PD 8.5 wildtype ovaries and compared the transcriptomes with those of Erβ^KO^ ovaries at corresponding time points. Our results identified differential expression of novel transcripts during early ovarian development and elucidated the epigenetic and transcriptional basis of the Erβ^KO^ phenotype in postnatal ovaries.

Rodent ovarian follicles develop during the perinatal and early postnatal period[32]. Oocyte nest breakdown and formation of primordial follicles occur in two successive waves [33,34]. The first-wave primordial follicles are formed in the ovarian medulla and are rapidly activated [35]. However, the second-wave primordial follicles start assembling in the ovarian cortex at PD4.5 and are completed by PD8.5. In contrast to the first-wave primordial follicles, the second-wave primordial follicles remain dormant until recruited for PFA[36,37]. Wildtype PD4.5 follicles contained activated first-wave follicles, primordial follicles of first-wave, and oocyte nests; however, the second-wave follicles remained as oocyte nests (**Figure 7A, 7D**). Activation of first-wave primordial follicles increased in PD 6.5 and 8.5 wildtype ovaries (**Figure 7B, 7C**), which was quantitatively more in Erβ^KO^ ovaries [7,23] (**Figure 7E, 7F**). While PD 8.5 ovaries contained primordial follicles of second-wave (**Figure 7C**), a portion of the second-wave primordial follicles became activated in Erβ^KO^ ovaries [7,23] (**Figure 7F**). As we have performed the transcriptomic analyses on whole ovaries, the results represented gene expression in both the first and the second wave follicles.

**Figure 7.**
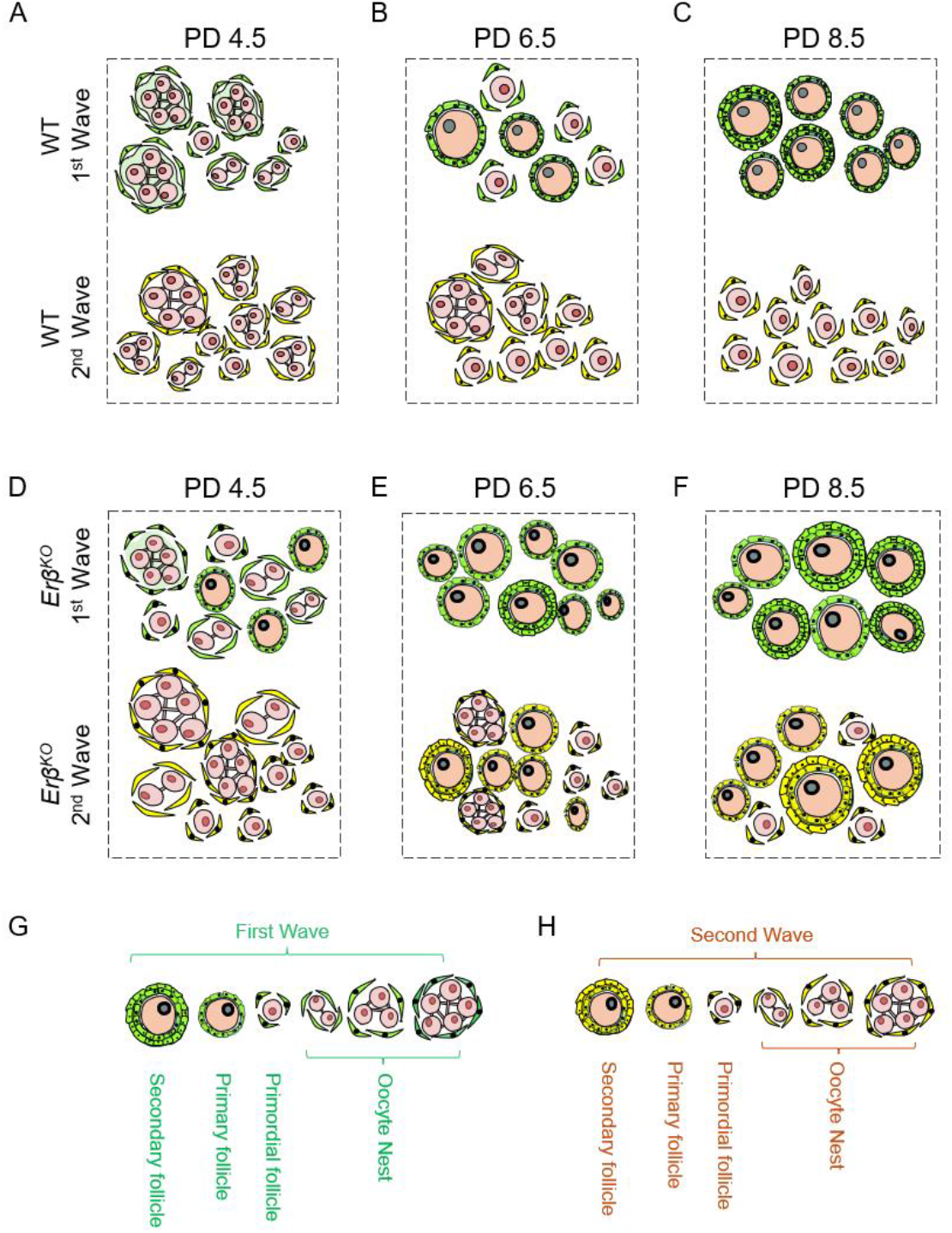
Ovarian follicles in wildtype and Erβ^KO^ rat ovaries during the postnatal period. Wildtype (WT) PD4.5 follicles contained activated first-wave (1^st^ wave) follicles, primordial follicles of first-wave, and oocyte nests of both waves. However, at this time point, second-wave (2^nd^ wave) follicles remained as oocyte nests. Activation of 1^st^ wave primordial follicles increased in PD 6.5 and 8.5 wildtype ovaries, which was quantitatively more in Erβ^KO^ ovaries. While PD 8.5 ovaries contained primordial follicles of 2^nd^ wave, a portion of the primordial follicles became activated in Erβ^KO^ ovaries.

The transcriptome analyses in this study represent the transcript variants of the genes expressed in ovarian follicles and interstitial cells. The follicular cells represented many GCs, followed by theca cells and oocytes. We analyzed the changes in epigenetic regulators and transcription factors during early ovarian development: in PD 6.5 ovaries compared to PD 4.5 ovaries and in PD 8.5 ovaries compared to PD 6.5 ovaries. We performed similar analyses in wildtype ovaries as well as in Erβ^KO^ ovaries and compared the findings between wildtype and Erβ^KO^ ovaries. We further analyzed the DEERs, DETFs, and DEDGs to identify the critical transcripts according to their cellular origin and functional involvement in follicle development and/or steroidogenesis.

We observed that the number of DEERs and DETFs proportionately correlated with the number of DEDGs in PD6.5 and PD8.5 in either wildtype or Erβ^KO^ ovaries. Compared to PD4.5 wildtype ovaries, PD 6.5 wildtype ovaries showed 17 DEERs and 23 DETFs, resulting in 581 DEDGs. In age-matched PD6.5 Erβ^KO^ ovaries, we detected 46 DEERs and 55 DETFs, resulting in 744 DEDGs. Similarly, compared to PD6.5 wildtype ovaries, PD8.5 wildtype ovaries showed 24 DEERs and 68 DETFs, resulting in 920 DEDGs. Remarkably, PD8.5 Erβ^KO^ ovaries showed only 8 DEERs and 10 DETFs, resulting in only 191 DEDGs. These observations clearly indicate that the DEERs and DETFs determine the numbers of DEDGs during early ovarian development. Our observations also suggest that loss of ERβ dysregulates epigenetic regulars and transcription factors as early as PD 4.5, which became more prominent during the following days.

In addition to the difference in numbers of DEERs, DETFs, and DEDGs during ovarian development between wildtype and Erβ^KO^ ovaries, the identities of the transcripts were also vastly diverged. Only 7% of DEERs, 5% of DETFs, and 13% of DEDGs were common to PD 6.5 wildtype and Erβ^KO^ ovaries. Although the proportion of the shared genes was increased in PD 8.5 ovaries, those were still 25% of DEERs, 30% of DETFs, and 315 of DEDGs between wildtype and Erβ^KO^ ovaries. These results suggest that loss of ERβ in mutant rat ovaries impacts the development of follicular cells from a very early stage. While changes in known epigenetic regulators transcription factors and downstream other vital genes were observed in wildtype ovaries as expected, those were not present in Erβ^KO^ ovaries. These findings indicate that loss of ERβ results in the regulation of gene expression from a very early stage. While ERβ is predominantly expressed in granulosa cells, the loss of ERβ is expected to impact the proliferation and differentiation of granulosa cells directly [38,39]. However, bidirectional signaling between granulosa cells and theca cells and between granulosa cells and oocytes is well-known in ovarian biology[40,41]. Therefore, it may also be expected that loss of ERβ in granulosa cells may indirectly impact gene regulation in theca cells and oocytes[42,43].

## 4. Materials and Methods

### 4.1. Experimental Model

Wildtype and Erβ^KO^ Holtzman Sprague-Dawley (HSD) female rats were included in this study. The Erβ^KO^ rat model was generated by targeted deletion of exon 3 in the Erβ gene, causing a frameshift and null mutation[44]. Rats were screened for the presence of mutations by genotyping PCR using tail-tip or toe-clip DNA samples as described previously[44]. All procedures were performed according to the protocol approved by the University of Kansas Medical Center Animal Care and Use Committee.

### 4.2. RNA Sequencing

Ovaries were collected from 4.5, 6.5, and 8.5 days-old wildtype and Erβ^KO^ female rats. Ovaries were snap-frozen in liquid nitrogen, preserved at -80°C, and later used for RNA purification [7,23,45]. Total RNAs were extracted from the ovaries using TRI Reagent (Millipore-Sigma, St. Louis, MO), and gene expressions were evaluated using RNA sequencing (RNA-Seq) analyses. RNA samples with a RIN value ≥ 9 were used for library preparation. 500 ng of total RNA from each sample was used for the RNA-Seq library preparation using the TruSeq Stranded mRNA kit (Illumina, San Diego, CA) following the manufacturer’s instructions[7]. The RNA-Seq libraries were evaluated for quality at the KUMC Genomics Core and sequenced on an Illumina HiSeq X sequencer at Novogene Corporation (Sacramento, CA). The RNA-Seq data have been submitted to the Sequencing Read Archive (PRJNA576013).

### 4.3. Analysis of RNA Sequencing Data

All RNA-Seq data were analyzed using CLC Genomics Workbench (Qiagen Bioinformatics) as described in our previous publications[11,12]. Clean reads were obtained by removing low-quality reads and trimming the adapter sequences. The high-quality reads were aligned to the Rattus norvegicus reference genome (GRCm39), gene (GRCm39.111_Gene), and mRNA sequences (GRCm39.111_mRNA) using the default parameters[25]. The expression values were measured in TPM, and the values of individual transcript variants (TE) in whole rat ovaries were determined as described in our previous publications[11,12]. We have recently demonstrated the relative advantage of analyzing transcript variants over gene expression (GE) values[25].

### 4.4. Analysis of the Transcript Variants

We have analyzed three categories of differentially expressed transcript variants into three categories: epigenetic regulators, transcription factors, and all the transcript variants indicated as downstream genes. A list of 690 epigenetic regulators was prepared according to a recent publication[46]. The list of 1,374 mouse transcription factors that were curated by the Gifford lab (https://cgs.csail.mit.edu/ReprogrammingRecovery/mouse_tf_list.html) according to the human TFs[47]. Each RNA-Seq data file containing TE values was used to generate new tracks containing only the epigenetic regulators or transcription factors, which were used in subsequent analyses.

This study focused on the moderately expressed transcripts with TPM values ≥ 5. The threshold *p*-values were selected according to the false discovery rate (FDR) to identify the differentially expressed transcript variants. A transcript variant was considered differentially expressed if the absolute fold change was ≥ 2 and the FDR *p*-value was ≤ 0.05 [11,12,48]. The differentially expressed transcript variants were divided into 3 groups: up-regulated (≥2-absolute fold changes and FDR *p* ≤0.05), downregulated (≤ -2-fold changes and FDR p ≤0.05), and insignificant (either ≤ absolute 2-fold changes and/or FDR *p* ≥ 0.05).

Differential expressions of the transcript variants were determined in two different ways. In the first approach, the TE values of PD 6.5 ovaries were compared to that of PD 4.5 ovaries, and then TE values of PD 8.5 ovaries were compared to that of PD 6.5 ovaries within the wildtype or the Erβ^KO^ group. In the second approach, the differentially expressed transcript variants (DEERs, DETFs, and DEDGs) in PD 6.5 or PD 8.5 wildtype ovaries were compared to those of Erβ^KO^ ovaries.

### 4.5. Statistical analysis

Each RNA-Seq library was prepared using pooled RNA samples from three individual wildtype or Erβ^KO^ rats. Each group of RNA sequencing data consisted of three to four different libraries. For RNA Seq, each study group contained three library samples. In CLC Genomics Workbench, the ‘differential expression for RNA-Seq tool’ performs multifactorial statistics on a set of expression tracks based on a negative binomial generalized linear model (GLM). The final GLM-fit and dispersion estimate calculates the total likelihood of the model given the data and the uncertainty of each fitted coefficient. Two statistical tests-the Wald and the Likelihood Ratio tests-use one of these values. The Across groups (ANOVA-like) comparison uses the Likelihood Ratio test.

## 5. Conclusions

We analyzed the whole ovary transcriptome during the postnatal period to identify the genes involved in early ovarian development. We emphasized the epigenetic regulators and transcription factors to understand the mechanism of downstream gene expression. Our observations indicate that loss of ERβ dysregulates the epigenetic regulators and transcription factors in Erβ^KO^ ovaries, which disrupts the downstream genes in ovarian follicles and increases follicle activation.

## Supplementary Materials

The following supporting information can be downloaded at: www.mdpi.com/xxx/s1, Figure S1: title; Table S1: title; Video S1: title.

Author Contributions

M.A.K.R. conceptualization, supervision, funding acquisition, resources, and writing; K.V. data curation, methodology, investigation, formal analysis, and original draft preparation; R.M., A.M, and Y.S. software and data validation; P.E. F. review and editing. All authors have read and agreed on the contents of the manuscript.

## Institutional Review Board Statement

This study was approved by the IACUC of KUMC (2021-2603).

## Informed Consent Statement

Not applicable

## Data Availability Statement

SRA, NCBI

Acknowledgments

We acknowledge the editorial board of IJMS’s willingness to waive the publication fees. We also acknowledge Qiagen Bioinformatics for their continued support.

## Disclaimer/Publisher’s Note

The statements, opinions, and data contained in all publications are solely those of the individual author(s) and contributor(s) and not of MDPI and/or the editor(s). MDPI and/or the editor(s) disclaim responsibility for any injury to people or property resulting from any ideas, methods, instructions, or products referred to in the content.

