## Supplemental Tables for "Epigenetic and transcriptional regulation of ovarian development altered in *Erβ*^KO^ ovaries"

### A.1. Epigenetic Regulators

**Supplementary Table A1-1.** Differentially expressed epigenetic regulators in PD 6.5 wildtype ovaries compared to PD 4.5 wildtype ovaries.

| Name | Chrom | Max group mean | Fold change | FDR p-value | ENSEMBL |
|---|---|---|---|---|---|
| Exosc9-203 | 2 | 10.71 | 3.41 | 0.01 | ENSRNOT00000108461 |
| Chd6-202 | 3 | 10.67 | -5.36 | 6.56E-06 | ENSRNOT00000089958 |
| Phf14-208 | 4 | 5.77 | 2.01 | 1.15E-04 | ENSRNOT00000108109 |
| Mysm1-201 | 5 | 13.68 | 2.37 | 1.63E-03 | ENSRNOT00000039554 |
| Chrac1-201 | 7 | 51.61 | 2.09 | 4.11E-08 | ENSRNOT00000012121 |
| Morf4l1-201 | 8 | 30.77 | 2,917.98 | 3.98E-04 | ENSRNOT00000080421 |
| Setd2-201 | 8 | 6.07 | -148.22 | 4.90E-07 | ENSRNOT00000028409 |
| Glyr1-202 | 10 | 26.89 | -2.6 | 2.87E-06 | ENSRNOT00000095486 |
| Kat7-204 | 10 | 6.24 | -278.56 | 4.74E-05 | ENSRNOT00000109616 |
| Msl1-201 | 10 | 12.52 | 3.12 | 1.69E-03 | ENSRNOT00000012812 |
| Msl1-202 | 10 | 6.37 | -128.82 | 5.32E-04 | ENSRNOT00000101887 |
| Bptf-202 | 10 | 5.38 | -3.11 | 4.29E-05 | ENSRNOT00000098328 |
| Actb-204 | 12 | 5.6 | 614.8 | 0.01 | ENSRNOT00000116486 |
| Phf2-201 | 17 | 5.13 | 47.81 | 1.35E-07 | ENSRNOT00000022669 |
| Hdac6-203 | X | 13.57 | -2.72 | 5.88E-03 | ENSRNOT00000116471 |
| Phf8-202 | X | 7.33 | -2.08 | 0.02 | ENSRNOT00000093422 |
| Ogt-203 | X | 44.21 | -5.92 | 1.26E-06 | ENSRNOT00000100805 |

**Supplementary Table A1-2.** Differentially expressed epigenetic regulators in PD 8.5 wildtype ovaries compared to PD 6.5 wildtype ovaries.

| Name | Chrom | Max group mean | Fold change | FDR p-value | ENSEMBL |
|---|---|---|---|---|---|
| Idh2-203 | 1 | 8.46 | -2.01 | 6.54E-03 | ENSRNOT00000109065 |
| Actl6a-202 | 2 | 17.24 | -2,068.55 | 4.27E-03 | ENSRNOT00000098294 |
| Hdgf-202 | 2 | 16.18 | -326.91 | 1.19E-06 | ENSRNOT00000065211 |
| Setdb1-203 | 2 | 8.44 | -2.04 | 1.50E-06 | ENSRNOT00000111803 |
| Chd6-202 | 3 | 6.71 | 3.4 | 1.77E-03 | ENSRNOT00000089958 |
| Phf14-208 | 4 | 5.77 | -2.62 | 7.36E-04 | ENSRNOT00000108109 |
| Sfpq-201 | 5 | 129.18 | 2.01 | 2.89E-29 | ENSRNOT00000084202 |
| Cbx6-201 | 7 | 20.8 | 2.01 | 4.29E-03 | ENSRNOT00000068033 |
| Syncrip-204 | 8 | 7.8 | -2.09 | 4.71E-07 | ENSRNOT00000095785 |
| Morf4l1-201 | 8 | 30.77 | -3,077.06 | 2.57E-03 | ENSRNOT00000080421 |
| Safb-205 | 9 | 12.48 | -2.03 | 2.71E-07 | ENSRNOT00000115324 |
| Glyr1-202 | 10 | 20.62 | 2.11 | 5.74E-05 | ENSRNOT00000095486 |
| Ube2b-202 | 10 | 31.35 | -2.14 | 3.06E-04 | ENSRNOT00000106084 |
| Chd3-204 | 10 | 9.2 | 2.66 | 5.32E-12 | ENSRNOT00000114886 |
| Kat7-204 | 10 | 6.46 | 322.85 | 1.19E-06 | ENSRNOT00000109616 |
| Actb-204 | 12 | 5.6 | -649.91 | 5.00E-03 | ENSRNOT00000116486 |
| Atxn7-201 | 15 | 14.75 | -2.8 | 3.20E-07 | ENSRNOT00000010103 |
| Phf2-202 | 17 | 20.15 | 2.05 | 4.17E-03 | ENSRNOT00000087692 |
| Chd9-202 | 19 | 5.79 | 2.28 | 1.32E-06 | ENSRNOT00000076138 |
| Kdm6a-205 | X | 8.03 | 3.02 | 6.15E-03 | ENSRNOT00000109873 |
| Bcor-203 | X | 5.33 | 2.83 | 0.01 | ENSRNOT00000116761 |
| Hdac6-203 | X | 13.11 | 2.84 | 7.72E-05 | ENSRNOT00000116471 |
| Ogt-202 | X | 200.37 | -2.23 | 2.18E-62 | ENSRNOT00000082967 |
| Morf4l2-207 | X | 5.74 | -2.06 | 7.96E-05 | ENSRNOT00000116026 |

**Supplementary Table A1-3.** Differentially expressed epigenetic regulators in PD 6.5 Erβ$^{KO}$ ovaries compared to PD 4.5 Erβ$^{KO}$ ovaries.

| Name | Chrom | Max group mean | Fold change | FDR p-value | ENSEMBL |
|---|---|---|---|---|---|
| Dpf1-203 | 1 | 5.3 | -9.51 | 0.04 | ENSRNOT00000108923 |
| Rnf40-203 | 1 | 5.6 | -12 | 7.68E-03 | ENSRNOT00000098677 |
| Kdm2a-202 | 1 | 37.55 | -2.03 | 2.27E-11 | ENSRNOT00000098321 |
| Btaf1-205 | 1 | 5.18 | 4.79 | 1.59E-05 | ENSRNOT00000113933 |
| Phc3-202 | 2 | 8.03 | 2.04 | 0.01 | ENSRNOT00000089477 |
| Phc3-201 | 2 | 8.39 | 2.27 | 0.04 | ENSRNOT00000066495 |
| Hdgf-202 | 2 | 32.36 | -7.44 | 0.02 | ENSRNOT00000065211 |
| Pogz-201 | 2 | 5.61 | 342.82 | 1.79E-07 | ENSRNOT00000082849 |
| Tp53bp1-203 | 3 | 9.84 | -14.92 | 3.07E-06 | ENSRNOT00000107143 |
| Tp53bp1-202 | 3 | 8.25 | -175.1 | 4.77E-16 | ENSRNOT00000106843 |
| Tp53bp1-201 | 3 | 30.36 | 2.11 | 3.86E-03 | ENSRNOT00000019025 |
| Chd6-205 | 3 | 13.81 | -11.59 | 5.19E-06 | ENSRNOT00000114568 |
| Kmt2c-203 | 4 | 5.63 | 19.88 | 2.31E-24 | ENSRNOT00000103015 |
| Prkag2-203 | 4 | 9.92 | -2.31 | 0.04 | ENSRNOT00000106083 |
| Tet3-201 | 4 | 12.07 | 3.15 | 3.68E-10 | ENSRNOT00000015296 |
| Mysm1-201 | 5 | 13.61 | 2.66 | 1.31E-04 | ENSRNOT00000039554 |
| Nasp-201 | 5 | 53.05 | 2.07 | 4.51E-10 | ENSRNOT00000022170 |
| Ppp4r3a-204 | 6 | 8.3 | -290.47 | 3.68E-10 | ENSRNOT00000114004 |
| Ubr5-201 | 7 | 25.59 | -3.71 | 1.12E-25 | ENSRNOT00000009115 |
| Phip-201 | 8 | 16.25 | 2.11 | 4.52E-13 | ENSRNOT00000011864 |
| Syncrip-202 | 8 | 45.9 | 2.38 | 2.77E-03 | ENSRNOT00000074515 |
| Setd2-201 | 8 | 11.56 | -709.46 | 1.89E-19 | ENSRNOT00000028409 |
| Sf3b1-202 | 9 | 116.78 | -2.07 | 1.98E-21 | ENSRNOT00000101949 |
| Ino80d-201 | 9 | 8.86 | 2.87 | 0.02 | ENSRNOT00000035879 |
| Glyr1-204 | 10 | 5.43 | 7.14 | 5.98E-03 | ENSRNOT00000106608 |
| Chd3-204 | 10 | 5.8 | 2.17 | 0.02 | ENSRNOT00000114886 |
| Kdm6b-202 | 10 | 16.42 | 2.07 | 1.66E-07 | ENSRNOT00000116399 |
| Dzip3-201 | 11 | 16.55 | 2.95 | 3.98E-06 | ENSRNOT00000002678 |
| Actl6b-201 | 12 | 5.04 | -486.42 | 0.02 | ENSRNOT00000001910 |
| Ncor2-205 | 12 | 32.93 | -2.12 | 1.96E-06 | ENSRNOT00000104620 |
| Ep400-204 | 12 | 8.46 | 11.02 | 1.72E-11 | ENSRNOT00000118850 |
| Supt16h-201 | 15 | 29.65 | 11.25 | 4.67E-06 | ENSRNOT00000016288 |
| Nsd1-205 | 17 | 21.12 | 3.03 | 8.78E-13 | ENSRNOT00000118233 |
| Nsd1-203 | 17 | 50.52 | 2 | 2.60E-22 | ENSRNOT00000107527 |
| Dek-202 | 17 | 5.29 | 392.8 | 0.03 | ENSRNOT00000115805 |
| Brd8-207 | 18 | 19.83 | -2.44 | 0.05 | ENSRNOT00000118939 |
| Kdm3b-203 | 18 | 15.74 | -3.83 | 8.78E-13 | ENSRNOT00000111068 |
| Hdac3-203 | 18 | 36.39 | -2.01 | 1.06E-05 | ENSRNOT00000084735 |
| Hdac3-201 | 18 | 11.4 | 8.21 | 0.01 | ENSRNOT00000060417 |
| Chd9-203 | 19 | 11.28 | 2.24 | 2.93E-06 | ENSRNOT00000078786 |
| Jmjd1c-202 | 20 | 9.46 | 2.59 | 1.31E-05 | ENSRNOT00000099911 |
| Tet1-201 | 20 | 5.2 | 2.62 | 3.44E-03 | ENSRNOT00000000302 |
| Huwe1-203 | X | 29.47 | 3.01 | 1.92E-14 | ENSRNOT00000098599 |
| Zmym3-204 | X | 18.9 | -2.04 | 3.75E-03 | ENSRNOT00000076709 |
| Ogt-203 | X | 100.89 | -97.48 | 2.80E-29 | ENSRNOT00000100805 |
| Smarca1-202 | X | 6.98 | 4.18 | 6.08E-03 | ENSRNOT00000114801 |

**Supplementary Table A1-4.** Differentially expressed epigenetic regulators in PD 6.5 Erβ$^{KO}$ ovaries compared to PD 4.5 Erβ$^{KO}$ ovaries.

| Name | Chrom | Max group mean | Fold change | FDR p-value | ENSEMBL |
|---|---|---|---|---|---|
| Jade1-202 | 2 | 5.49 | -41.27 | 1.69E-06 | ENSRNOT00000098865 |
| Chd6-202 | 3 | 10 | -2.29 | 0.03 | ENSRNOT00000089958 |
| Mta1-201 | 6 | 12.48 | 2.36 | 0.04 | ENSRNOT00000006521 |
| Ubr5-201 | 7 | 14.12 | 2.07 | 4.97E-04 | ENSRNOT00000009115 |
| Morf4l1-201 | 8 | 35.45 | 3,009.70 | 7.40E-03 | ENSRNOT00000080421 |
| Hdac3-201 | 18 | 11.4 | -5.36 | 0.02 | ENSRNOT00000060417 |
| Ogt-201 | X | 59.06 | -2.13 | 0.04 | ENSRNOT00000004692 |
| Taf9b-201 | X | 5.9 | -1,053.93 | 0.01 | ENSRNOT00000090833 |

### A.2. Transcription Factors

**Supplementary Table A2-1.** Differentially expressed transcription factors in PD 6.5 wildtype ovaries compared to PD 4.5 wildtype ovaries.

| Name | Chrom | Max group mean | Fold change | FDR p-value | ENSEMBL |
|---|---|---|---|---|---|
| Prr12-202 | 1 | 7.14 | -29.13 | 7.11E-11 | ENSRNOT00000094541 |
| Zfp518a-204 | 1 | 8.09 | 16.39 | 2.12E-07 | ENSRNOT00000105629 |
| Zfp455-202 | 2 | 10.98 | 2.51 | 0.03 | ENSRNOT00000070821 |
| Nfe2l2-203 | 3 | 21.52 | 2.76 | 0.03 | ENSRNOT00000110389 |
| Rbck1-203 | 3 | 17.55 | -3.8 | 5.46E-04 | ENSRNOT00000085995 |
| Chchd3-204 | 4 | 24.36 | 4.25 | 8.09E-05 | ENSRNOT00000107634 |
| Mysm1-201 | 5 | 13.68 | 2.41 | 3.36E-03 | ENSRNOT00000039554 |
| Nfia-203 | 5 | 7.58 | 2.34 | 0.03 | ENSRNOT00000086628 |
| Zbed4-203 | 7 | 8.17 | -2.51 | 0.01 | ENSRNOT00000103004 |
| Zbed4-201 | 7 | 6.87 | 2.99 | 6.25E-03 | ENSRNOT00000006050 |
| Ets1-204 | 8 | 12.83 | 2.82 | 5.00E-03 | ENSRNOT00000102138 |
| Hmg20a-202 | 8 | 7.24 | 2.02 | 0.02 | ENSRNOT00000102284 |
| Tgif1-202 | 9 | 5.79 | -77.26 | 2.32E-03 | ENSRNOT00000102373 |
| Glyr1-202 | 10 | 26.89 | -2.69 | 2.34E-07 | ENSRNOT00000095486 |
| Kat7-204 | 10 | 6.24 | -277.85 | 1.49E-05 | ENSRNOT00000109616 |
| Mlx-204 | 10 | 5.05 | -491.94 | 0.02 | ENSRNOT00000119770 |
| Bptf-202 | 10 | 5.38 | -2.74 | 3.09E-03 | ENSRNOT00000098328 |
| Foxk2-203 | 10 | 8.64 | 5.27 | 0.05 | ENSRNOT00000099287 |
| Nr5a2-201 | 13 | 6.75 | 2.48 | 0.04 | ENSRNOT00000000812 |
| Zfp322a-201 | 17 | 9.55 | 2.16 | 6.94E-05 | ENSRNOT00000023268 |
| Crem-204 | 17 | 16.43 | 2.08 | 7.65E-03 | ENSRNOT00000068545 |
| Tead3-203 | 20 | 13.87 | -2.69 | 0.04 | ENSRNOT00000107495 |
| Zfp711-202 | X | 5.48 | 4.31 | 6.11E-03 | ENSRNOT00000095292 |

**Supplementary Table A2-2.** Differentially expressed transcription factors in PD 8.5 wildtype ovaries compared to PD 6.5 wildtype ovaries.

| Name | Chrom | Max group mean | Fold change | FDR p-value | ENSEMBL |
|---|---|---|---|---|---|
| Peg3-202 | 1 | 11.32 | -2.87 | 1.40E-16 | ENSRNOT00000106399 |
| Zfp180-202 | 1 | 6.16 | -3.74 | 0.02 | ENSRNOT00000111936 |
| Nr2f2-201 | 1 | 56.91 | 2.46 | 2.80E-10 | ENSRNOT00000014152 |
| Deaf1-203 | 1 | 10.96 | -2.25 | 2.27E-04 | ENSRNOT00000098192 |
| Myrf-204 | 1 | 9.04 | -2.05 | 2.10E-03 | ENSRNOT00000116485 |
| Emx2-201 | 1 | 22.97 | -2.3 | 2.47E-11 | ENSRNOT00000012564 |
| Foxo1-201 | 2 | 77.47 | 3.5 | 1.02E-36 | ENSRNOT00000018244 |
| Sohlh2-201 | 2 | 7.7 | -3.04 | 3.33E-05 | ENSRNOT00000044424 |
| Rorc-204 | 2 | 8.04 | -3.74 | 6.72E-06 | ENSRNOT00000118722 |
| Setdb1-203 | 2 | 8.44 | -2.07 | 3.49E-07 | ENSRNOT00000111803 |
| Lef1-201 | 2 | 6.18 | -2.84 | 6.47E-03 | ENSRNOT00000013694 |
| Sohlh1-201 | 3 | 54.97 | -2.66 | 2.63E-21 | ENSRNOT00000051729 |
| Nr5a1-201 | 3 | 135.56 | 2.7 | 7.78E-44 | ENSRNOT00000017651 |
| Zeb2-201 | 3 | 7.28 | 2.25 | 3.34E-03 | ENSRNOT00000006350 |
| Sp5-201 | 3 | 6.39 | -3.6 | 4.39E-03 | ENSRNOT00000078819 |
| Nfe2l2-203 | 3 | 21.52 | -20.96 | 3.12E-12 | ENSRNOT00000110389 |
| Wt1-203 | 3 | 82.39 | -2.19 | 2.58E-22 | ENSRNOT00000102019 |
| Mafb-201 | 3 | 12.06 | 2.36 | 5.77E-03 | ENSRNOT00000021452 |
| Zhx3-201 | 3 | 6.7 | 2.44 | 9.03E-04 | ENSRNOT00000032588 |
| Tcfl5-201 | 3 | 7.86 | -2.51 | 5.13E-04 | ENSRNOT00000082266 |
| Sox18-201 | 3 | 32.79 | 2.13 | 1.83E-06 | ENSRNOT00000021790 |
| Chchd3-204 | 4 | 24.36 | -3.36 | 2.98E-03 | ENSRNOT00000107634 |
| Figla-201 | 4 | 166.52 | -3.62 | 4.26E-43 | ENSRNOT00000021272 |
| Klf15-201 | 4 | 8.23 | 2.23 | 3.40E-03 | ENSRNOT00000024011 |
| Pparg-202 | 4 | 11.18 | 3.67 | 2.42E-07 | ENSRNOT00000082969 |
| Plag1-201 | 5 | 11.91 | -2.48 | 3.83E-06 | ENSRNOT00000011721 |
| Nfia-201 | 5 | 11.25 | 2.54 | 3.69E-03 | ENSRNOT00000004017 |
| Epas1-202 | 6 | 37.88 | 2.22 | 4.37E-17 | ENSRNOT00000090420 |
| Epas1-201 | 6 | 9.44 | 2.63 | 7.03E-03 | ENSRNOT00000034991 |
| Fosl2-201 | 6 | 14.08 | 5.84 | 4.08E-06 | ENSRNOT00000107475 |
| Fosl2-202 | 6 | 30.23 | 2.41 | 5.43E-13 | ENSRNOT00000113775 |
| Esr2-204 | 6 | 10.74 | 2.48 | 3.23E-03 | ENSRNOT00000043602 |
| Fos-201 | 6 | 18.5 | 2.9 | 0.01 | ENSRNOT00000010712 |
| Osr2-201 | 7 | 73.25 | 2.7 | 6.42E-21 | ENSRNOT00000014879 |
| Myc-201 | 7 | 38.82 | 2.09 | 2.14E-08 | ENSRNOT00000006188 |
| Scx-201 | 7 | 10.6 | -2.8 | 0.03 | ENSRNOT00000029768 |
| Nr4a1-201 | 7 | 72.91 | 2.19 | 2.13E-03 | ENSRNOT00000010171 |
| Hoxc6-202 | 7 | 7.17 | -3.87 | 2.03E-35 | ENSRNOT00000107142 |
| Hoxc5-201 | 7 | 5.56 | -2.83 | 0.02 | ENSRNOT00000022247 |
| Hoxc4-201 | 7 | 8.55 | -2.26 | 2.60E-03 | ENSRNOT00000022398 |
| Pgr-201 | 8 | 5.76 | 2.49 | 2.45E-03 | ENSRNOT00000038313 |
| Zfp426-201 | 8 | 6.51 | -3.44 | 2.10E-05 | ENSRNOT00000038069 |
| Zglp1-201 | 8 | 26.97 | -2.94 | 1.85E-05 | ENSRNOT00000068598 |
| Ets1-204 | 8 | 12.83 | -6.19 | 1.27E-08 | ENSRNOT00000102138 |
| Hmg20a-202 | 8 | 7.24 | -3.24 | 8.95E-04 | ENSRNOT00000102284 |
| Safb-205 | 9 | 12.48 | -2.06 | 1.87E-06 | ENSRNOT00000115324 |
| L3mbtl4-201 | 9 | 5.77 | 2.12 | 0.02 | ENSRNOT00000029210 |
| Glyr1-202 | 10 | 20.62 | 2.16 | 2.01E-09 | ENSRNOT00000095486 |

| | | | | | |
|---|---|---|---|---|---|
| Sgsm2-203 | 10 | 11.24 | 2.19 | 3.13E-04 | ENSRNOT00000092314 |
| Hic1-201 | 10 | 12.52 | 3.06 | 8.35E-06 | ENSRNOT00000004143 |
| Kat7-204 | 10 | 6.46 | 327.56 | 2.37E-08 | ENSRNOT00000109616 |
| Mlx-204 | 10 | 9.31 | 1,075.75 | 2.97E-03 | ENSRNOT00000119770 |
| Gtf2i-205 | 12 | 19.82 | 2.15 | 1.62E-06 | ENSRNOT00000098927 |
| Nr5a2-202 | 13 | 123.63 | 2.03 | 2.54E-29 | ENSRNOT00000097279 |
| Lhx9-201 | 13 | 28.68 | -5.35 | 3.09E-33 | ENSRNOT00000014218 |
| Lhx9-203 | 13 | 6.82 | -5.46 | 3.26E-05 | ENSRNOT00000116473 |
| Pbx1-202 | 13 | 18.13 | 2.08 | 7.47E-09 | ENSRNOT00000103202 |
| Zfp326-204 | 14 | 11.49 | -2.25 | 1.25E-03 | ENSRNOT00000095214 |
| Sox7-201 | 15 | 8.48 | 2.67 | 1.80E-04 | ENSRNOT00000016503 |
| Dzip1-201 | 15 | 9.69 | -2.68 | 1.53E-03 | ENSRNOT00000038596 |
| Crem-210 | 17 | 6.1 | -1,627.15 | 5.15E-03 | ENSRNOT00000115950 |
| Egr1-201 | 18 | 52.87 | 2.2 | 5.98E-04 | ENSRNOT00000026303 |
| Prdm6-201 | 18 | 7.52 | 2.73 | 4.10E-03 | ENSRNOT00000047755 |
| Terb1-201 | 19 | 32.18 | -2.52 | 5.29E-36 | ENSRNOT00000087469 |
| Irx5-202 | 19 | 7.32 | -2.57 | 3.65E-04 | ENSRNOT00000089975 |
| Irx3-201 | 19 | 23.73 | -4.21 | 1.88E-29 | ENSRNOT00000015583 |
| Zfp612-202 | 19 | 13.81 | -2.59 | 5.29E-32 | ENSRNOT00000112604 |
| Zfp711-202 | X | 5.48 | -3.04 | 0.03 | ENSRNOT00000095292 |

**Supplementary Table A2-3.** Differentially expressed transcription factors in PD 6.5 Erβ$^{KO}$ ovaries compared to PD 4.5 Erβ$^{KO}$ ovaries.

| Name | Chrom | Max group mean | Fold change | FDR p-value | ENSEMBL |
|---|---|---|---|---|---|
| Peg3-202 | 1 | 14.68 | 2.16 | 2.00E-10 | ENSRNOT00000106399 |
| Dpf1-203 | 1 | 5.3 | -9.38 | 0.03 | ENSRNOT00000108923 |
| Zfp260-202 | 1 | 6.97 | -141.72 | 1.46E-09 | ENSRNOT00000074301 |
| Zfp260-204 | 1 | 33.81 | 2.29 | 4.08E-12 | ENSRNOT00000113539 |
| Tead1-203 | 1 | 6.48 | 3.47 | 0.02 | ENSRNOT00000096192 |
| Tead1-201 | 1 | 9.46 | 2.85 | 0.03 | ENSRNOT00000021020 |
| Kdm2a-202 | 1 | 37.55 | -2.04 | 3.31E-10 | ENSRNOT00000098321 |
| Myrf-204 | 1 | 15.57 | -2.96 | 2.58E-09 | ENSRNOT00000116485 |
| Lcor-202 | 1 | 8.11 | 2.63 | 4.76E-04 | ENSRNOT00000076842 |
| Zbed3-201 | 2 | 32.37 | -3.9 | 1.18E-04 | ENSRNOT00000040983 |
| Zfp704-201 | 2 | 17.91 | 2.55 | 1.21E-08 | ENSRNOT00000080313 |
| Zfp639-203 | 2 | 5.49 | 4.77 | 0.01 | ENSRNOT00000100539 |
| Creb3l4-201 | 2 | 6.93 | -3.13 | 0.04 | ENSRNOT00000032062 |
| Rorc-204 | 2 | 11.39 | -2.17 | 1.05E-03 | ENSRNOT00000118722 |
| Arnt-202 | 2 | 12.58 | 2.02 | 0.02 | ENSRNOT00000064442 |
| Lef1-201 | 2 | 8.29 | -2.99 | 3.10E-03 | ENSRNOT00000013694 |
| Zeb2-201 | 3 | 9.53 | -3.04 | 0.01 | ENSRNOT00000006350 |
| Zeb2-203 | 3 | 12.72 | 5.67 | 0.01 | ENSRNOT00000085801 |
| Nfe2l2-203 | 3 | 33.48 | -2.77 | 1.57E-04 | ENSRNOT00000110389 |
| Phf20-202 | 3 | 6.23 | 3.26 | 5.25E-03 | ENSRNOT00000095858 |
| Zhx3-201 | 3 | 7.11 | 2.17 | 0.01 | ENSRNOT00000032588 |
| Nrf1-205 | 4 | 5.69 | 2.03 | 0.03 | ENSRNOT00000105926 |
| Zfp786-201 | 4 | 5.5 | 869.67 | 0.01 | ENSRNOT00000038550 |
| Tet3-201 | 4 | 12.07 | 3.01 | 1.53E-08 | ENSRNOT00000015296 |
| Nr2c2-201 | 4 | 24.29 | 2.28 | 5.98E-03 | ENSRNOT00000014353 |
| Zfp384-203 | 4 | 5.92 | -1,163.82 | 6.71E-03 | ENSRNOT00000036654 |
| Nfx1-201 | 5 | 14.33 | 9.34 | 2.04E-12 | ENSRNOT00000060691 |
| Mysm1-201 | 5 | 13.61 | 2.66 | 3.09E-04 | ENSRNOT00000039554 |
| Pou3f1-201 | 5 | 5.23 | -2.66 | 0.03 | ENSRNOT00000074492 |
| Esr2-203 | 6 | 10.76 | 6.79 | 8.65E-05 | ENSRNOT00000042682 |
| Nfic-203 | 7 | 11.89 | 2.41 | 0.02 | ENSRNOT00000097025 |
| Eea1-206 | 7 | 11.8 | 2.8 | 2.49E-03 | ENSRNOT00000117665 |
| Nr4a1-201 | 7 | 15.75 | 3.68 | 3.95E-05 | ENSRNOT00000010171 |
| Zfp426-201 | 8 | 9.71 | -2.01 | 0.03 | ENSRNOT00000038069 |
| Tcf12-201 | 8 | 45.59 | 2.17 | 6.43E-09 | ENSRNOT00000081185 |
| Safb2-202 | 9 | 5.17 | 3.45 | 5.28E-03 | ENSRNOT00000097383 |
| Glyr1-204 | 10 | 5.43 | 7.16 | 0.01 | ENSRNOT00000106608 |
| Hic1-201 | 10 | 7.99 | 2.38 | 5.23E-03 | ENSRNOT00000004143 |
| Mlx-202 | 10 | 6.56 | 698.55 | 0.02 | ENSRNOT00000099833 |
| Bptf-204 | 10 | 5.91 | 2.19 | 0.02 | ENSRNOT00000104873 |
| Foxk2-203 | 10 | 17.69 | -4.64 | 3.67E-05 | ENSRNOT00000099287 |
| Son-201 | 11 | 34.05 | 6.27 | 4.45E-06 | ENSRNOT00000002769 |
| Cux1-204 | 12 | 7.52 | 3.8 | 1.69E-07 | ENSRNOT00000112152 |
| Zbed6-202 | 13 | 34.35 | 2.29 | 5.91E-12 | ENSRNOT00000115014 |
| Zbtb18-204 | 13 | 28.35 | -2.02 | 4.10E-06 | ENSRNOT00000105416 |
| Nr1d2-203 | 15 | 18.92 | -4.77 | 0.02 | ENSRNOT00000106171 |
| Hmbox1-202 | 15 | 12.86 | 2.1 | 6.89E-10 | ENSRNOT00000109493 |
| Dzip1-201 | 15 | 22.78 | -2.88 | 0.03 | ENSRNOT00000038596 |

| | | | | | |
|---|---|---|---|---|---|
| Zbtb7c-202 | 18 | 6 | -12.33 | 0.02 | ENSRNOT00000101836 |
| Zfp423-203 | 19 | 6.39 | 3.22 | 7.91E-04 | ENSRNOT00000101895 |
| Zfp423-201 | 19 | 11.18 | -3.26 | 1.62E-04 | ENSRNOT00000020028 |
| Zfp57-201 | 20 | 98.07 | 2.17 | 1.39E-17 | ENSRNOT00000011704 |
| Tet1-201 | 20 | 5.2 | 2.62 | 3.22E-04 | ENSRNOT00000000302 |
| Ar-201 | X | 9.59 | 3.38 | 1.41E-06 | ENSRNOT00000009129 |
| Mecp2-201 | X | 11.65 | 2.64 | 0.05 | ENSRNOT00000085723 |

**Supplementary Table A2-4.** Differentially expressed transcription factors in PD 8.5 Erβ$^{KO}$ ovaries compared to PD 6.5 Erβ$^{KO}$ ovaries.

| Name | Chrom | Max group mean | Fold change | FDR p-value | ENSEMBL |
|---|---|---|---|---|---|
| Zbed3-201 | 2 | 10.19 | -16.55 | 2.91E-03 | ENSRNOT00000040983 |
| Nfe2l2-203 | 3 | 11.16 | -18.09 | 2.60E-06 | ENSRNOT00000110389 |
| Zfp410-204 | 6 | 7.23 | 9.89 | 0.04 | ENSRNOT00000109998 |
| Eea1-202 | 7 | 5.43 | 9.65 | 7.95E-03 | ENSRNOT00000087297 |
| Eea1-206 | 7 | 11.8 | -2.15 | 0.04 | ENSRNOT00000117665 |
| Hoxc6-202 | 7 | 5.93 | -2.52 | 0.04 | ENSRNOT00000107142 |
| L3mbtl4-201 | 9 | 6.37 | 2.08 | 0.05 | ENSRNOT00000029210 |
| Camta2-203 | 10 | 5.24 | 6.75 | 0.04 | ENSRNOT00000109888 |
| Zbtb7c-202 | 18 | 9.55 | 37.5 | 3.66E-09 | ENSRNOT00000101836 |
| Tead3-203 | 20 | 9.78 | -8.09 | 5.47E-03 | ENSRNOT00000107495 |

### A.3. Downstream Genes

| **Supplementary Table A3-1.** Differentially expressed downstream genes in PD6.5 wildtype ovaries compared to PD 4.5 wildtype ovaries. | | | | | |
|---|---|---|---|---|---|
| Name | Chrom | Max group mean | Fold change | FDR p-value | ENSEMBL |
| Utrn-205 | 1 | 7.5 | -2.18 | 0.05 | ENSRNOT00000110616 |
| Utrn-201 | 1 | 7.91 | 2.39 | 1.66E-06 | ENSRNOT00000016273 |
| Rps12-201 | 1 | 218.44 | 2.51 | 1.14E-12 | ENSRNOT00000022184 |
| Rps12-202 | 1 | 2,197.74 | 2.4 | 1.01E-27 | ENSRNOT00000115390 |
| Slc12a7-202 | 1 | 11.62 | -2.34 | 8.23E-07 | ENSRNOT00000078039 |
| Mtrr-201 | 1 | 6.64 | 2.86 | 0.02 | ENSRNOT00000024041 |
| Akap12-202 | 1 | 15.96 | -2.17 | 3.83E-05 | ENSRNOT00000060767 |
| Scaf8-201 | 1 | 16.52 | -2.94 | 3.26E-05 | ENSRNOT00000065733 |
| Mrpl18-203 | 1 | 15.89 | 3.3 | 3.12E-03 | ENSRNOT00000101637 |
| AABR07001512.1-201 | 1 | 26.79 | 99.79 | 5.79E-07 | ENSRNOT00000023387 |
| Sft2d1-202 | 1 | 14.83 | 5.9 | 0.02 | ENSRNOT00000113562 |
| Psmb1-201 | 1 | 10.04 | 3.72 | 0.02 | ENSRNOT00000002037 |
| Nlrp5-201 | 1 | 44.76 | -2.28 | 3.08E-14 | ENSRNOT00000041542 |
| Nlrp5-202 | 1 | 356.31 | -2.47 | 8.14E-27 | ENSRNOT00000097841 |
| Tnni3-201 | 1 | 21.85 | 2.26 | 0.04 | ENSRNOT00000024640 |
| LOC100364427-201 | 1 | 69.6 | 2.62 | 7.69E-06 | ENSRNOT00000073324 |
| Rpl9-201 | 1 | 4,344.69 | 2.62 | 6.80E-31 | ENSRNOT00000052231 |
| Slc8a2-202 | 1 | 5.81 | -10.68 | 1.35E-13 | ENSRNOT00000086396 |
| Dmwd-201 | 1 | 8.37 | 2.46 | 0.01 | ENSRNOT00000020134 |
| Rps19-201 | 1 | 4,177.16 | 2.04 | 7.11E-18 | ENSRNOT00000027246 |
| Ltbp4-201 | 1 | 73.62 | -3.92 | 6.80E-31 | ENSRNOT00000028322 |
| Rps16-202 | 1 | 1,182.53 | 2.03 | 7.84E-18 | ENSRNOT00000093558 |
| Rps16-201 | 1 | 1,600.71 | 2.01 | 1.06E-17 | ENSRNOT00000026576 |
| Spint2-201 | 1 | 38.16 | -2,330.83 | 3.61E-04 | ENSRNOT00000028006 |
| Lsm14a-201 | 1 | 7.52 | 3.83 | 6.05E-03 | ENSRNOT00000030186 |
| Ankrd27-202 | 1 | 12.12 | -2.5 | 7.85E-07 | ENSRNOT00000081465 |
| Etfb-202 | 1 | 38.56 | 2.05 | 1.91E-03 | ENSRNOT00000118657 |
| Fv1-202 | 1 | 14.19 | 2.29 | 1.01E-06 | ENSRNOT00000096151 |
| Prr12-202 | 1 | 7.14 | -29.96 | 1.03E-11 | ENSRNOT00000094541 |
| Rpl13a-201 | 1 | 2,867.71 | 2.06 | 7.56E-18 | ENSRNOT00000027976 |
| Ppp1r15a-201 | 1 | 10.04 | 2.39 | 0.04 | ENSRNOT00000044788 |
| Rpl18-204 | 1 | 16.74 | -4.47 | 7.52E-03 | ENSRNOT00000103343 |
| Gtf2h1-201 | 1 | 8.82 | 2.7 | 8.23E-03 | ENSRNOT00000066791 |
| Herc2-201 | 1 | 8.45 | -2.98 | 3.14E-04 | ENSRNOT00000019245 |
| Mcee-201 | 1 | 16.47 | 3.29 | 0.03 | ENSRNOT00000021884 |
| Mcee-203 | 1 | 7.35 | -326.81 | 5.55E-03 | ENSRNOT00000101373 |
| St8sia2-201 | 1 | 5.07 | -2.92 | 2.79E-03 | ENSRNOT00000113519 |
| Klhl25-201 | 1 | 10.45 | -2.43 | 3.19E-03 | ENSRNOT00000014553 |
| Mfge8-202 | 1 | 178.46 | -2.17 | 2.68E-09 | ENSRNOT00000079100 |
| Anpep-201 | 1 | 15.67 | -2.1 | 4.17E-06 | ENSRNOT00000020002 |
| LOC100365810-202 | 1 | 5,605.74 | 2.42 | 1.30E-26 | ENSRNOT00000100646 |
| Tmem126a-201 | 1 | 21.05 | 2.02 | 2.92E-06 | ENSRNOT00000038333 |
| Rps3-204 | 1 | 1,584.42 | 2.26 | 1.50E-21 | ENSRNOT00000114051 |
| Rps3-201 | 1 | 1,794.90 | 2 | 6.02E-17 | ENSRNOT00000023935 |
| C2cd3-201 | 1 | 6.65 | -2.27 | 4.65E-03 | ENSRNOT00000023802 |
| Mrpl48-204 | 1 | 26.02 | 2.23 | 0.03 | ENSRNOT00000110624 |

| | | | | | |
|---|---|---|---|---|---|
| Fam168a-203 | 1 | 6.59 | 101.49 | 2.16E-06 | ENSRNOT00000097710 |
| Hbb-210 | 1 | 1,826.96 | -2.9 | 1.91E-03 | ENSRNOT00000114441 |
| Dennd2b-208 | 1 | 23.19 | -2.09 | 2.76E-05 | ENSRNOT00000114796 |
| Dennd2b-202 | 1 | 9.81 | -2.09 | 1.18E-03 | ENSRNOT00000099927 |
| Tead1-201 | 1 | 8.53 | 2.25 | 0.04 | ENSRNOT00000021020 |
| Rps13-203 | 1 | 1,210.85 | 2.21 | 4.75E-23 | ENSRNOT00000118821 |
| Rps15a-201 | 1 | 2,470.86 | 2.37 | 5.48E-26 | ENSRNOT00000092282 |
| Rps15a-202 | 1 | 90.66 | 2.26 | 4.40E-04 | ENSRNOT00000092513 |
| Vps35l-202 | 1 | 10.94 | -3.87 | 5.13E-06 | ENSRNOT00000083863 |
| Spns1-203 | 1 | 11.5 | -3.5 | 3.03E-03 | ENSRNOT00000102967 |
| LOC100361060-201 | 1 | 318.96 | 2.43 | 2.90E-33 | ENSRNOT00000045531 |
| LOC100302465-202 | 1 | 11.95 | -8.69 | 2.29E-06 | ENSRNOT00000073836 |
| Cttn-203 | 1 | 8.6 | 7.62 | 0.03 | ENSRNOT00000054859 |
| Ndufv1-201 | 1 | 13.29 | -5.93 | 3.73E-03 | ENSRNOT00000024517 |
| Brms1-204 | 1 | 8.08 | 8.03 | 7.27E-03 | ENSRNOT00000113489 |
| Rtn3-203 | 1 | 44.81 | -2.08 | 2.67E-08 | ENSRNOT00000099817 |
| Prpf19-202 | 1 | 36.7 | -3.03 | 5.10E-11 | ENSRNOT00000095505 |
| Fam111a-201 | 1 | 122.64 | 2.15 | 1.31E-07 | ENSRNOT00000016152 |
| Gldc-204 | 1 | 5.68 | -2.26 | 0.05 | ENSRNOT00000110722 |
| Kif11-202 | 1 | 11.91 | 7.12 | 4.48E-13 | ENSRNOT00000095978 |
| Rbp4-201 | 1 | 57.58 | 2.18 | 3.79E-03 | ENSRNOT00000021055 |
| Sorbs1-205 | 1 | 5.74 | -2.37 | 0.04 | ENSRNOT00000021347 |
| Zfp518a-204 | 1 | 8.09 | 14.41 | 2.88E-07 | ENSRNOT00000105629 |
| Scd-201 | 1 | 5.88 | 2.04 | 0.03 | ENSRNOT00000018447 |
| Arl3-201 | 1 | 32.37 | 2.24 | 8.74E-03 | ENSRNOT00000027093 |
| Cyp17a1-201 | 1 | 105.73 | 5.95 | 1.14E-11 | ENSRNOT00000027160 |
| Atp5mk-201 | 1 | 802.04 | 2.54 | 6.80E-31 | ENSRNOT00000027496 |
| Sh3pxd2a-201 | 1 | 13.28 | 2.28 | 9.25E-05 | ENSRNOT00000027574 |
| Gsto1-201 | 1 | 118.45 | 2.05 | 3.72E-08 | ENSRNOT00000016851 |
| RGD1561333-201 | 1 | 20.34 | 2.23 | 0.05 | ENSRNOT00000048691 |
| Smc3-201 | 1 | 5.35 | -1,143.52 | 1.77E-04 | ENSRNOT00000019560 |
| Afap1l2-203 | 1 | 8.02 | -2.83 | 0.01 | ENSRNOT00000105824 |
| Cox7c-201 | 2 | 360.68 | 2.17 | 2.48E-21 | ENSRNOT00000051895 |
| Rps23-201 | 2 | 2,605.86 | 2.01 | 2.68E-17 | ENSRNOT00000022348 |
| Ak6-202 | 2 | 34.76 | 2.05 | 2.48E-04 | ENSRNOT00000061166 |
| Erbin-202 | 2 | 10.92 | 2.38 | 6.94E-07 | ENSRNOT00000095613 |
| Pde4d-204 | 2 | 6.96 | 3.2 | 4.10E-03 | ENSRNOT00000101684 |
| Ndufs4-202 | 2 | 32.05 | 8.82 | 1.07E-10 | ENSRNOT00000093889 |
| Nnt-205 | 2 | 5.18 | 1,145.68 | 3.28E-03 | ENSRNOT00000112693 |
| Rps15al4-201 | 2 | 89.12 | 2.33 | 6.40E-05 | ENSRNOT00000041483 |
| ENSRNOT00000113072 | 2 | 177.1 | 2.23 | 3.47E-15 | ENSRNOT00000113072 |
| Zfr-203 | 2 | 32.27 | 2.22 | 1.51E-05 | ENSRNOT00000087636 |
| Myo10-203 | 2 | 16.57 | -12.7 | 2.14E-20 | ENSRNOT00000102421 |
| ENSRNOT00000107628 | 2 | 14.03 | 2.16 | 6.58E-04 | ENSRNOT00000107628 |
| Car3-201 | 2 | 15.27 | 5.53 | 2.02E-05 | ENSRNOT00000014180 |
| Car3-202 | 2 | 40.66 | 6.27 | 5.75E-33 | ENSRNOT00000105296 |
| Rbis-204 | 2 | 54.82 | 2.49 | 4.33E-07 | ENSRNOT00000104515 |
| Rbis-202 | 2 | 120.46 | 2.43 | 5.58E-11 | ENSRNOT00000059467 |
| Fabp4-201 | 2 | 86.63 | 5.65 | 1.41E-11 | ENSRNOT00000014701 |
| Pde7a-204 | 2 | 6.4 | -8.09 | 0.05 | ENSRNOT00000108651 |
| Rpl22l1-201 | 2 | 872.71 | 2.89 | 7.45E-42 | ENSRNOT00000015756 |

| | | | | | |
|---|---|---|---|---|---|
| Dnajc19-206 | 2 | 80.98 | 2.59 | 1.48E-05 | ENSRNOT00000118548 |
| Dnajc19-205 | 2 | 81.81 | 2.06 | 7.82E-06 | ENSRNOT00000118294 |
| Exosc9-203 | 2 | 10.71 | 3.38 | 7.84E-03 | ENSRNOT00000108461 |
| Trpc3-201 | 2 | 5.28 | 2.3 | 0.04 | ENSRNOT00000046700 |
| Larp1b-202 | 2 | 13.43 | -6.58 | 4.36E-04 | ENSRNOT00000090861 |
| Etfdh-201 | 2 | 19.37 | 2.08 | 9.71E-06 | ENSRNOT00000013262 |
| Rps3a-201 | 2 | 3,990.58 | 2.17 | 2.78E-21 | ENSRNOT00000016329 |
| Dclk2-204 | 2 | 14.26 | -2.58 | 1.30E-05 | ENSRNOT00000079954 |
| Rps27-201 | 2 | 4,765.79 | 2.3 | 1.52E-23 | ENSRNOT00000022897 |
| S100a6-201 | 2 | 179.31 | 2.08 | 8.42E-10 | ENSRNOT00000015612 |
| Lce1m-201 | 2 | 8.9 | 6.75 | 7.40E-03 | ENSRNOT00000012691 |
| ENSRNOT00000096678 | 2 | 9.43 | 5.62 | 0.02 | ENSRNOT00000096678 |
| Cgn-203 | 2 | 37.58 | -2.23 | 3.42E-08 | ENSRNOT00000116247 |
| Ecm1-201 | 2 | 53.16 | -2.55 | 7.76E-17 | ENSRNOT00000028740 |
| Ecm1-202 | 2 | 103.89 | -2.01 | 3.26E-13 | ENSRNOT00000105287 |
| Hsd3b3-201 | 2 | 338.05 | 2.84 | 2.13E-06 | ENSRNOT00000056172 |
| Hsd3b1-201 | 2 | 816.72 | 2.27 | 8.79E-03 | ENSRNOT00000086835 |
| Hao2-201 | 2 | 5.45 | 97.74 | 2.83E-04 | ENSRNOT00000046942 |
| Rap1a-203 | 2 | 31.47 | 2.81 | 0.02 | ENSRNOT00000110219 |
| Hiat1-ps1-205 | 2 | 6.26 | 31.15 | 3.02E-05 | ENSRNOT00000097342 |
| Rpl34-201 | 2 | 2,027.21 | 2.11 | 3.90E-19 | ENSRNOT00000009046 |
| Odf2l-204 | 2 | 8.28 | 2 | 0.04 | ENSRNOT00000119248 |
| Prkacb-202 | 2 | 10.31 | 37.13 | 1.20E-14 | ENSRNOT00000109432 |
| Depdc1-201 | 2 | 11.52 | 2.47 | 6.48E-06 | ENSRNOT00000012667 |
| Kcnt1-203 | 3 | 7.59 | -2.09 | 0.02 | ENSRNOT00000104747 |
| Qsox2-202 | 3 | 5.25 | 105.68 | 2.73E-09 | ENSRNOT00000116146 |
| Rpl9-ps31-201 | 3 | 41.86 | 2.35 | 4.79E-04 | ENSRNOT00000042201 |
| Usp20-203 | 3 | 7.29 | -6.67 | 1.96E-03 | ENSRNOT00000114450 |
| Ass1-202 | 3 | 21.13 | 2.43 | 1.22E-03 | ENSRNOT00000093365 |
| St6galnac6-202 | 3 | 9.98 | -3.04 | 0.02 | ENSRNOT00000094850 |
| AABR07051450.1-202 | 3 | 46.51 | 2.01 | 1.84E-06 | ENSRNOT00000091048 |
| Zbtb43-203 | 3 | 7.77 | -2.33 | 3.05E-03 | ENSRNOT00000108027 |
| Gapvd1-202 | 3 | 7.07 | -3.08 | 0.02 | ENSRNOT00000096520 |
| Gsn-201 | 3 | 14.98 | -2.73 | 0.05 | ENSRNOT00000025857 |
| Rps20-201 | 3 | 6,028.30 | 2.51 | 2.88E-28 | ENSRNOT00000041813 |
| Scai-203 | 3 | 11.55 | 2.06 | 1.35E-04 | ENSRNOT00000104956 |
| LOC100362366-201 | 3 | 10.78 | 6.4 | 0.02 | ENSRNOT00000037043 |
| Orc4-203 | 3 | 21.66 | 2.08 | 2.55E-04 | ENSRNOT00000103065 |
| Marchf7-202 | 3 | 10.53 | 6.48 | 4.80E-06 | ENSRNOT00000085563 |
| Fap-201 | 3 | 36.42 | 2.09 | 1.13E-06 | ENSRNOT00000008148 |
| Hnrnpa3-204 | 3 | 19.61 | 126.68 | 1.16E-08 | ENSRNOT00000114122 |
| Nfe2l2-203 | 3 | 21.52 | 2.69 | 0.02 | ENSRNOT00000110389 |
| Fkbp7-201 | 3 | 16.8 | 2.07 | 0.03 | ENSRNOT00000015722 |
| Zc3h15-202 | 3 | 7.12 | 228.15 | 1.82E-06 | ENSRNOT00000094114 |
| RGD1560017-201 | 3 | 55.47 | 2.45 | 3.68E-04 | ENSRNOT00000049695 |
| Eif3m-205 | 3 | 33.5 | 5.51 | 4.89E-05 | ENSRNOT00000117477 |
| Immp1l-202 | 3 | 10.76 | 2.17 | 0.02 | ENSRNOT00000100315 |
| Aven-204 | 3 | 10.67 | 17.65 | 4.69E-04 | ENSRNOT00000118480 |
| Rasgrp1-201 | 3 | 6.37 | -2.32 | 0.03 | ENSRNOT00000007687 |
| Nusap1-201 | 3 | 7.89 | -296.12 | 1.99E-06 | ENSRNOT00000052312 |
| Nusap1-203 | 3 | 9.22 | 2.91 | 9.13E-03 | ENSRNOT00000095179 |

| | | | | | |
|---|---|---|---|---|---|
| Bub1-202 | 3 | 6.67 | 2.76 | 1.15E-03 | ENSRNOT00000111791 |
| Anapc1-203 | 3 | 11.94 | -2.3 | 2.72E-06 | ENSRNOT00000111998 |
| ENSRNOT00000028881 | 3 | 105.04 | 3.19 | 1.67E-11 | ENSRNOT00000028881 |
| Snrpb2-202 | 3 | 50.67 | 2.1 | 5.21E-06 | ENSRNOT00000097908 |
| Smim26-201 | 3 | 55.86 | 2.62 | 4.64E-09 | ENSRNOT00000055588 |
| Rbck1-203 | 3 | 17.55 | -3.59 | 6.52E-04 | ENSRNOT00000085995 |
| Eif2s2-202 | 3 | 149.17 | 2.18 | 1.55E-23 | ENSRNOT00000110685 |
| Rpn2-201 | 3 | 40.11 | -3.48 | 1.67E-15 | ENSRNOT00000068135 |
| Chd6-205 | 3 | 8.46 | -7.21 | 3.35E-04 | ENSRNOT00000114568 |
| Chd6-202 | 3 | 10.67 | -5.37 | 2.98E-06 | ENSRNOT00000089958 |
| Svs5-201 | 3 | 31.11 | 14.99 | 5.11E-06 | ENSRNOT00000018537 |
| Zswim1-202 | 3 | 13.72 | -2.7 | 4.43E-04 | ENSRNOT00000106851 |
| Pfdn4-201 | 3 | 118.96 | 2.45 | 1.06E-17 | ENSRNOT00000070809 |
| Psma7-204 | 3 | 30.88 | 3.25 | 4.88E-04 | ENSRNOT00000116159 |
| Rps21-201 | 3 | 1,871.54 | 2.33 | 3.63E-25 | ENSRNOT00000084027 |
| Uckl1-207 | 3 | 5.35 | -507.19 | 0.02 | ENSRNOT00000113965 |
| Tomm7-201 | 4 | 447.85 | 2.06 | 5.52E-19 | ENSRNOT00000065962 |
| Bet1-202 | 4 | 8.54 | -586.67 | 1.78E-03 | ENSRNOT00000106013 |
| Ppp1r9a-202 | 4 | 7.36 | 2.2 | 1.29E-03 | ENSRNOT00000104466 |
| Ndufa4-203 | 4 | 479.78 | 2.31 | 1.69E-22 | ENSRNOT00000112879 |
| Phf14-208 | 4 | 5.77 | 2 | 9.58E-05 | ENSRNOT00000108109 |
| Cadps2-203 | 4 | 6.13 | -2.01 | 0.05 | ENSRNOT00000114656 |
| Podxl-201 | 4 | 23.82 | -2.72 | 1.53E-04 | ENSRNOT00000016991 |
| Chchd3-204 | 4 | 24.36 | 4.05 | 3.92E-04 | ENSRNOT00000107634 |
| Akr1b10-204 | 4 | 19.55 | 2.33 | 6.94E-03 | ENSRNOT00000116609 |
| Ubn2-201 | 4 | 6.31 | 2.7 | 2.69E-04 | ENSRNOT00000064175 |
| AC113910.1-201 | 4 | 127.02 | 2.15 | 9.98E-08 | ENSRNOT00000041621 |
| Lsm5-204 | 4 | 138.95 | 2.17 | 2.37E-07 | ENSRNOT00000114266 |
| Lsm5-205 | 4 | 57.75 | 2.29 | 5.71E-05 | ENSRNOT00000118324 |
| Mrpl35-201 | 4 | 6.32 | 122.85 | 2.65E-04 | ENSRNOT00000012021 |
| Immt-203 | 4 | 12.66 | -3.62 | 1.46E-03 | ENSRNOT00000098740 |
| Ggcx-201 | 4 | 25.52 | -2 | 3.86E-05 | ENSRNOT00000017927 |
| Capg-201 | 4 | 5.9 | 12.3 | 0.02 | ENSRNOT00000018562 |
| LOC687679-201 | 4 | 696.4 | 2.22 | 4.02E-24 | ENSRNOT00000022367 |
| Pcyox1-203 | 4 | 7.94 | -12.41 | 0.01 | ENSRNOT00000105157 |
| Rpl22-201 | 4 | 1,310.90 | 2.53 | 4.64E-31 | ENSRNOT00000034810 |
| Hmces-201 | 4 | 16.73 | -2.79 | 0.04 | ENSRNOT00000016137 |
| LOC685085-201 | 4 | 509.03 | 2.85 | 9.72E-38 | ENSRNOT00000099455 |
| Fbln2-203 | 4 | 84.85 | -2.07 | 7.97E-07 | ENSRNOT00000115833 |
| Tmem43-205 | 4 | 6.83 | -279.57 | 6.31E-05 | ENSRNOT00000107696 |
| Lrrn1-201 | 4 | 7 | -2.07 | 3.59E-04 | ENSRNOT00000008879 |
| Thumpd3-201 | 4 | 26.26 | 2.06 | 8.86E-06 | ENSRNOT00000009464 |
| Hnrnpf-201 | 4 | 71.5 | 2.26 | 1.14E-13 | ENSRNOT00000019529 |
| Wnk1-204 | 4 | 13.93 | 2.06 | 4.91E-03 | ENSRNOT00000080788 |
| Wnk1-203 | 4 | 7.37 | -2.84 | 0.02 | ENSRNOT00000079513 |
| A2m-201 | 4 | 13.56 | -2.18 | 2.65E-06 | ENSRNOT00000019346 |
| Rad51ap1-202 | 4 | 8.87 | -5.46 | 0.03 | ENSRNOT00000119755 |
| Tspan11-202 | 4 | 9.74 | 2.89 | 2.80E-03 | ENSRNOT00000103340 |
| Clec2g-201 | 4 | 68.95 | 2.23 | 6.30E-15 | ENSRNOT00000083188 |
| Eloc-201 | 5 | 19.71 | 2.08 | 2.21E-06 | ENSRNOT00000050236 |
| Lactb2-201 | 5 | 5.75 | 14.69 | 5.57E-03 | ENSRNOT00000010369 |

| | | | | | |
|---|---|---|---|---|---|
| Asph-210 | 5 | 7.21 | 2.4 | 3.32E-03 | ENSRNOT00000118158 |
| ENSRNOT00000113061 | 5 | 410.78 | -2.21 | 0.05 | ENSRNOT00000113061 |
| Plekhf2-202 | 5 | 8.59 | -8.93 | 8.03E-05 | ENSRNOT00000094306 |
| LOC120102832-201 | 5 | 5.82 | 19.23 | 4.33E-08 | ENSRNOT00000114212 |
| Tstd3-201 | 5 | 27.38 | 2.09 | 3.38E-06 | ENSRNOT00000030559 |
| LOC683961-201 | 5 | 196.71 | 2.03 | 1.50E-13 | ENSRNOT00000073017 |
| Ccl21-201 | 5 | 273.11 | 2.14 | 3.88E-20 | ENSRNOT00000023951 |
| Ncbp1-202 | 5 | 18.65 | 7.9 | 1.07E-09 | ENSRNOT00000094126 |
| Abca1-202 | 5 | 5.43 | 2.05 | 0.03 | ENSRNOT00000110113 |
| Ptpn3-201 | 5 | 10.76 | -2.27 | 3.75E-04 | ENSRNOT00000059627 |
| Txn1-201 | 5 | 510.41 | 2.1 | 6.05E-17 | ENSRNOT00000016447 |
| Alad-204 | 5 | 7.79 | 421.05 | 0.02 | ENSRNOT00000077013 |
| LOC100910017-201 | 5 | 71.03 | 2.14 | 2.88E-03 | ENSRNOT00000043610 |
| Tnc-205 | 5 | 32.07 | -2.64 | 2.94E-21 | ENSRNOT00000117346 |
| Tnc-204 | 5 | 13.61 | -3.77 | 3.63E-09 | ENSRNOT00000109041 |
| Rps6-202 | 5 | 2,655.92 | 2.05 | 3.43E-18 | ENSRNOT00000073270 |
| Mysm1-201 | 5 | 13.68 | 2.36 | 6.73E-05 | ENSRNOT00000039554 |
| Itgb3bp-205 | 5 | 12.1 | 3.76 | 0.02 | ENSRNOT00000115426 |
| Itgb3bp-204 | 5 | 17.59 | 2.38 | 0.04 | ENSRNOT00000103428 |
| Itgb3bp-201 | 5 | 20.7 | 4.25 | 3.26E-05 | ENSRNOT00000012947 |
| Magoh-201 | 5 | 202.22 | 2.14 | 3.37E-14 | ENSRNOT00000017422 |
| Txndc12-204 | 5 | 8.17 | 4.81 | 0.04 | ENSRNOT00000102984 |
| Osbpl9-204 | 5 | 5.5 | 361.07 | 0.02 | ENSRNOT00000105115 |
| Faf1-203 | 5 | 14.82 | 2,070.14 | 8.30E-03 | ENSRNOT00000108187 |
| Ppih-201 | 5 | 22.45 | 5.15 | 0.02 | ENSRNOT00000064712 |
| Rps27a-ps1-201 | 5 | 5,213.21 | 2.48 | 6.54E-28 | ENSRNOT00000049535 |
| Ppie-203 | 5 | 5.61 | -369.39 | 0.03 | ENSRNOT00000094173 |
| Macf1-202 | 5 | 6.28 | 10,571.87 | 6.89E-04 | ENSRNOT00000081482 |
| Rpl37a-ps1-201 | 5 | 516.5 | 2.1 | 5.65E-18 | ENSRNOT00000111350 |
| Ythdf2-201 | 5 | 6.95 | -4.39 | 2.67E-03 | ENSRNOT00000014479 |
| Grhl3-202 | 5 | 5.45 | -2.95 | 0.05 | ENSRNOT00000106206 |
| Srsf10-201 | 5 | 50.71 | 2.03 | 8.55E-09 | ENSRNOT00000064925 |
| Rpl11-201 | 5 | 2,602.37 | 2.01 | 2.54E-17 | ENSRNOT00000030043 |
| LOC100359951-201 | 5 | 415.12 | 2.01 | 3.20E-16 | ENSRNOT00000040481 |
| Pramef12-201 | 5 | 21.49 | -2.15 | 0.02 | ENSRNOT00000048232 |
| Rpl35a-201 | 5 | 2,746.95 | 2.45 | 1.52E-27 | ENSRNOT00000042285 |
| Ube4b-202 | 5 | 7 | -3.4 | 1.24E-03 | ENSRNOT00000105982 |
| Eno1-201 | 5 | 74.25 | -2.15 | 1.10E-06 | ENSRNOT00000024106 |
| Rps7-ps23-201 | 5 | 512.57 | 2.16 | 2.03E-22 | ENSRNOT00000011545 |
| Ints11-201 | 5 | 38.87 | -2.07 | 1.23E-06 | ENSRNOT00000026725 |
| Cebpzos-201 | 6 | 17.99 | 2.31 | 0.03 | ENSRNOT00000062031 |
| Mrpl33-202 | 6 | 131.54 | 2.18 | 3.40E-07 | ENSRNOT00000084751 |
| Mrpl33-201 | 6 | 63.23 | 2.25 | 1.34E-05 | ENSRNOT00000034712 |
| Ptrhd1-201 | 6 | 36.14 | 2.09 | 6.54E-04 | ENSRNOT00000005402 |
| Sf3b6-202 | 6 | 201.57 | 2.02 | 4.25E-15 | ENSRNOT00000110429 |
| Cpsf3-201 | 6 | 40.41 | -2.15 | 4.77E-08 | ENSRNOT00000091061 |
| Mboat2-202 | 6 | 6.48 | 4.17 | 0.02 | ENSRNOT00000093903 |
| Rps7-203 | 6 | 3,718.20 | 2.14 | 4.81E-20 | ENSRNOT00000092867 |
| Lamb1-201 | 6 | 29 | -2.41 | 1.08E-07 | ENSRNOT00000008321 |
| Sypl1-203 | 6 | 15.38 | 541.49 | 6.22E-06 | ENSRNOT00000097655 |
| Arl4a-201 | 6 | 6.29 | 2.61 | 4.43E-04 | ENSRNOT00000108440 |

| | | | | | |
|---|---|---|---|---|---|
| Fbxo33-201 | 6 | 6.6 | 2.24 | 0.04 | ENSRNOT00000059316 |
| Fkbp3-201 | 6 | 297.44 | 2.01 | 1.83E-19 | ENSRNOT00000006531 |
| Rpl10l-201 | 6 | 26.62 | -2.46 | 3.53E-05 | ENSRNOT00000117730 |
| Rps29-201 | 6 | 6,545.89 | 2.12 | 6.97E-20 | ENSRNOT00000005577 |
| Psma3-201 | 6 | 168.75 | 2.6 | 7.05E-23 | ENSRNOT00000010753 |
| Mthfd1-204 | 6 | 17.7 | 3.93 | 5.38E-09 | ENSRNOT00000100905 |
| Fntb-201 | 6 | 195.29 | 2.34 | 1.27E-14 | ENSRNOT00000009621 |
| AABR07064980.1-201 | 6 | 329.48 | 2.57 | 4.75E-23 | ENSRNOT00000083272 |
| Rbm25-201 | 6 | 8.87 | 369.91 | 9.63E-05 | ENSRNOT00000003834 |
| Slirp-201 | 6 | 19.86 | 2.53 | 1.07E-10 | ENSRNOT00000016479 |
| Slirp-202 | 6 | 113.32 | 2.27 | 1.55E-07 | ENSRNOT00000110874 |
| Ttc8-206 | 6 | 5.17 | -12.23 | 5.08E-03 | ENSRNOT00000102593 |
| Ttc7b-201 | 6 | 11.88 | -2.96 | 1.88E-03 | ENSRNOT00000037902 |
| Ppp2r5c-203 | 6 | 5.93 | -25.68 | 7.36E-04 | ENSRNOT00000095073 |
| Ppp2r5c-205 | 6 | 15.85 | -2.41 | 1.53E-04 | ENSRNOT00000104738 |
| Klc1-201 | 6 | 6.05 | -4.02 | 0.01 | ENSRNOT00000015935 |
| Zc3h10-202 | 7 | 1,055.45 | 2.22 | 1.49E-22 | ENSRNOT00000105251 |
| LOC102550657-201 | 7 | 8.46 | 2.08 | 6.30E-03 | ENSRNOT00000098807 |
| Tle2-202 | 7 | 12.71 | -2.04 | 0.02 | ENSRNOT00000084006 |
| Map2k2-203 | 7 | 14.68 | -3.4 | 4.36E-03 | ENSRNOT00000102006 |
| Lsm7-201 | 7 | 59.42 | 2.27 | 3.01E-07 | ENSRNOT00000026551 |
| Csrp2-203 | 7 | 7.47 | 2.51 | 1.20E-06 | ENSRNOT00000097951 |
| Tspan8-201 | 7 | 94.78 | 2.13 | 4.79E-06 | ENSRNOT00000005909 |
| Lyz2-201 | 7 | 64.4 | 2.34 | 2.80E-11 | ENSRNOT00000007747 |
| Cox6c-201 | 7 | 1,479.52 | 2.29 | 1.17E-24 | ENSRNOT00000014407 |
| Polr2k-201 | 7 | 97.51 | 2.02 | 1.27E-05 | ENSRNOT00000013869 |
| Eif3e-201 | 7 | 12.55 | 37.09 | 1.02E-07 | ENSRNOT00000038340 |
| Eif3e-203 | 7 | 290.92 | 2.1 | 2.61E-21 | ENSRNOT00000112962 |
| Eny2-202 | 7 | 9.47 | 2.08 | 0.01 | ENSRNOT00000108113 |
| Mal2-201 | 7 | 25.6 | 2.6 | 1.93E-10 | ENSRNOT00000011485 |
| LOC100359421-201 | 7 | 165.06 | 2.61 | 5.39E-13 | ENSRNOT00000049001 |
| Col14a1-201 | 7 | 79.56 | -2.28 | 1.22E-15 | ENSRNOT00000067649 |
| Chrac1-201 | 7 | 51.61 | 2.08 | 4.13E-11 | ENSRNOT00000012121 |
| Gga1-201 | 7 | 15.4 | 2.1 | 9.11E-03 | ENSRNOT00000063876 |
| Cyb5r3-202 | 7 | 22.13 | -2.77 | 2.79E-05 | ENSRNOT00000102247 |
| Fbln1-202 | 7 | 10.9 | -2.02 | 7.88E-03 | ENSRNOT00000088042 |
| Atxn10-202 | 7 | 33.14 | -2.81 | 4.50E-09 | ENSRNOT00000094792 |
| Zbed4-203 | 7 | 8.17 | -2.69 | 7.96E-05 | ENSRNOT00000103004 |
| Zbed4-201 | 7 | 6.87 | 3.2 | 6.50E-05 | ENSRNOT00000006050 |
| Pim3-202 | 7 | 5.73 | 999.55 | 0.03 | ENSRNOT00000085835 |
| Mov10l1-201 | 7 | 70.95 | -2.23 | 5.73E-11 | ENSRNOT00000042686 |
| Rpap3-203 | 7 | 13.49 | -3.72 | 1.27E-03 | ENSRNOT00000111323 |
| Rapgef3-202 | 7 | 6.5 | -9.63 | 4.84E-04 | ENSRNOT00000083182 |
| Rapgef3-203 | 7 | 7.85 | -2.3 | 2.24E-03 | ENSRNOT00000096691 |
| Cacnb3-201 | 7 | 22.7 | -2.11 | 4.52E-04 | ENSRNOT00000081206 |
| Nr4a1-201 | 7 | 18.02 | 2.04 | 0.03 | ENSRNOT00000010171 |
| Znf740-201 | 7 | 10.29 | -2,061.19 | 1.95E-03 | ENSRNOT00000016240 |
| Pfdn5-202 | 7 | 236.06 | 2.03 | 1.42E-07 | ENSRNOT00000109593 |
| Pfdn5-201 | 7 | 361.71 | 2.01 | 1.29E-12 | ENSRNOT00000017794 |
| Slc44a2-203 | 8 | 13.23 | -2.3 | 8.64E-03 | ENSRNOT00000115377 |
| Aplp2-203 | 8 | 40.99 | -2.03 | 7.23E-10 | ENSRNOT00000097481 |

| | | | | | |
|---|---|---|---|---|---|
| Ets1-205 | 8 | 15.05 | -2.03 | 2.41E-06 | ENSRNOT00000111888 |
| Ets1-204 | 8 | 12.83 | 2.73 | 2.04E-03 | ENSRNOT00000102138 |
| Stt3a-201 | 8 | 25.94 | -6.69 | 1.25E-06 | ENSRNOT00000043518 |
| Gramd1b-204 | 8 | 9.24 | 2.12 | 4.28E-03 | ENSRNOT00000099621 |
| Tbcel-201 | 8 | 5 | -4.65 | 0.05 | ENSRNOT00000059951 |
| Rps25-201 | 8 | 429.82 | 2.45 | 1.88E-27 | ENSRNOT00000029546 |
| Rps25-202 | 8 | 564.07 | 2.46 | 1.52E-27 | ENSRNOT00000104255 |
| Atp5mg-201 | 8 | 339.61 | 2.34 | 4.29E-27 | ENSRNOT00000050878 |
| Usp28-202 | 8 | 10.7 | -2.26 | 0.01 | ENSRNOT00000067420 |
| Il18-202 | 8 | 13.73 | 2.58 | 0.03 | ENSRNOT00000108779 |
| Timm8b-201 | 8 | 435.69 | 2.28 | 5.28E-26 | ENSRNOT00000013188 |
| Fdx1-201 | 8 | 124.6 | 2.68 | 1.83E-17 | ENSRNOT00000016263 |
| Cyp19a1-201 | 8 | 66.46 | 3.07 | 7.74E-09 | ENSRNOT00000000212 |
| Acsbg1-204 | 8 | 17.74 | 2.64 | 6.41E-03 | ENSRNOT00000119621 |
| Parp6-203 | 8 | 16.2 | -2.61 | 1.14E-04 | ENSRNOT00000105803 |
| Rplp1-201 | 8 | 3,207.32 | 2.37 | 4.00E-26 | ENSRNOT00000018820 |
| Cilp-201 | 8 | 12.51 | -2.12 | 2.33E-05 | ENSRNOT00000044887 |
| Rps27l-201 | 8 | 268.04 | 2.3 | 9.27E-29 | ENSRNOT00000090977 |
| Eef1a1-202 | 8 | 16.44 | -2.36 | 0.02 | ENSRNOT00000112753 |
| Morf4l1-201 | 8 | 30.77 | 3,025.94 | 3.96E-04 | ENSRNOT00000080421 |
| U2surp-203 | 8 | 18.25 | -2.04 | 2.00E-05 | ENSRNOT00000116411 |
| Slc25a36-202 | 8 | 19.04 | -1,863.43 | 5.83E-04 | ENSRNOT00000110158 |
| Tma7-201 | 8 | 138.32 | 2.04 | 1.91E-07 | ENSRNOT00000075819 |
| Setd2-201 | 8 | 6.07 | -165.63 | 1.59E-10 | ENSRNOT00000028409 |
| LOC100360828-201 | 8 | 108.11 | 2.02 | 9.59E-06 | ENSRNOT00000030221 |
| LOC102555317-202 | 8 | 12.18 | 2.36 | 6.59E-04 | ENSRNOT00000116868 |
| Rps20-ps11-201 | 9 | 13.9 | 64.33 | 5.27E-04 | ENSRNOT00000051449 |
| Gsta1-201 | 9 | 312.45 | 2.32 | 6.97E-18 | ENSRNOT00000018325 |
| Dst-203 | 9 | 7.71 | -6.85 | 3.37E-12 | ENSRNOT00000095842 |
| Rpl31-201 | 9 | 2,080.90 | 2.01 | 1.23E-14 | ENSRNOT00000033733 |
| Ecrg4-202 | 9 | 24.43 | 3.58 | 4.76E-03 | ENSRNOT00000095751 |
| Col5a2-201 | 9 | 63.91 | -2.02 | 6.21E-11 | ENSRNOT00000005073 |
| Gls-203 | 9 | 31.27 | 2.3 | 1.11E-08 | ENSRNOT00000110936 |
| Hspe1-202 | 9 | 215.06 | 2 | 1.68E-13 | ENSRNOT00000102977 |
| Hspe1-201 | 9 | 459.82 | 2.4 | 7.38E-30 | ENSRNOT00000082300 |
| Mob4-203 | 9 | 33.41 | 2.05 | 1.63E-06 | ENSRNOT00000103898 |
| ENSRNOT00000049571 | 9 | 1,010.40 | 2.5 | 9.42E-30 | ENSRNOT00000049571 |
| Zdbf2-201 | 9 | 5.84 | 2 | 1.09E-03 | ENSRNOT00000016038 |
| Atic-203 | 9 | 29.66 | -2.03 | 0.05 | ENSRNOT00000116844 |
| Fn1-201 | 9 | 5.55 | -116.87 | 2.13E-17 | ENSRNOT00000019772 |
| ENSRNOT00000098542 | 9 | 583.48 | 2.19 | 5.29E-22 | ENSRNOT00000098542 |
| Ptprn-201 | 9 | 5.74 | 2.47 | 0.03 | ENSRNOT00000026654 |
| Kcne4-201 | 9 | 18.12 | 2.35 | 3.43E-06 | ENSRNOT00000111796 |
| ENSRNOT00000110318 | 9 | 25.89 | 3.12 | 7.41E-12 | ENSRNOT00000110318 |
| Ppp4r1-202 | 9 | 5.87 | 245.83 | 4.18E-07 | ENSRNOT00000104139 |
| Ndufv2-202 | 9 | 26.04 | 2.17 | 0.05 | ENSRNOT00000106172 |
| Washc1-203 | 9 | 8.56 | 13.03 | 3.52E-05 | ENSRNOT00000104558 |
| Tgif1-202 | 9 | 5.79 | -117.71 | 4.13E-04 | ENSRNOT00000102373 |
| LOC120094815-208 | 9 | 10.2 | 2.29 | 2.64E-03 | ENSRNOT00000113468 |
| ENSRNOT00000101043 | 9 | 15.06 | 2.29 | 8.33E-03 | ENSRNOT00000101043 |
| Abcc1-202 | 10 | 5.46 | 4.44 | 0.03 | ENSRNOT00000103910 |

| | | | | | |
|---|---|---|---|---|---|
| Glyr1-202 | 10 | 26.89 | -2.74 | 1.56E-07 | ENSRNOT00000095486 |
| Mgrn1-201 | 10 | 17.01 | -2.61 | 5.27E-10 | ENSRNOT00000004416 |
| Srrm2-201 | 10 | 23.82 | 14.08 | 2.49E-07 | ENSRNOT00000091287 |
| Rnps1-204 | 10 | 25.68 | -2.4 | 3.16E-05 | ENSRNOT00000105026 |
| Tsc2-201 | 10 | 18.8 | -2.24 | 2.44E-09 | ENSRNOT00000016221 |
| Snrpc-ps3-201 | 10 | 27.15 | 5.39 | 9.23E-06 | ENSRNOT00000023626 |
| Mcrip2-201 | 10 | 32.06 | 2.16 | 0.01 | ENSRNOT00000027170 |
| Hba-a1-205 | 10 | 68.5 | -2.48 | 3.24E-03 | ENSRNOT00000119140 |
| Hba-a1-201 | 10 | 3,133.04 | -2.3 | 4.96E-05 | ENSRNOT00000051483 |
| Hba-a1-202 | 10 | 2,915.38 | -2.41 | 9.43E-06 | ENSRNOT00000052292 |
| Hba-a1-206 | 10 | 2,389.82 | -2.31 | 3.33E-06 | ENSRNOT00000119443 |
| Fbxw11-203 | 10 | 8.04 | -124.44 | 5.70E-12 | ENSRNOT00000098062 |
| Havcr1-201 | 10 | 15.35 | 2.35 | 0.02 | ENSRNOT00000009573 |
| Zfp2-202 | 10 | 6.11 | 2.08 | 0.04 | ENSRNOT00000102165 |
| Rpl30l2-201 | 10 | 3,158.69 | 2.47 | 3.23E-28 | ENSRNOT00000042688 |
| Uqcrq-201 | 10 | 523.12 | 2.04 | 9.33E-21 | ENSRNOT00000100770 |
| Anxa6-202 | 10 | 18.76 | -2.64 | 1.10E-05 | ENSRNOT00000096029 |
| Srebf1-201 | 10 | 10.53 | -2 | 1.62E-03 | ENSRNOT00000004753 |
| Pmp22-201 | 10 | 21.24 | 2,413.27 | 6.83E-03 | ENSRNOT00000041606 |
| Ndel1-202 | 10 | 14.95 | -3.63 | 2.65E-05 | ENSRNOT00000065505 |
| Rpl26-201 | 10 | 3,504.31 | 2.45 | 2.93E-26 | ENSRNOT00000005588 |
| Eif4a1-203 | 10 | 108.11 | 2.87 | 6.15E-13 | ENSRNOT00000116199 |
| Tmem256-201 | 10 | 125.15 | 2.11 | 2.37E-07 | ENSRNOT00000056870 |
| Alox15-201 | 10 | 133.56 | -2.48 | 2.73E-05 | ENSRNOT00000026038 |
| Rpl36al-201 | 10 | 1,820.92 | 2.58 | 2.34E-22 | ENSRNOT00000043171 |
| P2rx1-203 | 10 | 5.56 | -4.98 | 0.05 | ENSRNOT00000118157 |
| Rpl37l1-201 | 10 | 4,281.86 | 2.42 | 1.88E-27 | ENSRNOT00000046519 |
| Blmh-201 | 10 | 17.3 | 2.08 | 6.25E-03 | ENSRNOT00000005125 |
| Nufip2-201 | 10 | 6.79 | -15.35 | 2.90E-06 | ENSRNOT00000030501 |
| Rpl23a-202 | 10 | 837.64 | 2.01 | 5.51E-17 | ENSRNOT00000114672 |
| Nf1-201 | 10 | 5.16 | -3.18 | 2.60E-04 | ENSRNOT00000046262 |
| Ap2b1-203 | 10 | 7.6 | -18.02 | 1.96E-03 | ENSRNOT00000090446 |
| Bcas3-207 | 10 | 9.54 | -2.71 | 1.49E-04 | ENSRNOT00000119090 |
| Cltc-203 | 10 | 9.71 | -2.52 | 1.07E-03 | ENSRNOT00000098805 |
| Ypel2-202 | 10 | 9.52 | -22.36 | 1.52E-08 | ENSRNOT00000114367 |
| Elob-ps4-201 | 10 | 509.23 | 2.15 | 1.86E-11 | ENSRNOT00000095697 |
| Mmd-201 | 10 | 14.13 | 3.54 | 0.02 | ENSRNOT00000003308 |
| LOC100363469-201 | 10 | 299.93 | 3.19 | 2.19E-35 | ENSRNOT00000043005 |
| Utp18-202 | 10 | 12.46 | 2.16 | 0.03 | ENSRNOT00000111110 |
| Tob1-201 | 10 | 16.87 | -2.2 | 6.69E-03 | ENSRNOT00000003780 |
| Luc7l3-202 | 10 | 21.26 | 2.44 | 5.41E-06 | ENSRNOT00000101307 |
| Luc7l3-204 | 10 | 7.25 | 9.21 | 2.95E-04 | ENSRNOT00000111455 |
| Kat7-204 | 10 | 6.24 | -281.17 | 5.36E-06 | ENSRNOT00000109616 |
| Rpl23-201 | 10 | 1,362.80 | 2.17 | 1.01E-20 | ENSRNOT00000005471 |
| Med24-202 | 10 | 30.37 | -2.12 | 4.01E-11 | ENSRNOT00000100598 |
| Msl1-201 | 10 | 12.52 | 2.73 | 1.34E-04 | ENSRNOT00000012812 |
| Msl1-202 | 10 | 6.37 | -200.85 | 2.55E-05 | ENSRNOT00000101887 |
| Mlx-204 | 10 | 5.05 | -486.77 | 0.01 | ENSRNOT00000119770 |
| G6pc1-204 | 10 | 9.62 | -2.38 | 0.01 | ENSRNOT00000076820 |
| Arl4d-204 | 10 | 11.52 | 2.62 | 0.02 | ENSRNOT00000110523 |
| Slc4a1-201 | 10 | 6.55 | -3.79 | 1.90E-03 | ENSRNOT00000028445 |

| | | | | | |
|---|---|---|---|---|---|
| Nsf-201 | 10 | 24.55 | -2.7 | 1.75E-07 | ENSRNOT00000006361 |
| Ace-202 | 10 | 10.53 | 3.06 | 9.41E-04 | ENSRNOT00000092961 |
| Ddx5-204 | 10 | 72.36 | -2.45 | 2.52E-05 | ENSRNOT00000116552 |
| Bptf-202 | 10 | 5.38 | -3.12 | 7.18E-06 | ENSRNOT00000098328 |
| RGD1359290-201 | 10 | 51.34 | 2.46 | 6.92E-04 | ENSRNOT00000029782 |
| Rpl38-ps9-203 | 10 | 2,367.15 | 2.24 | 6.92E-24 | ENSRNOT00000095224 |
| Rps18l1-201 | 10 | 3,153.66 | 2.1 | 1.05E-19 | ENSRNOT00000046010 |
| Actg1-201 | 10 | 340.78 | -4.33 | 7.49E-39 | ENSRNOT00000054976 |
| Nploc4-201 | 10 | 25.62 | -2.36 | 4.62E-05 | ENSRNOT00000054973 |
| Foxk2-203 | 10 | 8.64 | 5.45 | 0.02 | ENSRNOT00000099287 |
| Nrip1-208 | 11 | 11.87 | 2.95 | 1.14E-07 | ENSRNOT00000114589 |
| Kcne2-204 | 11 | 36.11 | 2.28 | 7.05E-04 | ENSRNOT00000099713 |
| Cbr1l2-205 | 11 | 8.47 | -133.03 | 9.98E-04 | ENSRNOT00000085961 |
| Nit2-201 | 11 | 15.61 | 2.45 | 0.02 | ENSRNOT00000029420 |
| H3f3a-ps9-201 | 11 | 103.18 | 2.33 | 1.07E-10 | ENSRNOT00000110622 |
| Golgb1-204 | 11 | 6.79 | -2.48 | 0.04 | ENSRNOT00000110978 |
| Kpna1-203 | 11 | 6.48 | 2.14 | 3.65E-03 | ENSRNOT00000097087 |
| Pdia5-201 | 11 | 13.1 | -3.38 | 0.01 | ENSRNOT00000067984 |
| Tnk2-203 | 11 | 9.25 | -4.04 | 3.21E-06 | ENSRNOT00000085568 |
| Tnk2-205 | 11 | 13.88 | -2.12 | 6.47E-04 | ENSRNOT00000116648 |
| Opa1-205 | 11 | 5.35 | -5.13 | 0.03 | ENSRNOT00000117359 |
| Ccdc50-203 | 11 | 9.51 | 2.27 | 0.02 | ENSRNOT00000112380 |
| LOC100365839-201 | 11 | 103.73 | 2.1 | 1.14E-12 | ENSRNOT00000074617 |
| Adipoq-201 | 11 | 8.87 | 4.37 | 0.01 | ENSRNOT00000002492 |
| RGD1559972-201 | 11 | 1,411.90 | 2.15 | 1.92E-21 | ENSRNOT00000041326 |
| Alg3-201 | 11 | 10.03 | -5.37 | 0.01 | ENSRNOT00000002331 |
| Dgcr2-203 | 11 | 10.88 | 7.43 | 4.22E-06 | ENSRNOT00000107429 |
| Xab2-201 | 12 | 13.73 | -2.1 | 1.61E-03 | ENSRNOT00000001309 |
| Pomp-204 | 12 | 72.28 | 2.61 | 6.97E-18 | ENSRNOT00000118745 |
| Pomp-201 | 12 | 473.6 | 2.13 | 8.47E-19 | ENSRNOT00000071796 |
| Rpl21-202 | 12 | 531.81 | 2.18 | 5.24E-17 | ENSRNOT00000099991 |
| Rpl21-201 | 12 | 890.57 | 2.36 | 2.81E-27 | ENSRNOT00000001265 |
| Atp5mf-202 | 12 | 514.71 | 2.14 | 7.39E-21 | ENSRNOT00000104742 |
| Actb-204 | 12 | 5.6 | 637.43 | 9.66E-03 | ENSRNOT00000116486 |
| Zfp68-202 | 12 | 7.99 | 3.51 | 1.57E-06 | ENSRNOT00000079221 |
| Rpl31l4-201 | 12 | 516.05 | 3.73 | 9.21E-23 | ENSRNOT00000045401 |
| Mcm7-202 | 12 | 28.66 | -2.38 | 5.87E-07 | ENSRNOT00000105826 |
| Zp3-201 | 12 | 936.49 | -2.1 | 2.73E-18 | ENSRNOT00000001952 |
| Mdh2-202 | 12 | 28.28 | 2,361.01 | 6.82E-04 | ENSRNOT00000112030 |
| Clip1-206 | 12 | 6.01 | -504.29 | 1.10E-06 | ENSRNOT00000098458 |
| Pxn-201 | 12 | 9.83 | 5.76 | 5.45E-04 | ENSRNOT00000047466 |
| Pxn-203 | 12 | 11.86 | -8.14 | 1.87E-04 | ENSRNOT00000081134 |
| Relch-204 | 13 | 5.25 | 3.08 | 2.50E-03 | ENSRNOT00000113641 |
| LOC108348870-201 | 13 | 727.97 | 2.1 | 6.02E-16 | ENSRNOT00000046791 |
| Ptpn4-202 | 13 | 13.31 | 2.39 | 4.56E-13 | ENSRNOT00000095537 |
| Dbi-202 | 13 | 658.44 | 2.27 | 4.37E-19 | ENSRNOT00000074685 |
| Dbi-205 | 13 | 99.91 | 2.21 | 1.49E-12 | ENSRNOT00000117885 |
| Steap3-204 | 13 | 8.99 | 3.77 | 0.01 | ENSRNOT00000106336 |
| Steap3-202 | 13 | 8.45 | -280.14 | 1.96E-05 | ENSRNOT00000101486 |
| Actr3-201 | 13 | 29.64 | 4,907.80 | 3.76E-03 | ENSRNOT00000004520 |
| Rab3gap1-201 | 13 | 7.01 | 2.1 | 0.02 | ENSRNOT00000005289 |

| | | | | | |
|---|---|---|---|---|---|
| Atp2b4-204 | 13 | 5.9 | 79.52 | 1.45E-24 | ENSRNOT00000112237 |
| Camsap2-203 | 13 | 7.12 | 9.54 | 9.83E-09 | ENSRNOT00000098757 |
| Camsap2-201 | 13 | 10.38 | -3.03 | 9.91E-05 | ENSRNOT00000060410 |
| Xpr1-201 | 13 | 10.1 | -11.44 | 4.79E-09 | ENSRNOT00000000049 |
| Xpr1-205 | 13 | 6.34 | 7.36 | 4.34E-03 | ENSRNOT00000113923 |
| Tdrd5-201 | 13 | 80 | -2.03 | 2.02E-09 | ENSRNOT00000005353 |
| Kifap3-202 | 13 | 17.68 | -2 | 6.98E-04 | ENSRNOT00000085135 |
| Ildr2-204 | 13 | 6.62 | 3.23 | 9.87E-05 | ENSRNOT00000072897 |
| Copa-204 | 13 | 24.59 | -2.98 | 5.36E-11 | ENSRNOT00000113779 |
| LOC108348980-201 | 13 | 29.55 | 3.54 | 0.03 | ENSRNOT00000041384 |
| Cdc42bpa-206 | 13 | 12.19 | 3.62 | 0.04 | ENSRNOT00000116713 |
| Tlr5-203 | 13 | 14.17 | -2.18 | 1.11E-03 | ENSRNOT00000107238 |
| Tlr5-201 | 13 | 14.89 | -2.27 | 3.14E-04 | ENSRNOT00000004821 |
| Eprs1-202 | 13 | 36.49 | -2.1 | 4.29E-04 | ENSRNOT00000085214 |
| Tmed5-201 | 14 | 8.08 | 92.6 | 2.15E-04 | ENSRNOT00000000083 |
| Lrrc8d-205 | 14 | 6.45 | -1,313.96 | 2.62E-03 | ENSRNOT00000109682 |
| Mrps18c-203 | 14 | 56.25 | 2.7 | 1.80E-07 | ENSRNOT00000110810 |
| Sec31a-201 | 14 | 12.13 | -2.15 | 7.98E-04 | ENSRNOT00000003072 |
| Rpl9-ps30-201 | 14 | 63.84 | 2.01 | 1.36E-03 | ENSRNOT00000038927 |
| AABR07015346.1-201 | 14 | 783.45 | 2.12 | 2.82E-21 | ENSRNOT00000038497 |
| Wfs1-201 | 14 | 28.27 | -2.22 | 3.45E-10 | ENSRNOT00000034730 |
| Add1-202 | 14 | 22.7 | -2.58 | 2.36E-06 | ENSRNOT00000018340 |
| Spon2-201 | 14 | 17.63 | -2.01 | 8.73E-03 | ENSRNOT00000008262 |
| Morc2-201 | 14 | 7.91 | -6.36 | 7.79E-10 | ENSRNOT00000026569 |
| Zmiz2-202 | 14 | 5.4 | -3.41 | 0.03 | ENSRNOT00000077428 |
| Sec61g-201 | 14 | 726.71 | 2.59 | 1.66E-33 | ENSRNOT00000006906 |
| Rps27a-201 | 14 | 46.69 | 2.26 | 2.41E-06 | ENSRNOT00000005872 |
| ENSRNOT00000106404 | 14 | 57.33 | 2.69 | 0.01 | ENSRNOT00000106404 |
| Chchd1-201 | 15 | 193.47 | 2.47 | 5.44E-20 | ENSRNOT00000012372 |
| Mrps16-201 | 15 | 11.96 | 153.13 | 4.43E-06 | ENSRNOT00000009207 |
| Slc4a7-206 | 15 | 5.92 | 2.01 | 1.39E-04 | ENSRNOT00000118797 |
| Mcpt1l1-201 | 15 | 32.56 | -2.09 | 0.01 | ENSRNOT00000090206 |
| Clu-202 | 15 | 70.5 | -2.02 | 4.37E-08 | ENSRNOT00000078861 |
| Kctd9-202 | 15 | 6.3 | 2.64 | 0.04 | ENSRNOT00000105780 |
| Dpm3-201 | 15 | 159.38 | 2.15 | 7.34E-07 | ENSRNOT00000045477 |
| Itm2b-202 | 15 | 6.62 | 2.7 | 1.82E-03 | ENSRNOT00000106965 |
| Tpt1-201 | 15 | 7,454.15 | 2.06 | 6.95E-19 | ENSRNOT00000001383 |
| Dnajc15-201 | 15 | 104.9 | 2.13 | 5.24E-13 | ENSRNOT00000012530 |
| Rps24-201 | 16 | 5,804.32 | 2.47 | 1.24E-28 | ENSRNOT00000013588 |
| Rps24-203 | 16 | 244.04 | 2.19 | 4.58E-11 | ENSRNOT00000101331 |
| Rps24-205 | 16 | 38.97 | 2.07 | 0.03 | ENSRNOT00000118714 |
| Uqcc5-202 | 16 | 75.89 | 2.26 | 1.77E-07 | ENSRNOT00000116930 |
| Mettl6-204 | 16 | 15 | -8.05 | 6.29E-10 | ENSRNOT00000103715 |
| Tma16-201 | 16 | 28.59 | 2.02 | 6.01E-03 | ENSRNOT00000019363 |
| Hmgb2-201 | 16 | 158.93 | -2.57 | 1.87E-06 | ENSRNOT00000017635 |
| LOC103694242-201 | 16 | 399.72 | 2.14 | 4.63E-23 | ENSRNOT00000009702 |
| Hpgd-201 | 16 | 15.71 | 2.27 | 0.02 | ENSRNOT00000014229 |
| Gpm6a-204 | 16 | 5.34 | 3.95 | 0.02 | ENSRNOT00000100181 |
| Fat1-203 | 16 | 19.35 | -2.4 | 2.12E-06 | ENSRNOT00000095529 |
| Fat1-201 | 16 | 16.49 | 2.43 | 1.19E-04 | ENSRNOT00000067486 |
| Pcm1-201 | 16 | 9.99 | -2.51 | 3.21E-04 | ENSRNOT00000014202 |

| | | | | | |
|---|---|---|---|---|---|
| Mtmr7-204 | 16 | 13.25 | -2.8 | 3.34E-03 | ENSRNOT00000105685 |
| Star-201 | 16 | 103.61 | 2.09 | 5.70E-12 | ENSRNOT00000020606 |
| ENSRNOT00000102571 | 17 | 5.71 | 2.16 | 2.22E-03 | ENSRNOT00000102571 |
| LOC100361933-201 | 17 | 2,488.05 | 2.19 | 5.64E-22 | ENSRNOT00000093460 |
| Phf2-201 | 17 | 5.13 | 51.35 | 3.54E-12 | ENSRNOT00000022669 |
| Phactr1-203 | 17 | 6 | -2.47 | 0.03 | ENSRNOT00000107421 |
| Ssr1-204 | 17 | 11.14 | -4.32 | 0.03 | ENSRNOT00000089736 |
| Fars2-205 | 17 | 11.84 | -2.57 | 0.03 | ENSRNOT00000113971 |
| Serpinb6a-201 | 17 | 30.76 | 2.05 | 3.98E-04 | ENSRNOT00000024161 |
| Serpinb9-204 | 17 | 11.89 | 1,804.43 | 0.01 | ENSRNOT00000115382 |
| Carmil1-204 | 17 | 7.02 | -30.69 | 5.01E-11 | ENSRNOT00000108358 |
| Zfp322a-201 | 17 | 9.55 | 2.12 | 5.63E-05 | ENSRNOT00000023268 |
| Sfrp4-202 | 17 | 20.76 | 2.44 | 3.53E-04 | ENSRNOT00000079368 |
| Stard3nl-202 | 17 | 32.98 | 2.7 | 1.25E-03 | ENSRNOT00000097142 |
| Crem-204 | 17 | 16.43 | 2.04 | 5.66E-03 | ENSRNOT00000068545 |
| Lgals8-202 | 17 | 7.08 | 2.49 | 0.02 | ENSRNOT00000107534 |
| Gdi2-202 | 17 | 94.31 | 3.86 | 2.33E-31 | ENSRNOT00000104020 |
| Rpl39-201 | 18 | 4,302.18 | 2.84 | 1.08E-38 | ENSRNOT00000043244 |
| Cdh2-201 | 18 | 16.6 | -2.24 | 8.54E-05 | ENSRNOT00000021170 |
| Fhod3-204 | 18 | 11.96 | -2.53 | 1.62E-04 | ENSRNOT00000093285 |
| Hspa9-202 | 18 | 47.32 | 312.08 | 9.12E-40 | ENSRNOT00000079460 |
| Pfdn1-202 | 18 | 61.61 | 2.71 | 7.44E-06 | ENSRNOT00000099632 |
| Dpysl3-202 | 18 | 12.57 | -2.42 | 3.64E-03 | ENSRNOT00000087876 |
| LOC100362479-201 | 18 | 16.32 | 4.31 | 0.02 | ENSRNOT00000109265 |
| Rps14-201 | 18 | 2,982.16 | 2.33 | 1.27E-23 | ENSRNOT00000059501 |
| Tcof1-203 | 18 | 8.76 | 3,271.29 | 3.53E-04 | ENSRNOT00000115228 |
| Tcof1-201 | 18 | 20.52 | -5.35 | 2.96E-17 | ENSRNOT00000045041 |
| Csnk1a1-203 | 18 | 20.39 | 3.05 | 1.80E-18 | ENSRNOT00000103047 |
| Txnl1-204 | 18 | 19.99 | 238.58 | 4.69E-06 | ENSRNOT00000100860 |
| Cycs-201 | 18 | 133.54 | 2.8 | 4.75E-20 | ENSRNOT00000102631 |
| Cyb5a-207 | 18 | 26.3 | 2 | 3.41E-03 | ENSRNOT00000112034 |
| Katnb1-201 | 19 | 14.67 | -2.05 | 0.01 | ENSRNOT00000019770 |
| Adgrg1-201 | 19 | 10.71 | -2.06 | 0.03 | ENSRNOT00000020921 |
| Adgre5-201 | 19 | 20.77 | -2.55 | 7.06E-07 | ENSRNOT00000006152 |
| Arhgap10-203 | 19 | 5.27 | -7.38 | 9.44E-03 | ENSRNOT00000104023 |
| LOC100360573-201 | 19 | 31.83 | 4.34 | 1.15E-03 | ENSRNOT00000027225 |
| Cyb5b-202 | 19 | 6.1 | 14.49 | 5.12E-03 | ENSRNOT00000086435 |
| Calb2-201 | 19 | 9.68 | 2.79 | 0.02 | ENSRNOT00000022943 |
| Ctrb1-201 | 19 | 23.35 | -2.76 | 0.02 | ENSRNOT00000026017 |
| Hsd17b2-201 | 19 | 17.58 | 2.47 | 0.02 | ENSRNOT00000018795 |
| RT1-N2-201 | 20 | 14.56 | -2.8 | 4.99E-03 | ENSRNOT00000050159 |
| Col11a2-202 | 20 | 5.58 | -2.2 | 0.03 | ENSRNOT00000084117 |
| Tead3-203 | 20 | 13.87 | -2.71 | 0.02 | ENSRNOT00000107495 |
| Clps-201 | 20 | 16.26 | 4.28 | 0.03 | ENSRNOT00000000611 |
| Kctd20-203 | 20 | 5.38 | 3.95 | 2.82E-04 | ENSRNOT00000105827 |
| RGD735065-203 | 20 | 7.04 | -77.13 | 6.52E-04 | ENSRNOT00000093188 |
| Mcm3ap-201 | 20 | 5.76 | -2.03 | 0.02 | ENSRNOT00000064637 |
| Susd2-202 | 20 | 6.64 | -2.18 | 0.05 | ENSRNOT00000079996 |
| Hnrnph3-204 | 20 | 28.27 | 2.48 | 5.24E-17 | ENSRNOT00000105550 |
| Oit3-202 | 20 | 9.13 | 37.08 | 1.74E-09 | ENSRNOT00000112351 |
| Oit3-201 | 20 | 49.74 | -2.27 | 4.87E-11 | ENSRNOT00000075682 |

| | | | | | |
|---|---|---|---|---|---|
| Psap-203 | 20 | 112.07 | -2.12 | 1.02E-14 | ENSRNOT00000104593 |
| Ppa1-207 | 20 | 41.67 | 2.5 | 8.91E-07 | ENSRNOT00000107042 |
| Man1a1-201 | 20 | 13.28 | 2.1 | 3.18E-03 | ENSRNOT00000057422 |
| Crybg1-201 | 20 | 29.1 | -3.02 | 2.01E-24 | ENSRNOT00000036753 |
| Uba1-202 | X | 162.08 | -2.08 | 3.35E-15 | ENSRNOT00000085338 |
| Hdac6-203 | X | 13.57 | -2.58 | 3.06E-03 | ENSRNOT00000116471 |
| Spin2b-202 | X | 87.09 | 2.16 | 2.07E-11 | ENSRNOT00000113757 |
| Phf8-202 | X | 7.33 | -2.09 | 2.87E-03 | ENSRNOT00000093422 |
| Tmsb4x-201 | X | 5,488.67 | 2.15 | 4.75E-20 | ENSRNOT00000071708 |
| Acot9-202 | X | 20.33 | 2.06 | 3.44E-03 | ENSRNOT00000111580 |
| Ogt-203 | X | 44.21 | -5.95 | 7.85E-07 | ENSRNOT00000100805 |
| Cox7b-201 | X | 713.89 | 2.13 | 4.98E-20 | ENSRNOT00000090007 |
| Zfp711-202 | X | 5.48 | 4.09 | 0.02 | ENSRNOT00000095292 |
| Rpl36a-201 | X | 981.09 | 2.06 | 1.42E-12 | ENSRNOT00000057222 |
| Tceal9-202 | X | 494.5 | 2 | 7.28E-13 | ENSRNOT00000110789 |
| Tmsb15b2-201 | X | 171.04 | 2.7 | 1.12E-22 | ENSRNOT00000057140 |
| Zcchc18-201 | X | 31.53 | 2.43 | 2.51E-13 | ENSRNOT00000092013 |
| LOC100361854-201 | X | 938.75 | 2.19 | 8.33E-23 | ENSRNOT00000051134 |
| Il13ra1-ps1-202 | X | 8.41 | 166.04 | 1.42E-12 | ENSRNOT00000112175 |
| Rpl39-ps13-201 | X | 24.72 | 2.21 | 0.04 | ENSRNOT00000092456 |
| Ints6l-201 | X | 22.34 | 2.04 | 6.89E-06 | ENSRNOT00000071888 |
| RGD1559982-201 | X | 8.47 | -2.61 | 0.03 | ENSRNOT00000096951 |
| Bgn-201 | X | 175.92 | -2.24 | 7.00E-20 | ENSRNOT00000085201 |
| Fam3a-203 | X | 12.14 | -3.56 | 0.05 | ENSRNOT00000096091 |
| Mt-nd3-201 | MT | 7,921.34 | 2.03 | 2.44E-15 | ENSRNOT00000041241 |

**Supplementary Table A3-2.** Differentially expressed downstream genes in PD8.5 wildtype ovaries compared to PD 6.5 wildtype ovaries.

| Name | Chrom | Max group mean | Fold change | FDR p-value | ENSEMBL |
|---|---|---|---|---|---|
| Vom2r6-202 | 1 | 6.08 | -2.09 | 7.45E-03 | ENSRNOT00000044187 |
| Rrlt-201 | 1 | 15.14 | -2.28 | 0.02 | ENSRNOT00000101208 |
| Sash1-203 | 1 | 6.09 | 2.12 | 0.01 | ENSRNOT00000098748 |
| Sash1-206 | 1 | 5.11 | -3.74 | 2.89E-03 | ENSRNOT00000107987 |
| Lama2-201 | 1 | 22.43 | 2.09 | 6.62E-21 | ENSRNOT00000014917 |
| Arhgap18-202 | 1 | 10.33 | -2.35 | 2.98E-03 | ENSRNOT00000096976 |
| LOC100360791-201 | 1 | 30.74 | -4.4 | 4.83E-03 | ENSRNOT00000044222 |
| Akap7-202 | 1 | 7.55 | 2.96 | 3.35E-03 | ENSRNOT00000096819 |
| Sgk1-201 | 1 | 47.02 | 2.42 | 1.92E-17 | ENSRNOT00000016121 |
| Sgk1-203 | 1 | 18.99 | 2.23 | 3.80E-04 | ENSRNOT00000061157 |
| Sgk1-202 | 1 | 64.08 | 2.08 | 2.67E-10 | ENSRNOT00000040736 |
| Srd5a1-201 | 1 | 27.98 | 2.93 | 6.33E-07 | ENSRNOT00000023659 |
| ENSRNOT00000116757 | 1 | 20.59 | -2.68 | 4.73E-42 | ENSRNOT00000116757 |
| Scaf8-201 | 1 | 14.06 | 2.82 | 7.65E-05 | ENSRNOT00000065733 |
| Ezr-203 | 1 | 21.39 | -9.07 | 1.88E-21 | ENSRNOT00000105371 |
| Mpc1-202 | 1 | 5.2 | -2.18 | 0.01 | ENSRNOT00000098843 |
| Rps6ka2-201 | 1 | 5.51 | 2.12 | 0.02 | ENSRNOT00000017809 |
| Smoc2-201 | 1 | 59.64 | 2.91 | 2.16E-29 | ENSRNOT00000019317 |
| Smoc2-202 | 1 | 65.99 | 2.64 | 2.66E-32 | ENSRNOT00000079718 |
| LOC102548728-201 | 1 | 5.86 | -2.35 | 0.01 | ENSRNOT00000112121 |
| LOC682225-201 | 1 | 8.54 | -2 | 0.05 | ENSRNOT00000033187 |
| Peg3-202 | 1 | 11.32 | -2.77 | 8.08E-15 | ENSRNOT00000106399 |
| Tnni3-201 | 1 | 21.85 | -2.08 | 0.03 | ENSRNOT00000024640 |
| Tnnt1-201 | 1 | 5.6 | -61.4 | 9.94E-04 | ENSRNOT00000034957 |
| Nlrp2-201 | 1 | 12.55 | 2.97 | 4.48E-09 | ENSRNOT00000073464 |
| Nlrp4b-201 | 1 | 7.65 | 2.6 | 5.79E-03 | ENSRNOT00000035435 |
| Nlrp4b-202 | 1 | 6.73 | 3.16 | 3.12E-03 | ENSRNOT00000041128 |
| Rtn2-201 | 1 | 10.04 | 3.12 | 6.53E-04 | ENSRNOT00000022528 |
| Apoc1-201 | 1 | 240.52 | -8.11 | 1.67E-68 | ENSRNOT00000046169 |
| Apoe-205 | 1 | 31.05 | -2.2 | 1.98E-04 | ENSRNOT00000108411 |
| Apoe-204 | 1 | 35.34 | -2.3 | 1.23E-10 | ENSRNOT00000099048 |
| Apoe-202 | 1 | 1,131.37 | -2.31 | 3.85E-61 | ENSRNOT00000080453 |
| Zfp180-202 | 1 | 6.16 | -3.91 | 0.02 | ENSRNOT00000111936 |
| Phldb3-201 | 1 | 7.86 | 2.23 | 0.03 | ENSRNOT00000083493 |
| Tex101-201 | 1 | 6.43 | -6.72 | 2.18E-03 | ENSRNOT00000027202 |
| Lipe-202 | 1 | 6.4 | 2.13 | 0.05 | ENSRNOT00000099694 |
| Mia-201 | 1 | 29.26 | -2.42 | 0.01 | ENSRNOT00000002050 |
| Lgals7-201 | 1 | 9.14 | -4.63 | 0.04 | ENSRNOT00000027620 |
| LOC100361913-201 | 1 | 117.86 | -2.31 | 1.41E-06 | ENSRNOT00000027998 |
| Spint2-201 | 1 | 12.36 | 919.26 | 2.98E-03 | ENSRNOT00000028006 |
| Svip-201 | 1 | 24.4 | -2.27 | 1.46E-04 | ENSRNOT00000056024 |
| Synm-202 | 1 | 14.35 | 2.45 | 4.20E-20 | ENSRNOT00000076511 |
| Nr2f2-201 | 1 | 56.91 | 2.3 | 5.43E-09 | ENSRNOT00000014152 |
| Mfge8-202 | 1 | 149.4 | 2.01 | 5.46E-28 | ENSRNOT00000079100 |
| Abhd2-201 | 1 | 34.5 | 2.17 | 1.71E-09 | ENSRNOT00000023506 |
| Zfp710-201 | 1 | 7.68 | 2.25 | 1.68E-04 | ENSRNOT00000064034 |
| Pde8a-204 | 1 | 10.75 | 2.1 | 0.03 | ENSRNOT00000103564 |

| | | | | | |
|---|---|---|---|---|---|
| Il16-201 | 1 | 9.69 | -3.05 | 3.07E-12 | ENSRNOT00000016289 |
| Eed-203 | 1 | 5.41 | -2.57 | 0.05 | ENSRNOT00000115575 |
| Slco2b1-203 | 1 | 18.14 | 4.43 | 1.37E-10 | ENSRNOT00000095615 |
| Plekhb1-202 | 1 | 12.99 | -2.04 | 0.02 | ENSRNOT00000082697 |
| Hbb-204 | 1 | 2,791.01 | -2.93 | 5.56E-31 | ENSRNOT00000089125 |
| Hbb-203 | 1 | 224.32 | -2.12 | 1.35E-12 | ENSRNOT00000089102 |
| Hbb-206 | 1 | 176.27 | -2.14 | 1.43E-12 | ENSRNOT00000090678 |
| Hbb-209 | 1 | 161.79 | -2.11 | 2.59E-11 | ENSRNOT00000113702 |
| Hbb-201 | 1 | 713.7 | -2.85 | 1.72E-31 | ENSRNOT00000019913 |
| Hbb-207 | 1 | 723.49 | -2.99 | 3.86E-34 | ENSRNOT00000090745 |
| Mical2-204 | 1 | 15.1 | 2.18 | 5.40E-08 | ENSRNOT00000081595 |
| Mical2-205 | 1 | 54.41 | 2.17 | 2.25E-21 | ENSRNOT00000102148 |
| Parva-201 | 1 | 25.83 | 2.95 | 9.71E-07 | ENSRNOT00000021671 |
| Spon1-201 | 1 | 25.72 | -2.33 | 7.62E-21 | ENSRNOT00000092099 |
| Vps35l-202 | 1 | 6.4 | 2.55 | 8.35E-03 | ENSRNOT00000083863 |
| Acsm3-201 | 1 | 5.41 | -6.1 | 1.67E-03 | ENSRNOT00000020039 |
| Aqp8-201 | 1 | 33.96 | 3.41 | 5.14E-14 | ENSRNOT00000019939 |
| Spns1-204 | 1 | 8.32 | -2.51 | 6.27E-03 | ENSRNOT00000105544 |
| Spns1-203 | 1 | 7.4 | 2.53 | 0.05 | ENSRNOT00000102967 |
| Aldoa-203 | 1 | 138.68 | 2.08 | 5.61E-23 | ENSRNOT00000112001 |
| Zfp689-206 | 1 | 5.01 | 822.61 | 3.83E-04 | ENSRNOT00000024969 |
| Rnf40-201 | 1 | 8.2 | 2.04 | 8.38E-03 | ENSRNOT00000025499 |
| Chst15-201 | 1 | 13.88 | 2.36 | 2.37E-08 | ENSRNOT00000022157 |
| Ptpre-202 | 1 | 10.38 | 2.12 | 8.69E-03 | ENSRNOT00000098076 |
| Cyp2e1-201 | 1 | 10.15 | -2.67 | 3.18E-03 | ENSRNOT00000016883 |
| Scgb1c1-201 | 1 | 12.46 | -11.82 | 3.50E-04 | ENSRNOT00000017137 |
| Deaf1-203 | 1 | 10.96 | -2.15 | 3.51E-04 | ENSRNOT00000098192 |
| H19-201 | 1 | 207.3 | -2.24 | 2.77E-23 | ENSRNOT00000102917 |
| Igf2-201 | 1 | 37.12 | -2.07 | 1.44E-15 | ENSRNOT00000080246 |
| Trpm5-201 | 1 | 12.28 | -3.05 | 7.20E-11 | ENSRNOT00000040850 |
| Nap1l4-202 | 1 | 12.14 | 2.56 | 0.04 | ENSRNOT00000087961 |
| Dhcr7-201 | 1 | 19.15 | 2.04 | 1.05E-06 | ENSRNOT00000028195 |
| Chka-201 | 1 | 12.98 | -2.17 | 3.42E-04 | ENSRNOT00000022824 |
| Aldh3b1-201 | 1 | 9.41 | 3.5 | 6.77E-04 | ENSRNOT00000023789 |
| Ndufv1-201 | 1 | 9.33 | 7.39 | 0.01 | ENSRNOT00000024517 |
| Clcf1-201 | 1 | 7.67 | -3.5 | 4.22E-03 | ENSRNOT00000025342 |
| Brms1-204 | 1 | 8.08 | -4.68 | 0.02 | ENSRNOT00000113489 |
| Tmem151a-201 | 1 | 12.61 | -3.39 | 4.90E-05 | ENSRNOT00000075365 |
| Efemp2-208 | 1 | 28.48 | 2.11 | 1.18E-06 | ENSRNOT00000117314 |
| Fads2-201 | 1 | 105.36 | 2.19 | 3.55E-12 | ENSRNOT00000027756 |
| Gna14-201 | 1 | 5.84 | -5.58 | 9.97E-06 | ENSRNOT00000020201 |
| Aldh1a7-201 | 1 | 5.67 | -8.59 | 2.07E-06 | ENSRNOT00000024093 |
| Acta2-201 | 1 | 84.28 | 2.33 | 1.90E-19 | ENSRNOT00000083468 |
| Kif11-202 | 1 | 11.91 | -15.14 | 1.29E-07 | ENSRNOT00000095978 |
| Rbp4-201 | 1 | 57.58 | -4.5 | 1.56E-23 | ENSRNOT00000021055 |
| Got1-201 | 1 | 37.75 | 2.15 | 2.24E-11 | ENSRNOT00000022309 |
| Cyp17a1-201 | 1 | 274.92 | 2.4 | 3.66E-25 | ENSRNOT00000027160 |
| Smc3-201 | 1 | 6.46 | 1,646.93 | 1.26E-04 | ENSRNOT00000019560 |
| Vwa2-201 | 1 | 7.8 | -2.69 | 6.74E-07 | ENSRNOT00000051834 |
| Afap1l2-203 | 1 | 6.84 | 2.73 | 0.02 | ENSRNOT00000105824 |
| Emx2-201 | 1 | 22.97 | -2.2 | 2.16E-11 | ENSRNOT00000012564 |

| | | | | | |
|---|---|---|---|---|---|
| Serf1-201 | 2 | 21.35 | -2.08 | 0.04 | ENSRNOT00000076577 |
| Il31ra-201 | 2 | 5.89 | -2.43 | 0.03 | ENSRNOT00000041452 |
| Ddx4-203 | 2 | 26.1 | -2.08 | 2.61E-11 | ENSRNOT00000117202 |
| Mtrex-202 | 2 | 7.19 | 9.38 | 0.01 | ENSRNOT00000095778 |
| Nnt-205 | 2 | 5.18 | -1,152.25 | 9.58E-03 | ENSRNOT00000112693 |
| Ccdc152-201 | 2 | 8.09 | -3.29 | 0.01 | ENSRNOT00000060492 |
| Card6-202 | 2 | 10.4 | 2.04 | 4.62E-04 | ENSRNOT00000108176 |
| Prlr-201 | 2 | 11.17 | 2.38 | 0.01 | ENSRNOT00000080786 |
| Zfr-202 | 2 | 5.29 | 3.46 | 1.46E-03 | ENSRNOT00000080241 |
| Car3-201 | 2 | 15.27 | -3.65 | 1.19E-05 | ENSRNOT00000014180 |
| Fabp4-201 | 2 | 80.17 | -2.57 | 9.81E-07 | ENSRNOT00000014701 |
| Zc2hc1a-201 | 2 | 5.48 | 4.75 | 0.02 | ENSRNOT00000066966 |
| Cpa3-201 | 2 | 13.34 | -4.96 | 3.65E-06 | ENSRNOT00000014970 |
| Fndc3b-201 | 2 | 70.54 | 2.12 | 2.58E-28 | ENSRNOT00000034096 |
| Pld1-201 | 2 | 12.19 | -2.58 | 2.44E-04 | ENSRNOT00000039296 |
| Pld1-202 | 2 | 44.05 | 2.91 | 1.41E-49 | ENSRNOT00000039308 |
| Actl6a-202 | 2 | 17.24 | -2,046.37 | 3.72E-03 | ENSRNOT00000098294 |
| Fxr1-205 | 2 | 21.5 | -2.2 | 5.93E-11 | ENSRNOT00000117973 |
| Bbs7-204 | 2 | 5.36 | -4.92 | 0.03 | ENSRNOT00000112087 |
| Mgarp-201 | 2 | 6.63 | 4.17 | 0.02 | ENSRNOT00000016475 |
| Foxo1-201 | 2 | 77.47 | 3.63 | 1.85E-45 | ENSRNOT00000018244 |
| Frem2-201 | 2 | 11.14 | -2.36 | 2.27E-25 | ENSRNOT00000031487 |
| Postn-202 | 2 | 42.37 | -2.23 | 3.71E-14 | ENSRNOT00000084527 |
| Postn-204 | 2 | 16.42 | -1,007.10 | 6.94E-09 | ENSRNOT00000110085 |
| Postn-201 | 2 | 90.6 | -2.54 | 4.65E-31 | ENSRNOT00000017453 |
| Exosc8-205 | 2 | 12.13 | -2.33 | 6.75E-03 | ENSRNOT00000111232 |
| Sohlh2-201 | 2 | 7.7 | -2.91 | 9.40E-04 | ENSRNOT00000044424 |
| Etfdh-204 | 2 | 6.95 | 3.78 | 5.47E-04 | ENSRNOT00000105634 |
| Dclk2-202 | 2 | 30.57 | -2.23 | 1.59E-21 | ENSRNOT00000057062 |
| Dclk2-204 | 2 | 11.88 | 2.44 | 8.51E-06 | ENSRNOT00000079954 |
| Hdgf-202 | 2 | 16.18 | -314.06 | 5.09E-08 | ENSRNOT00000065211 |
| Sema4a-201 | 2 | 5.88 | 2.91 | 0.01 | ENSRNOT00000056862 |
| Scamp3-204 | 2 | 17.76 | 2.07 | 0.01 | ENSRNOT00000111900 |
| Them5-201 | 2 | 162.77 | -3.23 | 6.45E-45 | ENSRNOT00000066020 |
| Rorc-204 | 2 | 8.04 | -3.58 | 4.40E-06 | ENSRNOT00000118722 |
| Bnipl-201 | 2 | 7.15 | -5.56 | 2.21E-04 | ENSRNOT00000028671 |
| C2H1orf54-201 | 2 | 30.44 | -2.25 | 0.01 | ENSRNOT00000091888 |
| H2bc18-201 | 2 | 7.26 | -2.8 | 2.83E-04 | ENSRNOT00000095604 |
| Hmgcs2-201 | 2 | 586.14 | 3.6 | 7.19E-77 | ENSRNOT00000026121 |
| Hsd3b3-201 | 2 | 752.08 | 2.18 | 2.50E-20 | ENSRNOT00000056172 |
| Hsd3b1-201 | 2 | 2,082.48 | 2.39 | 2.13E-20 | ENSRNOT00000086835 |
| Hao2-202 | 2 | 9.82 | 4.22 | 2.18E-03 | ENSRNOT00000091444 |
| Hao2-201 | 2 | 13.9 | 2.37 | 0.02 | ENSRNOT00000046942 |
| Wnt2b-202 | 2 | 16.75 | -3.82 | 7.02E-16 | ENSRNOT00000115886 |
| Wnt2b-201 | 2 | 20.7 | -4.17 | 2.81E-16 | ENSRNOT00000019349 |
| Gstm1-201 | 2 | 130.89 | 2.07 | 7.12E-15 | ENSRNOT00000047139 |
| Ampd2-202 | 2 | 11.01 | 2.03 | 1.27E-03 | ENSRNOT00000088657 |
| Rnpc3-201 | 2 | 14.62 | -2.24 | 1.25E-05 | ENSRNOT00000023366 |
| Col11a1-201 | 2 | 113.7 | 2.14 | 5.90E-39 | ENSRNOT00000068413 |
| Ptbp2-204 | 2 | 30.8 | -2.31 | 4.36E-07 | ENSRNOT00000108304 |
| Ptbp2-201 | 2 | 25.27 | -2.72 | 2.06E-09 | ENSRNOT00000015036 |

| | | | | | |
|---|---|---|---|---|---|
| Cnn3-202 | 2 | 168.92 | -2.06 | 8.69E-27 | ENSRNOT00000104611 |
| Bcar3-201 | 2 | 47.96 | 2.41 | 2.57E-26 | ENSRNOT00000065111 |
| Sec24b-203 | 2 | 13.57 | 2.59 | 2.88E-08 | ENSRNOT00000109543 |
| Lef1-201 | 2 | 6.18 | -2.71 | 3.85E-03 | ENSRNOT00000013694 |
| Ddit4l-202 | 2 | 24.25 | 2.11 | 2.09E-10 | ENSRNOT00000111236 |
| Bmpr1b-204 | 2 | 17.77 | 3.58 | 1.00E-07 | ENSRNOT00000114929 |
| Odf2l-204 | 2 | 8.28 | -2.48 | 2.66E-03 | ENSRNOT00000119248 |
| Prkacb-202 | 2 | 10.31 | -2.35 | 1.65E-03 | ENSRNOT00000109432 |
| Ifi44-202 | 2 | 39.64 | -2.14 | 5.09E-08 | ENSRNOT00000084873 |
| Ptgfr-201 | 2 | 9.02 | -2.01 | 1.39E-03 | ENSRNOT00000071195 |
| ENSRNOT00000116619 | 3 | 18.03 | -3.05 | 1.37E-07 | ENSRNOT00000116619 |
| ENSRNOT00000099227 | 3 | 53.88 | -2.37 | 5.47E-06 | ENSRNOT00000099227 |
| Nsmf-203 | 3 | 6.02 | -2.46 | 0.01 | ENSRNOT00000048137 |
| Man1b1-202 | 3 | 10.53 | -2.5 | 5.96E-03 | ENSRNOT00000102607 |
| Clic3-201 | 3 | 8.69 | -4.38 | 0.03 | ENSRNOT00000020497 |
| Ptgds-201 | 3 | 14.13 | -4.3 | 2.66E-03 | ENSRNOT00000020926 |
| Sohlh1-201 | 3 | 54.97 | -2.54 | 1.96E-12 | ENSRNOT00000051729 |
| Qsox2-201 | 3 | 35.31 | 2.12 | 3.48E-16 | ENSRNOT00000061735 |
| Qsox2-202 | 3 | 5.25 | -4.85 | 0.01 | ENSRNOT00000116146 |
| Col5a1-201 | 3 | 35.27 | 2.43 | 2.27E-33 | ENSRNOT00000012334 |
| Gtf3c4-202 | 3 | 5.67 | 2.02 | 0.04 | ENSRNOT00000105740 |
| Phyhd1-201 | 3 | 90.89 | -3.56 | 4.29E-56 | ENSRNOT00000022552 |
| Ass1-202 | 3 | 109.66 | 4.71 | 3.22E-32 | ENSRNOT00000093365 |
| Lamc3-201 | 3 | 56.05 | 3.28 | 1.21E-48 | ENSRNOT00000090137 |
| Aif1l-201 | 3 | 105.37 | 3.19 | 4.70E-56 | ENSRNOT00000013286 |
| Fam78a-201 | 3 | 6.88 | 3.06 | 2.16E-06 | ENSRNOT00000035580 |
| Prrc2b-201 | 3 | 6.07 | 2.21 | 3.30E-04 | ENSRNOT00000013546 |
| Eng-201 | 3 | 13.49 | 2.08 | 4.56E-04 | ENSRNOT00000071801 |
| Gsn-201 | 3 | 14.34 | 3.01 | 2.79E-03 | ENSRNOT00000025857 |
| Nr5a1-201 | 3 | 135.56 | 2.71 | 1.32E-35 | ENSRNOT00000017651 |
| Scai-203 | 3 | 11.55 | -2.03 | 1.28E-08 | ENSRNOT00000104956 |
| Zeb2-201 | 3 | 7.28 | 2.42 | 3.57E-04 | ENSRNOT00000006350 |
| Orc4-203 | 3 | 21.66 | -2.13 | 6.68E-05 | ENSRNOT00000103065 |
| Rif1-201 | 3 | 17.65 | -4.18 | 5.09E-11 | ENSRNOT00000090575 |
| Marchf7-202 | 3 | 10.53 | -5.82 | 6.85E-05 | ENSRNOT00000085563 |
| Pla2r1-201 | 3 | 8.83 | -2.16 | 6.35E-08 | ENSRNOT00000011003 |
| Grb14-201 | 3 | 87.05 | 2.19 | 8.44E-20 | ENSRNOT00000089243 |
| Grb14-202 | 3 | 68.9 | 2.27 | 3.34E-08 | ENSRNOT00000097641 |
| Cobll1-202 | 3 | 7.27 | 2.01 | 2.86E-03 | ENSRNOT00000104492 |
| Sp5-201 | 3 | 6.39 | -3.44 | 3.45E-03 | ENSRNOT00000078819 |
| Gorasp2-203 | 3 | 39.01 | 2.17 | 1.27E-08 | ENSRNOT00000111143 |
| Hnrnpa3-204 | 3 | 19.61 | -16.79 | 1.13E-04 | ENSRNOT00000114122 |
| Nfe2l2-203 | 3 | 21.52 | -20.38 | 2.44E-14 | ENSRNOT00000110389 |
| Ctnnd1-201 | 3 | 6.79 | 2.09 | 8.08E-03 | ENSRNOT00000064009 |
| Aplnr-201 | 3 | 25.11 | 2.05 | 9.23E-15 | ENSRNOT00000012379 |
| Celf1-202 | 3 | 22.96 | 2.02 | 9.34E-13 | ENSRNOT00000106910 |
| Chst1-206 | 3 | 5.78 | 3 | 1.27E-03 | ENSRNOT00000114058 |
| Chst1-201 | 3 | 7.97 | 3.56 | 4.06E-03 | ENSRNOT00000010510 |
| Tp53i11-201 | 3 | 74.4 | 2.11 | 1.50E-15 | ENSRNOT00000057001 |
| Tp53i11-202 | 3 | 41.81 | 2.53 | 2.76E-22 | ENSRNOT00000116468 |
| Pdhx-203 | 3 | 14.27 | 2.05 | 0.02 | ENSRNOT00000108445 |

| | | | | | |
|---|---|---|---|---|---|
| Eif3m-205 | 3 | 33.5 | -3.44 | 4.51E-06 | ENSRNOT00000117477 |
| Wt1-203 | 3 | 82.39 | -2.09 | 1.46E-20 | ENSRNOT00000102019 |
| Fibin-201 | 3 | 5.79 | 2.8 | 0.02 | ENSRNOT00000006203 |
| Aven-201 | 3 | 12.57 | -5.37 | 9.29E-05 | ENSRNOT00000065752 |
| Thbs1-201 | 3 | 189.46 | 2.02 | 6.78E-23 | ENSRNOT00000083351 |
| Exd1-201 | 3 | 8.43 | -2.01 | 1.32E-03 | ENSRNOT00000038365 |
| Nusap1-201 | 3 | 9.28 | 313.88 | 1.64E-06 | ENSRNOT00000052312 |
| Nusap1-203 | 3 | 9.22 | -1,404.17 | 6.17E-03 | ENSRNOT00000095179 |
| Frmd5-202 | 3 | 9.19 | 2.12 | 0.01 | ENSRNOT00000064663 |
| Slc30a4-202 | 3 | 11.15 | -2 | 4.74E-05 | ENSRNOT00000106881 |
| Sema6d-203 | 3 | 5.55 | -2.17 | 0.02 | ENSRNOT00000098848 |
| Sema6d-206 | 3 | 10.64 | -2.46 | 6.41E-08 | ENSRNOT00000113650 |
| Sema6d-202 | 3 | 8.76 | -2.75 | 5.75E-07 | ENSRNOT00000090802 |
| Myef2-201 | 3 | 5.53 | -3.65 | 9.85E-04 | ENSRNOT00000007103 |
| Gpat2-201 | 3 | 11.65 | -2.12 | 8.88E-05 | ENSRNOT00000018677 |
| Anapc1-203 | 3 | 11.79 | 2.28 | 5.20E-15 | ENSRNOT00000111998 |
| Adam33-202 | 3 | 6.09 | -3.67 | 1.29E-03 | ENSRNOT00000051404 |
| Adra1d-201 | 3 | 5.75 | 2.79 | 6.54E-03 | ENSRNOT00000028877 |
| Lrrn4-201 | 3 | 19.49 | -2.31 | 3.13E-10 | ENSRNOT00000040802 |
| Defb24-201 | 3 | 81.74 | -2.24 | 8.88E-05 | ENSRNOT00000055408 |
| Hm13-202 | 3 | 47.24 | 2.12 | 2.45E-09 | ENSRNOT00000087075 |
| Bpifb6-201 | 3 | 8.54 | -3.01 | 2.20E-03 | ENSRNOT00000017876 |
| Procr-202 | 3 | 80.68 | -2.04 | 7.17E-09 | ENSRNOT00000114811 |
| Rbm39-205 | 3 | 80.33 | -2.45 | 7.76E-47 | ENSRNOT00000117296 |
| Mroh8-201 | 3 | 16.28 | -2.74 | 7.55E-15 | ENSRNOT00000055219 |
| Lbp-201 | 3 | 5.13 | 2.66 | 0.04 | ENSRNOT00000019787 |
| Mafb-201 | 3 | 12.06 | 2.46 | 2.62E-03 | ENSRNOT00000021452 |
| Zhx3-201 | 3 | 6.7 | 2.54 | 3.40E-04 | ENSRNOT00000032588 |
| Chd6-202 | 3 | 6.71 | 3.54 | 4.30E-04 | ENSRNOT00000089958 |
| Tox2-203 | 3 | 30.71 | 3.01 | 4.86E-25 | ENSRNOT00000119983 |
| Tox2-201 | 3 | 9.47 | 6.45 | 2.24E-04 | ENSRNOT00000064159 |
| Sdc4-201 | 3 | 48.6 | -2.17 | 6.02E-22 | ENSRNOT00000019386 |
| Spint3-201 | 3 | 19.18 | 3.17 | 0.02 | ENSRNOT00000078633 |
| Pltp-201 | 3 | 38.52 | 2.18 | 6.72E-07 | ENSRNOT00000022594 |
| Pcif1-202 | 3 | 13.46 | 2.56 | 8.65E-05 | ENSRNOT00000094436 |
| Sulf2-201 | 3 | 9.43 | 2.04 | 1.17E-03 | ENSRNOT00000008478 |
| Sall4-202 | 3 | 11.15 | 2.01 | 6.30E-04 | ENSRNOT00000071119 |
| Nelfcd-201 | 3 | 5.81 | -2.49 | 0.05 | ENSRNOT00000073634 |
| LOC103690425-201 | 3 | 8.28 | -2.01 | 0.03 | ENSRNOT00000109993 |
| Sycp2-202 | 3 | 14.17 | -5.82 | 1.85E-36 | ENSRNOT00000079019 |
| Tcfl5-201 | 3 | 7.86 | -2.4 | 8.75E-03 | ENSRNOT00000082266 |
| Sox18-201 | 3 | 32.79 | 2.23 | 2.46E-08 | ENSRNOT00000021790 |
| Fastk-204 | 4 | 11.89 | 2.87 | 0.03 | ENSRNOT00000118434 |
| Slc4a2-202 | 4 | 5.76 | 1,502.93 | 5.89E-05 | ENSRNOT00000108170 |
| LOC103692025-202 | 4 | 29.32 | 3.28 | 1.25E-09 | ENSRNOT00000104480 |
| RGD1565355-208 | 4 | 9.58 | -2.01 | 0.01 | ENSRNOT00000091249 |
| Sema3a-202 | 4 | 19.51 | -2 | 1.72E-13 | ENSRNOT00000096965 |
| Elapor2-202 | 4 | 18.39 | 2.31 | 1.64E-23 | ENSRNOT00000100645 |
| Elapor2-201 | 4 | 10.02 | 5.48 | 8.44E-07 | ENSRNOT00000007723 |
| Abcb1b-202 | 4 | 20.51 | 2.74 | 3.90E-21 | ENSRNOT00000047126 |
| Cdk14-202 | 4 | 13.51 | -13.39 | 1.59E-16 | ENSRNOT00000094279 |

| | | | | | |
|---|---|---|---|---|---|
| Peg10-202 | 4 | 8.82 | 2.97 | 1.02E-04 | ENSRNOT00000104707 |
| Phf14-208 | 4 | 5.77 | -2.55 | 6.20E-08 | ENSRNOT00000108109 |
| Asz1-201 | 4 | 6.66 | -2.55 | 0.05 | ENSRNOT00000091311 |
| Impdh1-203 | 4 | 49.65 | 2.12 | 1.10E-16 | ENSRNOT00000098302 |
| Impdh1-204 | 4 | 15.77 | 2.09 | 9.17E-08 | ENSRNOT00000110384 |
| Impdh1-201 | 4 | 21.47 | 2.34 | 3.12E-05 | ENSRNOT00000027173 |
| Cpa2-201 | 4 | 72.54 | -2.87 | 2.57E-32 | ENSRNOT00000029608 |
| Podxl-202 | 4 | 71.02 | -2.33 | 1.52E-32 | ENSRNOT00000077406 |
| Plxna4-201 | 4 | 11.95 | -3.19 | 5.31E-40 | ENSRNOT00000017536 |
| Chchd3-204 | 4 | 24.36 | -3.26 | 4.20E-03 | ENSRNOT00000107634 |
| RGD1565367-201 | 4 | 11.47 | -3.91 | 2.17E-06 | ENSRNOT00000038950 |
| Stk31-201 | 4 | 8.21 | -2.71 | 2.35E-04 | ENSRNOT00000086307 |
| Stk31-203 | 4 | 15 | -2.88 | 1.87E-12 | ENSRNOT00000118450 |
| Osbpl3-203 | 4 | 5.92 | -2 | 0.03 | ENSRNOT00000117089 |
| Mrpl35-201 | 4 | 6.32 | -6.02 | 0.03 | ENSRNOT00000012021 |
| Immt-203 | 4 | 11.76 | 3.85 | 5.21E-05 | ENSRNOT00000098740 |
| Tcf7l1-201 | 4 | 18.54 | 2.01 | 3.85E-06 | ENSRNOT00000020005 |
| Actg2-201 | 4 | 14.21 | 2.39 | 0.04 | ENSRNOT00000042699 |
| Figla-201 | 4 | 166.52 | -3.34 | 4.97E-37 | ENSRNOT00000021272 |
| Tia1-201 | 4 | 10.74 | -3.02 | 0.04 | ENSRNOT00000023103 |
| AABR07061385.2-201 | 4 | 61.01 | 2.03 | 6.60E-10 | ENSRNOT00000078142 |
| Klf15-201 | 4 | 8.23 | 2.33 | 0.01 | ENSRNOT00000024011 |
| Fbln2-203 | 4 | 26.67 | -3.16 | 6.22E-32 | ENSRNOT00000115833 |
| Cntn3-201 | 4 | 6.24 | -2.43 | 6.02E-04 | ENSRNOT00000008221 |
| Trnt1-203 | 4 | 11.04 | 8.07 | 0.04 | ENSRNOT00000119654 |
| Lmcd1-202 | 4 | 9.76 | -5.59 | 4.26E-03 | ENSRNOT00000117173 |
| Pparg-202 | 4 | 11.18 | 3.83 | 3.34E-05 | ENSRNOT00000082969 |
| H1f8-201 | 4 | 41.16 | 2.03 | 2.10E-06 | ENSRNOT00000042673 |
| Plxnd1-201 | 4 | 34.79 | 2.04 | 3.71E-22 | ENSRNOT00000032158 |
| Erc1-202 | 4 | 10 | 2.08 | 1.84E-04 | ENSRNOT00000091473 |
| Tspan11-201 | 4 | 37.87 | 2.1 | 6.59E-11 | ENSRNOT00000087834 |
| Lpal2-201 | 4 | 15.05 | -4.03 | 1.11E-03 | ENSRNOT00000034403 |
| Fam234b-201 | 4 | 7.11 | 2.44 | 2.59E-04 | ENSRNOT00000068663 |
| Atf7ip-201 | 4 | 11.7 | 2.07 | 5.32E-07 | ENSRNOT00000011803 |
| Plbd1-201 | 4 | 95.01 | 3.62 | 6.64E-34 | ENSRNOT00000011930 |
| Art4-201 | 4 | 12.73 | 2.43 | 8.33E-04 | ENSRNOT00000007485 |
| Rerg-201 | 4 | 39.24 | 3.12 | 2.88E-14 | ENSRNOT00000030850 |
| Med21-201 | 4 | 157.89 | -2.05 | 3.36E-15 | ENSRNOT00000002500 |
| Caprin2-201 | 4 | 6.45 | -2.78 | 1.08E-03 | ENSRNOT00000070907 |
| Resf1-202 | 4 | 43.2 | -2.63 | 2.93E-32 | ENSRNOT00000094270 |
| Lactb2-201 | 5 | 5.75 | -8.93 | 7.03E-03 | ENSRNOT00000010369 |
| Plag1-201 | 5 | 11.91 | -2.37 | 2.75E-07 | ENSRNOT00000011721 |
| Sdr16c5-201 | 5 | 6.42 | -5.8 | 1.08E-03 | ENSRNOT00000011813 |
| ENSRNOT00000113061 | 5 | 276.92 | 2.26 | 1.03E-07 | ENSRNOT00000113061 |
| Plekhf2-202 | 5 | 5.97 | 6.98 | 4.09E-04 | ENSRNOT00000094306 |
| Slc26a7-203 | 5 | 8.18 | 2.12 | 0.05 | ENSRNOT00000093409 |
| Gpr63-205 | 5 | 6.69 | -2.42 | 2.81E-03 | ENSRNOT00000117217 |
| Zcchc7-202 | 5 | 20.29 | -2.81 | 4.95E-07 | ENSRNOT00000112430 |
| Fsd1l-202 | 5 | 6.55 | -2.08 | 8.88E-07 | ENSRNOT00000112399 |
| Akap2-205 | 5 | 9.14 | 2.22 | 6.26E-05 | ENSRNOT00000098662 |
| Inip-201 | 5 | 5.19 | -5.64 | 0.03 | ENSRNOT00000095336 |

| | | | | | |
|---|---|---|---|---|---|
| Bspry-201 | 5 | 9.28 | -2.58 | 2.35E-03 | ENSRNOT00000020366 |
| Alad-204 | 5 | 7.79 | -416.11 | 9.13E-03 | ENSRNOT00000077013 |
| Alad-202 | 5 | 6.38 | 515.47 | 1.69E-03 | ENSRNOT00000075916 |
| Ptprd-205 | 5 | 5.66 | -4.07 | 8.33E-14 | ENSRNOT00000115119 |
| Elavl2-203 | 5 | 37.57 | -2.42 | 7.17E-25 | ENSRNOT00000093032 |
| Hook1-201 | 5 | 6.07 | -2.61 | 4.72E-04 | ENSRNOT00000012396 |
| Nfia-201 | 5 | 11.25 | 2.47 | 8.38E-03 | ENSRNOT00000004017 |
| Patj-201 | 5 | 5.68 | -2.58 | 4.56E-03 | ENSRNOT00000010211 |
| Czib-205 | 5 | 13.47 | -525.16 | 5.53E-03 | ENSRNOT00000104267 |
| Osbpl9-204 | 5 | 5.5 | -45.91 | 9.68E-04 | ENSRNOT00000105115 |
| Faf1-203 | 5 | 14.82 | -50.84 | 5.55E-17 | ENSRNOT00000108187 |
| Faah-201 | 5 | 9.5 | -2.09 | 1.33E-04 | ENSRNOT00000015667 |
| Tie1-201 | 5 | 42.12 | 2.3 | 5.68E-26 | ENSRNOT00000027414 |
| Ppih-202 | 5 | 18.64 | -142.19 | 6.01E-04 | ENSRNOT00000067943 |
| Rspo1-201 | 5 | 59.96 | -3.31 | 2.43E-43 | ENSRNOT00000012811 |
| Map7d1-202 | 5 | 17.81 | 2.09 | 4.77E-06 | ENSRNOT00000108369 |
| Sfpq-201 | 5 | 129.18 | 2.07 | 2.15E-34 | ENSRNOT00000084202 |
| Sync-203 | 5 | 13.96 | -3.43 | 4.35E-16 | ENSRNOT00000119345 |
| Sync-201 | 5 | 34.08 | -2.18 | 8.31E-21 | ENSRNOT00000010730 |
| Ythdf2-201 | 5 | 5.6 | 4.09 | 0.01 | ENSRNOT00000014479 |
| Cd164l2-201 | 5 | 19.75 | -2.58 | 6.08E-03 | ENSRNOT00000012626 |
| Cep85-201 | 5 | 21.66 | -2.09 | 4.27E-13 | ENSRNOT00000065983 |
| Paqr7-201 | 5 | 8.89 | 2.49 | 5.72E-03 | ENSRNOT00000022713 |
| Srrm1-204 | 5 | 25.25 | -2.18 | 3.64E-05 | ENSRNOT00000116482 |
| Rap1gap-201 | 5 | 6.26 | -2.11 | 0.02 | ENSRNOT00000046802 |
| Eif4g3-209 | 5 | 7.65 | 17.52 | 1.23E-15 | ENSRNOT00000115071 |
| Pla2g5-202 | 5 | 5.64 | -103.45 | 4.47E-05 | ENSRNOT00000098996 |
| Pla2g5-201 | 5 | 16.39 | -2.29 | 1.92E-04 | ENSRNOT00000022716 |
| Pla2g2a-201 | 5 | 27.38 | -3.04 | 6.65E-05 | ENSRNOT00000022827 |
| Oog1-201 | 5 | 12.99 | 2.09 | 0.03 | ENSRNOT00000046115 |
| Mthfr-201 | 5 | 22.85 | -3.1 | 6.84E-18 | ENSRNOT00000011384 |
| Fbxo44-202 | 5 | 29.77 | 2.32 | 5.88E-09 | ENSRNOT00000101645 |
| Fbxo2-201 | 5 | 7.52 | 3.08 | 0.04 | ENSRNOT00000012670 |
| Masp2-201 | 5 | 9.32 | -2.22 | 8.86E-03 | ENSRNOT00000016317 |
| Pik3cd-202 | 5 | 83.01 | 2.25 | 2.36E-42 | ENSRNOT00000095643 |
| Spsb1-201 | 5 | 96.46 | 2.33 | 3.00E-29 | ENSRNOT00000085251 |
| Slc2a5-201 | 5 | 9.86 | -303.24 | 3.85E-05 | ENSRNOT00000024054 |
| Plekhg5-204 | 5 | 5.34 | 2.45 | 0.04 | ENSRNOT00000107438 |
| C5h1orf174-202 | 5 | 11.45 | 9.23 | 9.13E-04 | ENSRNOT00000112064 |
| Megf6-201 | 5 | 16.23 | -2.7 | 1.20E-18 | ENSRNOT00000000169 |
| Megf6-202 | 5 | 7.57 | -3.22 | 4.90E-05 | ENSRNOT00000055386 |
| Prkcz-201 | 5 | 9.08 | -2.09 | 6.31E-03 | ENSRNOT00000021285 |
| Lhcgr-201 | 6 | 64.32 | 5.78 | 3.32E-71 | ENSRNOT00000022481 |
| Epcam-201 | 6 | 20.21 | -2.79 | 5.56E-06 | ENSRNOT00000021135 |
| Calm2-202 | 6 | 1,227.90 | -2.06 | 2.04E-35 | ENSRNOT00000112676 |
| Epas1-202 | 6 | 37.88 | 2.39 | 5.02E-28 | ENSRNOT00000090420 |
| Epas1-201 | 6 | 9.44 | 2.74 | 4.29E-04 | ENSRNOT00000034991 |
| Prkce-201 | 6 | 10.84 | 2.04 | 3.07E-03 | ENSRNOT00000020959 |
| Pkdcc-202 | 6 | 50.71 | 2.09 | 1.56E-15 | ENSRNOT00000104247 |
| Cyp1b1-202 | 6 | 48.93 | -2.6 | 1.40E-38 | ENSRNOT00000084171 |
| Ltbp1-201 | 6 | 79.31 | 2.47 | 2.54E-48 | ENSRNOT00000047674 |

| | | | | | |
|---|---|---|---|---|---|
| Fosl2-201 | 6 | 14.08 | 6.09 | 2.64E-08 | ENSRNOT00000107475 |
| Fosl2-202 | 6 | 30.23 | 2.54 | 1.64E-13 | ENSRNOT00000113775 |
| Greb1-201 | 6 | 59.68 | 4.4 | 1.89E-110 | ENSRNOT00000032417 |
| Itgb1bp1-203 | 6 | 10 | -5.17 | 0.05 | ENSRNOT00000116410 |
| Iah1-202 | 6 | 6.44 | -25.97 | 0.01 | ENSRNOT00000120299 |
| Iah1-201 | 6 | 153.3 | -2.06 | 5.69E-32 | ENSRNOT00000087180 |
| Rps7-202 | 6 | 7.81 | -2.27 | 0.04 | ENSRNOT00000011333 |
| Tpo-201 | 6 | 9.01 | -2.45 | 1.51E-05 | ENSRNOT00000006526 |
| Tmem18-202 | 6 | 6.07 | 13.05 | 1.29E-03 | ENSRNOT00000079520 |
| Lamb1-201 | 6 | 25.44 | 2.3 | 8.12E-07 | ENSRNOT00000008321 |
| Prkar2b-201 | 6 | 609.21 | 2.12 | 1.01E-42 | ENSRNOT00000012415 |
| Sypl1-203 | 6 | 15.38 | -2,017.26 | 5.16E-03 | ENSRNOT00000097655 |
| Agr3-201 | 6 | 6.14 | -4.12 | 0.03 | ENSRNOT00000006652 |
| Sostdc1-201 | 6 | 14.52 | 5.36 | 1.98E-07 | ENSRNOT00000008106 |
| Arl4a-201 | 6 | 6.29 | -2.1 | 7.87E-05 | ENSRNOT00000108440 |
| Slc25a21-203 | 6 | 7.25 | 2.07 | 0.04 | ENSRNOT00000119506 |
| Esr2-204 | 6 | 10.74 | 2.58 | 1.58E-03 | ENSRNOT00000043602 |
| Mthfd1-204 | 6 | 17.7 | -8.06 | 8.42E-19 | ENSRNOT00000100905 |
| Plekhg3-201 | 6 | 10.87 | 2.08 | 4.54E-04 | ENSRNOT00000008573 |
| Rdh11-203 | 6 | 5.68 | -77.43 | 1.29E-03 | ENSRNOT00000105155 |
| Smoc1-202 | 6 | 196.73 | 2.17 | 2.31E-36 | ENSRNOT00000101574 |
| Smoc1-203 | 6 | 157.39 | 2.17 | 1.60E-49 | ENSRNOT00000114143 |
| Rbm25-201 | 6 | 8.87 | -1,686.68 | 6.99E-03 | ENSRNOT00000003834 |
| Acot2-201 | 6 | 9.03 | 2.74 | 3.93E-03 | ENSRNOT00000013515 |
| Fos-201 | 6 | 18.5 | 3.09 | 6.10E-03 | ENSRNOT00000010712 |
| Flvcr2-201 | 6 | 46.52 | 2.78 | 1.10E-29 | ENSRNOT00000011599 |
| Adck1-202 | 6 | 8.7 | 2.28 | 3.05E-03 | ENSRNOT00000104832 |
| Spata7-207 | 6 | 6.47 | -2.52 | 0.03 | ENSRNOT00000108318 |
| Ttc8-206 | 6 | 6.89 | 21.78 | 5.47E-06 | ENSRNOT00000102593 |
| Ttc7b-201 | 6 | 8.61 | 2.44 | 0.02 | ENSRNOT00000037902 |
| Ifi27-202 | 6 | 16.07 | -2.98 | 0.01 | ENSRNOT00000063970 |
| Serpina9-201 | 6 | 6.37 | -2.74 | 0.04 | ENSRNOT00000012861 |
| Papola-202 | 6 | 10.66 | 4.5 | 5.36E-03 | ENSRNOT00000083732 |
| Eif5-202 | 6 | 6.92 | -3.46 | 0.02 | ENSRNOT00000102717 |
| Klc1-201 | 6 | 7.67 | 6.16 | 5.10E-06 | ENSRNOT00000015935 |
| ENSRNOT00000109415 | 6 | 22.5 | 6.93 | 6.50E-03 | ENSRNOT00000109415 |
| Kif26a-201 | 6 | 5.33 | 2.86 | 5.38E-06 | ENSRNOT00000018278 |
| Pa2g4-201 | 7 | 42.86 | 2.11 | 2.64E-04 | ENSRNOT00000006578 |
| Rab5b-203 | 7 | 12.34 | -3.34 | 0.02 | ENSRNOT00000120020 |
| Zfp709l1-203 | 7 | 13.32 | -2 | 4.01E-13 | ENSRNOT00000086920 |
| Tle2-201 | 7 | 9.93 | -2.05 | 0.01 | ENSRNOT00000061279 |
| Tle5-202 | 7 | 9.76 | -2.34 | 7.73E-03 | ENSRNOT00000098364 |
| Tmem259-201 | 7 | 46.73 | 2.38 | 1.17E-07 | ENSRNOT00000017245 |
| Rab11b-203 | 7 | 9.37 | -2.12 | 1.55E-14 | ENSRNOT00000114766 |
| Sycp3-201 | 7 | 14.56 | -4.6 | 7.81E-05 | ENSRNOT00000007088 |
| Mybpc1-205 | 7 | 6.4 | -3.01 | 2.08E-04 | ENSRNOT00000099906 |
| ENSRNOT00000108822 | 7 | 8.02 | -8.71 | 1.73E-07 | ENSRNOT00000108822 |
| Lum-202 | 7 | 18.63 | 2.02 | 3.56E-14 | ENSRNOT00000111145 |
| Lum-201 | 7 | 429.21 | 2.02 | 8.59E-42 | ENSRNOT00000006109 |
| Tmtc2-201 | 7 | 25.41 | 2.52 | 7.03E-13 | ENSRNOT00000006086 |
| Csrp2-203 | 7 | 15.77 | 2.04 | 5.29E-11 | ENSRNOT00000097951 |

| | | | | | |
|---|---|---|---|---|---|
| Glipr1-201 | 7 | 40.53 | -2.67 | 5.30E-08 | ENSRNOT00000005399 |
| Tspan8-201 | 7 | 94.78 | -4.29 | 1.92E-45 | ENSRNOT00000005909 |
| Ptprb-204 | 7 | 7.2 | 2.57 | 1.08E-14 | ENSRNOT00000119174 |
| Cct2-203 | 7 | 33.66 | -2.9 | 1.12E-03 | ENSRNOT00000113939 |
| Mdm2-202 | 7 | 18.5 | -2.82 | 1.39E-08 | ENSRNOT00000086116 |
| C7h12orf56-201 | 7 | 8.77 | -2.4 | 9.97E-03 | ENSRNOT00000067395 |
| Kcns2-201 | 7 | 8.46 | 2.05 | 7.28E-06 | ENSRNOT00000115427 |
| Osr2-201 | 7 | 73.25 | 2.81 | 2.00E-20 | ENSRNOT00000014879 |
| Cthrc1-201 | 7 | 12.95 | -2.36 | 0.03 | ENSRNOT00000006142 |
| Angpt1-201 | 7 | 11.83 | 2.47 | 7.68E-08 | ENSRNOT00000007979 |
| Eif3e-201 | 7 | 12.55 | -1,360.94 | 6.65E-04 | ENSRNOT00000038340 |
| Deptor-201 | 7 | 12.84 | 3.07 | 0.02 | ENSRNOT00000005722 |
| Myc-201 | 7 | 38.82 | 2.17 | 1.83E-11 | ENSRNOT00000006188 |
| Dennd3-201 | 7 | 5.2 | 2.53 | 0.01 | ENSRNOT00000014541 |
| Ly6c-202 | 7 | 16.87 | -2.78 | 0.01 | ENSRNOT00000081444 |
| RGD1565410-202 | 7 | 9.37 | -4.79 | 8.50E-03 | ENSRNOT00000082405 |
| Ly6l-201 | 7 | 28.64 | -3.49 | 3.58E-03 | ENSRNOT00000071091 |
| Pycr3-201 | 7 | 6.67 | -2.04 | 0.03 | ENSRNOT00000029456 |
| Scrib-202 | 7 | 16.91 | 2.6 | 1.12E-09 | ENSRNOT00000097559 |
| Scx-201 | 7 | 10.6 | -2.68 | 0.03 | ENSRNOT00000029768 |
| Hsf1-204 | 7 | 13.99 | 2.01 | 0.01 | ENSRNOT00000106601 |
| Ppp1r16a-203 | 7 | 5.76 | -3.25 | 1.86E-03 | ENSRNOT00000117934 |
| Ift27-202 | 7 | 48.45 | -5.02 | 1.09E-04 | ENSRNOT00000112331 |
| Fam227a-201 | 7 | 8.35 | -2.86 | 3.27E-05 | ENSRNOT00000056179 |
| Cbx6-201 | 7 | 20.8 | 2.04 | 8.53E-04 | ENSRNOT00000068033 |
| A4galt-202 | 7 | 8.4 | 2.63 | 3.41E-03 | ENSRNOT00000096640 |
| Parvb-202 | 7 | 17.33 | -2.6 | 1.11E-13 | ENSRNOT00000082805 |
| Smc1b-201 | 7 | 12.98 | -2.95 | 1.48E-10 | ENSRNOT00000044883 |
| Zbed4-203 | 7 | 5.57 | 2.18 | 0.02 | ENSRNOT00000103004 |
| Ano6-204 | 7 | 21.06 | 2.05 | 1.07E-16 | ENSRNOT00000098487 |
| Ano6-207 | 7 | 6.13 | -2.1 | 1.80E-04 | ENSRNOT00000110510 |
| Prph-201 | 7 | 16.67 | -2.54 | 1.70E-05 | ENSRNOT00000089211 |
| Nr4a1-201 | 7 | 72.91 | 2.35 | 5.37E-04 | ENSRNOT00000010171 |
| Krt7-201 | 7 | 20.45 | -2.36 | 1.07E-05 | ENSRNOT00000010660 |
| Krt83-201 | 7 | 39.21 | -2.58 | 1.27E-12 | ENSRNOT00000068505 |
| Igfbp6-201 | 7 | 190.89 | 2.18 | 3.77E-28 | ENSRNOT00000014807 |
| Znf740-201 | 7 | 6.78 | 1,662.01 | 2.94E-03 | ENSRNOT00000016240 |
| Pfdn5-202 | 7 | 236.06 | -2.2 | 1.91E-19 | ENSRNOT00000109593 |
| Pfdn5-201 | 7 | 361.71 | -2.07 | 1.36E-28 | ENSRNOT00000017794 |
| Hoxc6-202 | 7 | 7.17 | -3.7 | 6.20E-27 | ENSRNOT00000107142 |
| Hoxc5-201 | 7 | 5.56 | -2.71 | 7.63E-03 | ENSRNOT00000022247 |
| Hoxc4-201 | 7 | 8.55 | -2.16 | 0.02 | ENSRNOT00000022398 |
| Gtsf1-201 | 7 | 102.64 | -2.67 | 2.08E-13 | ENSRNOT00000055275 |
| Ppp1r1a-201 | 7 | 9.95 | -2.59 | 0.01 | ENSRNOT00000055271 |
| ENSRNOT00000105553 | 8 | 302.4 | -2.53 | 7.00E-47 | ENSRNOT00000105553 |
| Pgr-201 | 8 | 5.76 | 2.6 | 4.52E-03 | ENSRNOT00000038313 |
| Fam76b-201 | 8 | 10.96 | -2.51 | 1.13E-05 | ENSRNOT00000068445 |
| Endod1-201 | 8 | 64.34 | 2.18 | 3.58E-25 | ENSRNOT00000033969 |
| Zfp426-201 | 8 | 6.51 | -2.9 | 3.76E-03 | ENSRNOT00000038069 |
| Eif3g-202 | 8 | 14.45 | -2.14 | 0.05 | ENSRNOT00000095125 |
| Zglp1-201 | 8 | 26.97 | -2.81 | 1.64E-04 | ENSRNOT00000068598 |

| | | | | | |
|---|---|---|---|---|---|
| Fli1-201 | 8 | 14.54 | 2.04 | 0.01 | ENSRNOT00000068037 |
| Ets1-204 | 8 | 12.83 | -5.98 | 2.55E-08 | ENSRNOT00000102138 |
| Dcps-208 | 8 | 112.42 | 3.46 | 3.10E-37 | ENSRNOT00000116357 |
| Dcps-205 | 8 | 26.15 | 2.65 | 4.28E-14 | ENSRNOT00000099496 |
| Stt3a-201 | 8 | 20.35 | 5.11 | 1.27E-03 | ENSRNOT00000043518 |
| RGD1311744-202 | 8 | 7.64 | -4.66 | 0.03 | ENSRNOT00000110068 |
| Gramd1b-207 | 8 | 15.3 | 2.19 | 1.21E-05 | ENSRNOT00000118974 |
| Clmp-202 | 8 | 7.25 | 6.78 | 3.17E-03 | ENSRNOT00000108273 |
| LOC120094123-202 | 8 | 35.97 | -4.25 | 1.82E-92 | ENSRNOT00000111823 |
| Tbcel-201 | 8 | 6.8 | 7.09 | 3.31E-04 | ENSRNOT00000059951 |
| Thy1-201 | 8 | 5.18 | 2.98 | 0.05 | ENSRNOT00000008685 |
| Scn4b-201 | 8 | 12.37 | -2 | 4.30E-08 | ENSRNOT00000030152 |
| Rbm7-203 | 8 | 13.67 | -2.35 | 2.77E-05 | ENSRNOT00000111739 |
| Arhgap20-201 | 8 | 5.94 | -2.18 | 0.04 | ENSRNOT00000035989 |
| Gldn-201 | 8 | 61.92 | 2.02 | 2.10E-13 | ENSRNOT00000033900 |
| ENSRNOT00000111617 | 8 | 15.09 | -2.43 | 0.02 | ENSRNOT00000111617 |
| Tspan3-202 | 8 | 23.85 | -2.5 | 1.59E-05 | ENSRNOT00000112890 |
| Hmg20a-202 | 8 | 7.24 | -3.09 | 2.24E-03 | ENSRNOT00000102284 |
| Commd4-205 | 8 | 8.31 | -2.08 | 0.05 | ENSRNOT00000106229 |
| Clk3-203 | 8 | 10.92 | -2.15 | 6.50E-09 | ENSRNOT00000110279 |
| Sema7a-201 | 8 | 41.32 | 2.08 | 1.59E-19 | ENSRNOT00000010620 |
| Cyp11a1-201 | 8 | 40.24 | 3.61 | 1.25E-18 | ENSRNOT00000010831 |
| Cyp11a1-202 | 8 | 59.81 | 5.21 | 2.68E-30 | ENSRNOT00000106594 |
| Itga11-201 | 8 | 5.83 | 2.68 | 8.66E-05 | ENSRNOT00000008971 |
| Zwilch-201 | 8 | 5.08 | -2.57 | 0.03 | ENSRNOT00000012404 |
| Dapk2-201 | 8 | 12.07 | -2.31 | 4.88E-03 | ENSRNOT00000089147 |
| Car12-202 | 8 | 16.77 | -2.16 | 3.23E-09 | ENSRNOT00000105264 |
| Car12-203 | 8 | 6.19 | -5.46 | 4.35E-04 | ENSRNOT00000116915 |
| Fam81a-201 | 8 | 7.72 | -2.4 | 3.16E-03 | ENSRNOT00000090747 |
| Aldh1a2-201 | 8 | 25.1 | -2.04 | 2.85E-08 | ENSRNOT00000079115 |
| Gsta2-201 | 8 | 42.75 | 2.76 | 9.10E-08 | ENSRNOT00000032185 |
| RGD1559995-201 | 8 | 48.35 | -2.57 | 8.27E-07 | ENSRNOT00000000215 |
| Syncrip-204 | 8 | 7.8 | -2.01 | 1.43E-05 | ENSRNOT00000095785 |
| Morf4l1-201 | 8 | 30.77 | -3,035.09 | 2.79E-03 | ENSRNOT00000080421 |
| Plod2-201 | 8 | 177.79 | 2.19 | 2.36E-32 | ENSRNOT00000041859 |
| Rbp1-201 | 8 | 312.64 | -2.12 | 4.91E-35 | ENSRNOT00000018622 |
| Cep70-202 | 8 | 28.98 | -2.7 | 8.49E-11 | ENSRNOT00000081029 |
| Slco2a1-202 | 8 | 13.43 | 2.53 | 6.30E-09 | ENSRNOT00000091275 |
| Tf-202 | 8 | 56.96 | -4.52 | 1.09E-71 | ENSRNOT00000045628 |
| Tmem108-203 | 8 | 5.1 | -2.71 | 2.88E-03 | ENSRNOT00000104880 |
| Ackr4-202 | 8 | 11.9 | -2.21 | 5.80E-05 | ENSRNOT00000119921 |
| Alas1-201 | 8 | 11.09 | 2.82 | 1.53E-06 | ENSRNOT00000083303 |
| Alas1-202 | 8 | 208.37 | 2.83 | 4.53E-18 | ENSRNOT00000086232 |
| Sema3b-201 | 8 | 21.22 | 2.93 | 3.12E-14 | ENSRNOT00000066296 |
| AABR07071482.2-201 | 8 | 8.65 | -37.46 | 3.32E-03 | ENSRNOT00000047247 |
| Rtp3-201 | 8 | 9.19 | 3.25 | 1.24E-03 | ENSRNOT00000056100 |
| Clasp2-205 | 8 | 6.68 | 4.7 | 2.49E-04 | ENSRNOT00000107377 |
| Itga9-201 | 8 | 12.48 | 2.14 | 1.46E-04 | ENSRNOT00000015113 |
| Vill-201 | 8 | 7.33 | -2.49 | 3.73E-03 | ENSRNOT00000015888 |
| LOC100360856-203 | 9 | 23.66 | -2.34 | 9.74E-05 | ENSRNOT00000075578 |
| LOC680955-201 | 9 | 9.92 | -4.53 | 0.03 | ENSRNOT00000061674 |

| | | | | | |
|---|---|---|---|---|---|
| RGD1562107-201 | 9 | 5.38 | -14.67 | 2.11E-03 | ENSRNOT00000038486 |
| Uggt1-202 | 9 | 12.69 | 2.46 | 5.25E-06 | ENSRNOT00000107876 |
| Neurl3-201 | 9 | 6 | -3.24 | 0.04 | ENSRNOT00000038610 |
| Lonrf2-201 | 9 | 21.67 | 2.33 | 9.17E-13 | ENSRNOT00000033964 |
| Fhl2-201 | 9 | 22.22 | 2.5 | 6.91E-03 | ENSRNOT00000023014 |
| Hibch-204 | 9 | 6.28 | -2.75 | 0.02 | ENSRNOT00000116077 |
| Gls-201 | 9 | 146.8 | -2.05 | 1.36E-28 | ENSRNOT00000078073 |
| Gls-203 | 9 | 31.27 | -2.5 | 2.57E-21 | ENSRNOT00000110936 |
| Spats2l-202 | 9 | 10.63 | -2.03 | 0.03 | ENSRNOT00000081497 |
| Atic-203 | 9 | 34.84 | 2.74 | 2.17E-04 | ENSRNOT00000116844 |
| Atic-202 | 9 | 18.95 | -5.14 | 2.08E-03 | ENSRNOT00000098789 |
| Fn1-202 | 9 | 13.54 | 2.11 | 1.35E-11 | ENSRNOT00000057585 |
| Wnt10a-201 | 9 | 7.1 | -2.45 | 7.98E-03 | ENSRNOT00000086375 |
| Ihh-201 | 9 | 93.68 | 4.91 | 6.67E-98 | ENSRNOT00000024419 |
| Dnpep-203 | 9 | 7.97 | -3.14 | 0.02 | ENSRNOT00000108004 |
| Obsl1-202 | 9 | 44.44 | 2.07 | 1.38E-20 | ENSRNOT00000103263 |
| Inha-201 | 9 | 2,678.04 | 2.44 | 1.44E-56 | ENSRNOT00000027227 |
| Epha4-201 | 9 | 6.23 | -2.02 | 3.63E-05 | ENSRNOT00000041689 |
| Mff-207 | 9 | 19.41 | -2.13 | 1.80E-05 | ENSRNOT00000100485 |
| Nppc-201 | 9 | 28.24 | 2.4 | 8.11E-04 | ENSRNOT00000025582 |
| Col6a3-203 | 9 | 97.82 | 2.86 | 4.48E-63 | ENSRNOT00000087595 |
| Tmem232-202 | 9 | 8.47 | -2.67 | 1.97E-03 | ENSRNOT00000093974 |
| Ndufv2-202 | 9 | 26.04 | -4.51 | 2.06E-05 | ENSRNOT00000106172 |
| L3mbtl4-201 | 9 | 5.77 | 2.21 | 0.02 | ENSRNOT00000029210 |
| Myl12b-204 | 9 | 5.46 | -210.84 | 0.04 | ENSRNOT00000108694 |
| Abcc1-202 | 10 | 5.46 | -562.4 | 4.86E-06 | ENSRNOT00000103910 |
| Myh11-201 | 10 | 7.1 | 2.28 | 4.01E-07 | ENSRNOT00000084608 |
| Litaf-202 | 10 | 44.94 | 2.2 | 2.36E-13 | ENSRNOT00000108567 |
| Rpl39l-202 | 10 | 22.3 | -3.03 | 1.14E-08 | ENSRNOT00000105843 |
| Rpl39l-203 | 10 | 15.33 | -253.5 | 2.13E-04 | ENSRNOT00000118208 |
| Ppl-201 | 10 | 13.13 | -3.19 | 2.46E-27 | ENSRNOT00000004026 |
| Glyr1-202 | 10 | 20.62 | 2.2 | 7.50E-08 | ENSRNOT00000095486 |
| Mgrn1-201 | 10 | 13.37 | 2.3 | 1.10E-06 | ENSRNOT00000004416 |
| Mgrn1-206 | 10 | 6.57 | 6.98 | 4.19E-03 | ENSRNOT00000117920 |
| Tnfrsf12a-202 | 10 | 5.12 | -5.94 | 0.05 | ENSRNOT00000107526 |
| Snrpc-ps3-201 | 10 | 27.15 | -6.5 | 1.48E-03 | ENSRNOT00000023626 |
| Gng13-201 | 10 | 18.49 | -3.75 | 5.34E-03 | ENSRNOT00000030273 |
| Msln-201 | 10 | 11.2 | -2.46 | 2.83E-03 | ENSRNOT00000026395 |
| Rhbdl1-201 | 10 | 6.75 | -3.49 | 7.72E-03 | ENSRNOT00000026971 |
| Hba-a3-201 | 10 | 715.14 | 3.03 | 3.85E-61 | ENSRNOT00000048977 |
| Havcr1-201 | 10 | 36.85 | 2.36 | 8.15E-10 | ENSRNOT00000009573 |
| Ube2b-202 | 10 | 31.35 | -2.16 | 2.88E-05 | ENSRNOT00000106084 |
| Rad50-202 | 10 | 7.55 | -2.08 | 9.33E-11 | ENSRNOT00000106947 |
| Slc22a4-201 | 10 | 5.24 | -3.15 | 0.02 | ENSRNOT00000058907 |
| Fnip1-202 | 10 | 10.47 | -2.32 | 5.55E-04 | ENSRNOT00000115556 |
| Olr1433-201 | 10 | 7.63 | -2.12 | 0.02 | ENSRNOT00000058503 |
| Btnl10-201 | 10 | 5.17 | 3.34 | 4.91E-03 | ENSRNOT00000048642 |
| Wnt9a-201 | 10 | 10.75 | -3.66 | 2.89E-12 | ENSRNOT00000058327 |
| Rasd1-201 | 10 | 109.13 | 2.28 | 6.42E-24 | ENSRNOT00000004475 |
| Pemt-203 | 10 | 24.13 | -2.15 | 0.03 | ENSRNOT00000112526 |
| Pmp22-201 | 10 | 21.24 | -8.91 | 9.02E-06 | ENSRNOT00000041606 |

| | | | | | |
|---|---|---|---|---|---|
| Myh10-201 | 10 | 40.01 | 2.01 | 4.68E-13 | ENSRNOT00000065895 |
| Ndel1-202 | 10 | 10.06 | 2.39 | 0.02 | ENSRNOT00000065505 |
| Chd3-204 | 10 | 9.2 | 2.73 | 3.67E-11 | ENSRNOT00000114886 |
| Atp1b2-201 | 10 | 34.5 | 2.67 | 1.51E-15 | ENSRNOT00000015076 |
| Plscr3-201 | 10 | 29.64 | 2.21 | 1.45E-07 | ENSRNOT00000031640 |
| Pitpnm3-201 | 10 | 11.33 | -2.15 | 1.00E-09 | ENSRNOT00000071764 |
| Smtnl2-201 | 10 | 16.17 | -2.27 | 1.17E-06 | ENSRNOT00000020379 |
| P2rx1-202 | 10 | 9.17 | -413.86 | 5.56E-06 | ENSRNOT00000088245 |
| Itgae-201 | 10 | 24.05 | -2.33 | 4.42E-15 | ENSRNOT00000073209 |
| Sgsm2-203 | 10 | 11.24 | 2.3 | 2.24E-05 | ENSRNOT00000092314 |
| Hic1-201 | 10 | 12.52 | 3.2 | 6.24E-09 | ENSRNOT00000004143 |
| Vtn-201 | 10 | 7.52 | 4.41 | 2.70E-03 | ENSRNOT00000039954 |
| Ksr1-203 | 10 | 6.68 | 2.9 | 1.16E-04 | ENSRNOT00000104397 |
| Slfn5-202 | 10 | 18.2 | 2.26 | 7.06E-11 | ENSRNOT00000076445 |
| Wfdc21-201 | 10 | 31.11 | -3.52 | 0.02 | ENSRNOT00000042643 |
| Ggnbp2-202 | 10 | 12.82 | -2.05 | 9.69E-03 | ENSRNOT00000085728 |
| Cltc-203 | 10 | 10.44 | 3.07 | 3.77E-06 | ENSRNOT00000098805 |
| Ypel2-202 | 10 | 8.05 | 19.01 | 1.27E-08 | ENSRNOT00000114367 |
| Gdpd1-203 | 10 | 52.45 | -2.05 | 6.28E-19 | ENSRNOT00000109310 |
| Gdpd1-201 | 10 | 38.58 | -2.28 | 6.63E-10 | ENSRNOT00000079311 |
| Msi2-203 | 10 | 11.81 | 2.64 | 0.05 | ENSRNOT00000106986 |
| Mmd-201 | 10 | 14.13 | -4.59 | 5.80E-08 | ENSRNOT00000003308 |
| Tob1-201 | 10 | 26.34 | 3.57 | 1.89E-08 | ENSRNOT00000003780 |
| Kat7-204 | 10 | 6.46 | 340.73 | 1.96E-08 | ENSRNOT00000109616 |
| Cdk5rap3-203 | 10 | 9.12 | -3.45 | 7.50E-04 | ENSRNOT00000097303 |
| Arhgap23-203 | 10 | 8.71 | 2.83 | 1.33E-03 | ENSRNOT00000100892 |
| Cdk12-205 | 10 | 5.63 | -3.3 | 0.02 | ENSRNOT00000038939 |
| Krt16-203 | 10 | 311.88 | -3.57 | 2.64E-89 | ENSRNOT00000019133 |
| Hsd17b1-201 | 10 | 342.77 | 6.45 | 9.23E-169 | ENSRNOT00000026878 |
| Mlx-204 | 10 | 9.31 | 1,091.39 | 2.18E-03 | ENSRNOT00000119770 |
| Dhx8-201 | 10 | 5.39 | 2.1 | 0.03 | ENSRNOT00000048140 |
| Nsf-201 | 10 | 31.68 | 3.61 | 2.55E-10 | ENSRNOT00000006361 |
| Ace-201 | 10 | 20.74 | -2.48 | 1.85E-15 | ENSRNOT00000010627 |
| Ace-202 | 10 | 10.53 | -4.83 | 1.95E-07 | ENSRNOT00000092961 |
| Fads6-201 | 10 | 19.15 | 3.53 | 3.23E-08 | ENSRNOT00000029001 |
| Cdr2l-201 | 10 | 23.43 | 2.11 | 1.69E-16 | ENSRNOT00000035865 |
| St6galnac2-203 | 10 | 6.13 | -3.88 | 7.86E-04 | ENSRNOT00000093828 |
| St6galnac2-201 | 10 | 25.17 | -2.44 | 1.54E-09 | ENSRNOT00000016622 |
| Septin9-204 | 10 | 13.63 | -2.66 | 2.34E-07 | ENSRNOT00000113646 |
| Septin9-203 | 10 | 24.91 | 2.07 | 6.99E-12 | ENSRNOT00000100054 |
| Tnrc6c-202 | 10 | 12.02 | 2.96 | 6.09E-06 | ENSRNOT00000106642 |
| Lgals3bp-202 | 10 | 9.7 | 245.52 | 1.73E-08 | ENSRNOT00000102908 |
| Actg1-201 | 10 | 164.43 | 2.6 | 2.11E-14 | ENSRNOT00000054976 |
| Nploc4-201 | 10 | 23.46 | 2.54 | 3.83E-08 | ENSRNOT00000054973 |
| P4hb-203 | 10 | 6.76 | -2.54 | 0.01 | ENSRNOT00000106240 |
| Stx19-201 | 11 | 7.25 | -2.07 | 1.27E-03 | ENSRNOT00000102767 |
| Robo1-203 | 11 | 15.57 | 2.92 | 1.93E-14 | ENSRNOT00000104951 |
| ENSRNOT00000100005 | 11 | 350.45 | -2.51 | 1.20E-16 | ENSRNOT00000100005 |
| Mrap-201 | 11 | 6.93 | 3.33 | 0.03 | ENSRNOT00000036027 |
| Smim11-202 | 11 | 10.75 | -2.23 | 6.15E-05 | ENSRNOT00000096936 |
| Sh3bgr-204 | 11 | 10.23 | -3.33 | 9.12E-03 | ENSRNOT00000119421 |

| | | | | | |
|---|---|---|---|---|---|
| Mx2-202 | 11 | 10.52 | -4.21 | 3.14E-07 | ENSRNOT00000087603 |
| Tfg-202 | 11 | 5.41 | -2.34 | 1.71E-03 | ENSRNOT00000088152 |
| Abi3bp-205 | 11 | 11.28 | -4.27 | 4.70E-05 | ENSRNOT00000063864 |
| Cd200-203 | 11 | 9.86 | -9.08 | 0.02 | ENSRNOT00000093295 |
| Golgb1-203 | 11 | 6.7 | -2.4 | 1.29E-16 | ENSRNOT00000106429 |
| Kpna1-203 | 11 | 6.48 | -2.02 | 1.60E-03 | ENSRNOT00000097087 |
| Tnk2-202 | 11 | 5.1 | -2.18 | 0.03 | ENSRNOT00000084768 |
| Pcyt1a-201 | 11 | 48.62 | 2.14 | 9.31E-35 | ENSRNOT00000002403 |
| Fam43a-201 | 11 | 56.8 | 2.56 | 1.99E-30 | ENSRNOT00000002355 |
| Masp1-203 | 11 | 14.27 | 3.11 | 4.91E-10 | ENSRNOT00000094154 |
| Masp1-204 | 11 | 120.39 | 2.62 | 5.44E-53 | ENSRNOT00000111260 |
| Dnajb11-204 | 11 | 9.34 | 6.79 | 1.77E-03 | ENSRNOT00000116132 |
| Dgcr2-203 | 11 | 10.88 | -2.48 | 8.41E-05 | ENSRNOT00000107429 |
| Med15-204 | 11 | 7.45 | 2.8 | 6.76E-03 | ENSRNOT00000117596 |
| Arhgef18-203 | 12 | 5.08 | 3.21 | 0.02 | ENSRNOT00000110862 |
| LOC120095871-202 | 12 | 18.18 | -2.14 | 7.98E-08 | ENSRNOT00000115928 |
| LOC304239-201 | 12 | 5.53 | -3.87 | 7.67E-06 | ENSRNOT00000076491 |
| Smurf1-203 | 12 | 6.36 | 2.26 | 7.72E-03 | ENSRNOT00000084577 |
| Fscn1-203 | 12 | 28.41 | 2.82 | 2.64E-08 | ENSRNOT00000102873 |
| Actb-204 | 12 | 5.6 | -641.56 | 3.57E-03 | ENSRNOT00000116486 |
| Sun1-206 | 12 | 6.32 | 2.7 | 0.03 | ENSRNOT00000113934 |
| Rpl31l4-201 | 12 | 516.05 | -3.07 | 1.92E-59 | ENSRNOT00000045401 |
| Stag3-201 | 12 | 9.94 | -3.33 | 1.35E-12 | ENSRNOT00000042006 |
| Irs3-201 | 12 | 9.48 | 5.33 | 4.97E-05 | ENSRNOT00000001874 |
| Cldn15-201 | 12 | 26.03 | -3.17 | 7.14E-12 | ENSRNOT00000001923 |
| Col26a1-201 | 12 | 11.54 | -3.31 | 2.37E-09 | ENSRNOT00000080068 |
| Upk3b-201 | 12 | 42.02 | -4.01 | 1.29E-14 | ENSRNOT00000037639 |
| Mdh2-202 | 12 | 28.28 | -612.83 | 5.65E-07 | ENSRNOT00000112030 |
| Por-203 | 12 | 75.74 | 2 | 8.02E-23 | ENSRNOT00000101964 |
| Trim50-201 | 12 | 32.59 | -2.02 | 4.68E-08 | ENSRNOT00000032250 |
| Gtf2i-205 | 12 | 19.82 | 2.27 | 1.17E-08 | ENSRNOT00000098927 |
| Gusb-201 | 12 | 53.37 | 2.07 | 4.04E-17 | ENSRNOT00000001215 |
| Scarb1-204 | 12 | 39.63 | 2.12 | 2.09E-07 | ENSRNOT00000110392 |
| Scarb1-201 | 12 | 250.84 | 2.39 | 1.52E-20 | ENSRNOT00000001299 |
| Rflna-201 | 12 | 12.24 | -2.41 | 7.22E-03 | ENSRNOT00000119390 |
| Rilpl1-202 | 12 | 16.62 | 2.24 | 4.14E-05 | ENSRNOT00000090831 |
| Clip1-206 | 12 | 5.53 | 524.86 | 1.73E-07 | ENSRNOT00000098458 |
| Cfap251-201 | 12 | 16.06 | -2.27 | 2.01E-05 | ENSRNOT00000001813 |
| Hpd-202 | 12 | 11.47 | -2.55 | 0.01 | ENSRNOT00000120179 |
| Oas1a-201 | 12 | 47.47 | 2.04 | 5.47E-07 | ENSRNOT00000038426 |
| Rasal1-201 | 12 | 7.51 | -2.09 | 7.97E-03 | ENSRNOT00000001856 |
| Tesc-203 | 12 | 5.65 | -59.41 | 2.81E-03 | ENSRNOT00000104891 |
| Wsb2-201 | 12 | 11.91 | 2.13 | 0.04 | ENSRNOT00000001499 |
| Hspb8-201 | 12 | 11.75 | -2.71 | 9.08E-04 | ENSRNOT00000039275 |
| Oasl-201 | 12 | 7.76 | 2.32 | 0.05 | ENSRNOT00000001570 |
| Trpv4-201 | 12 | 8.27 | 3.21 | 0.01 | ENSRNOT00000001586 |
| Ttc28-201 | 12 | 8.84 | 2.74 | 2.50E-13 | ENSRNOT00000046920 |
| Inhbb-201 | 13 | 607.84 | 2.78 | 4.33E-68 | ENSRNOT00000086350 |
| Actr3-201 | 13 | 29.64 | -42.76 | 1.89E-34 | ENSRNOT00000004520 |
| Slc41a1-201 | 13 | 29.77 | 2.04 | 4.18E-19 | ENSRNOT00000063882 |
| Lgr6-201 | 13 | 12.71 | 2.57 | 2.53E-16 | ENSRNOT00000006969 |

| | | | | | |
|---|---|---|---|---|---|
| Camsap2-203 | 13 | 7.12 | -7.85 | 1.70E-05 | ENSRNOT00000098757 |
| Camsap2-201 | 13 | 9.72 | 3.24 | 5.01E-05 | ENSRNOT00000060410 |
| Nr5a2-202 | 13 | 123.63 | 2.12 | 2.27E-33 | ENSRNOT00000097279 |
| Lhx9-201 | 13 | 28.68 | -5.13 | 1.92E-26 | ENSRNOT00000014218 |
| Lhx9-203 | 13 | 6.82 | -5.23 | 8.51E-06 | ENSRNOT00000116473 |
| Cfh-201 | 13 | 16.84 | 2.75 | 8.68E-16 | ENSRNOT00000060111 |
| Arpc5-204 | 13 | 12.24 | 1,372.79 | 8.10E-05 | ENSRNOT00000107757 |
| Glul-202 | 13 | 70.95 | 2.47 | 4.14E-21 | ENSRNOT00000118700 |
| Xpr1-201 | 13 | 6.55 | 8.1 | 4.86E-05 | ENSRNOT00000000049 |
| Xpr1-205 | 13 | 6.34 | -2.95 | 0.04 | ENSRNOT00000113923 |
| Myoc-201 | 13 | 11.22 | -3.74 | 1.10E-05 | ENSRNOT00000004386 |
| Gorab-202 | 13 | 5.31 | -3.1 | 1.76E-03 | ENSRNOT00000106402 |
| Styxl2-201 | 13 | 7.81 | 3.38 | 4.85E-06 | ENSRNOT00000004965 |
| Mael-201 | 13 | 71.77 | -3.44 | 9.60E-48 | ENSRNOT00000005060 |
| Ildr2-204 | 13 | 6.62 | -3.86 | 6.53E-04 | ENSRNOT00000072897 |
| Pbx1-202 | 13 | 18.13 | 2.15 | 2.64E-09 | ENSRNOT00000103202 |
| Nuf2-203 | 13 | 12.01 | -2.04 | 6.55E-04 | ENSRNOT00000113324 |
| Olfml2b-201 | 13 | 33.94 | 2.42 | 4.14E-21 | ENSRNOT00000004117 |
| Fcrla-201 | 13 | 17.17 | -4.02 | 1.30E-05 | ENSRNOT00000004183 |
| B4galt3-202 | 13 | 5.87 | -11.13 | 6.62E-04 | ENSRNOT00000108060 |
| Ly9-202 | 13 | 13.87 | 3.08 | 6.51E-13 | ENSRNOT00000096361 |
| Copa-204 | 13 | 12.96 | 2.07 | 2.83E-05 | ENSRNOT00000113779 |
| Capn8-202 | 13 | 17.03 | -2.76 | 9.25E-09 | ENSRNOT00000067005 |
| Capn8-201 | 13 | 16.21 | -2.79 | 9.89E-08 | ENSRNOT00000004649 |
| Susd4-201 | 13 | 44.12 | -2.93 | 1.04E-25 | ENSRNOT00000052407 |
| Mark1-202 | 13 | 45.1 | 2.27 | 3.47E-17 | ENSRNOT00000080309 |
| Mark1-201 | 13 | 61.93 | 2.06 | 1.54E-16 | ENSRNOT00000003198 |
| LOC102554605-202 | 14 | 8.14 | -2.23 | 0.04 | ENSRNOT00000099719 |
| Tmed5-201 | 14 | 8.08 | -4.96 | 0.03 | ENSRNOT00000000083 |
| Cdc7-203 | 14 | 5.37 | 6.79 | 6.23E-04 | ENSRNOT00000101926 |
| Zfp326-204 | 14 | 11.49 | -2.16 | 0.02 | ENSRNOT00000095214 |
| Gbp6-205 | 14 | 24.4 | -2.03 | 1.98E-20 | ENSRNOT00000109379 |
| Abcg3l3-201 | 14 | 15.09 | -2.37 | 4.19E-09 | ENSRNOT00000093929 |
| Thoc2l-203 | 14 | 7.83 | -2.02 | 3.99E-21 | ENSRNOT00000102766 |
| Hpse-201 | 14 | 5.28 | 2.28 | 0.04 | ENSRNOT00000002983 |
| Sec31a-201 | 14 | 11.56 | 2.34 | 3.77E-06 | ENSRNOT00000003072 |
| Anxa3-202 | 14 | 12.52 | -2.11 | 1.19E-03 | ENSRNOT00000068134 |
| Fras1-202 | 14 | 5.61 | -2.06 | 9.51E-11 | ENSRNOT00000100214 |
| Cxcl9-201 | 14 | 26.17 | 2.82 | 1.08E-05 | ENSRNOT00000003082 |
| Cdkl2-203 | 14 | 5.3 | -2.19 | 8.48E-03 | ENSRNOT00000079304 |
| Alb-201 | 14 | 5.88 | 5.84 | 0.02 | ENSRNOT00000003921 |
| Slc4a4-202 | 14 | 7.09 | -2.37 | 0.04 | ENSRNOT00000033806 |
| Slc4a4-203 | 14 | 6.43 | -3.3 | 2.00E-03 | ENSRNOT00000095074 |
| Gnrhr-201 | 14 | 40.45 | 6.21 | 5.32E-30 | ENSRNOT00000002755 |
| LOC108348180-201 | 14 | 12.35 | -2.07 | 7.21E-04 | ENSRNOT00000002862 |
| Uchl1-201 | 14 | 20.77 | -7.21 | 6.03E-05 | ENSRNOT00000003248 |
| Sod3-201 | 14 | 34.92 | 2.25 | 3.05E-11 | ENSRNOT00000005155 |
| Cpz-201 | 14 | 38.51 | 2.22 | 1.26E-12 | ENSRNOT00000012109 |
| Rgs12-202 | 14 | 7.04 | 2.29 | 4.17E-04 | ENSRNOT00000045068 |
| Add1-202 | 14 | 19.41 | 2.48 | 5.20E-06 | ENSRNOT00000018340 |
| Pik3ip1-201 | 14 | 6.52 | -2.11 | 0.03 | ENSRNOT00000057738 |

| | | | | | |
|---|---|---|---|---|---|
| Hormad2-202 | 14 | 12.21 | -2.25 | 5.75E-03 | ENSRNOT00000080078 |
| Ewsr1-205 | 14 | 11.98 | 4.02 | 2.70E-03 | ENSRNOT00000094261 |
| Ankrd36-202 | 14 | 5.48 | -4.26 | 3.27E-03 | ENSRNOT00000083756 |
| Ccdc201-201 | 14 | 120.89 | -3.92 | 1.00E-36 | ENSRNOT00000116147 |
| Egfr-201 | 14 | 14.09 | 2.3 | 5.11E-09 | ENSRNOT00000006087 |
| Etaa1l1-202 | 14 | 15.78 | -2.2 | 7.53E-10 | ENSRNOT00000097260 |
| Sanbr-203 | 14 | 7.59 | -2.02 | 4.24E-03 | ENSRNOT00000091143 |
| Plau-201 | 15 | 37.26 | 2.5 | 9.82E-20 | ENSRNOT00000014273 |
| Atxn7-201 | 15 | 14.75 | -2.72 | 1.06E-08 | ENSRNOT00000010103 |
| Ptprg-205 | 15 | 6.96 | 3.4 | 0.02 | ENSRNOT00000104654 |
| Ccnb1ip1-201 | 15 | 9.69 | -2.96 | 1.27E-03 | ENSRNOT00000029488 |
| Arhgef40-201 | 15 | 6.32 | 2.71 | 9.80E-04 | ENSRNOT00000078475 |
| Cma1-201 | 15 | 14.92 | -4.35 | 3.23E-03 | ENSRNOT00000067539 |
| Mcpt9-205 | 15 | 9.82 | -3.9 | 0.03 | ENSRNOT00000099690 |
| Mcpt1-201 | 15 | 9.93 | -8.51 | 1.11E-04 | ENSRNOT00000091757 |
| Rnf17-201 | 15 | 17.06 | -2.15 | 1.36E-14 | ENSRNOT00000010795 |
| Cab39l-204 | 15 | 7.16 | -2.63 | 0.01 | ENSRNOT00000103841 |
| Sox7-201 | 15 | 8.48 | 2.78 | 1.50E-04 | ENSRNOT00000016503 |
| Ints9-203 | 15 | 9.75 | 2.14 | 0.03 | ENSRNOT00000096924 |
| Dpysl2-202 | 15 | 27.64 | -2.26 | 9.24E-07 | ENSRNOT00000103229 |
| Stc1-201 | 15 | 58.54 | 3.23 | 2.11E-26 | ENSRNOT00000020728 |
| Piwil2-201 | 15 | 15.57 | -2.73 | 6.17E-18 | ENSRNOT00000013284 |
| Gfra2-202 | 15 | 12.64 | 2.2 | 7.09E-05 | ENSRNOT00000094535 |
| Itm2b-202 | 15 | 6.62 | -2.9 | 1.13E-04 | ENSRNOT00000106965 |
| Dct-201 | 15 | 9.35 | -2.73 | 1.33E-03 | ENSRNOT00000076707 |
| Dzip1-201 | 15 | 9.69 | -2.57 | 2.02E-04 | ENSRNOT00000038596 |
| Clybl-202 | 15 | 60.84 | -3.5 | 2.47E-24 | ENSRNOT00000107907 |
| Pcca-203 | 15 | 10.31 | -2.01 | 0.04 | ENSRNOT00000094922 |
| Rps24-203 | 16 | 244.04 | -2.45 | 7.27E-23 | ENSRNOT00000101331 |
| Il17rd-201 | 16 | 8.62 | -2.43 | 4.94E-06 | ENSRNOT00000030102 |
| Nisch-201 | 16 | 11.79 | 27.72 | 7.79E-41 | ENSRNOT00000036910 |
| Tnnc1-201 | 16 | 25.37 | -2.72 | 2.80E-04 | ENSRNOT00000025606 |
| Phf7-203 | 16 | 5.11 | -2.69 | 0.01 | ENSRNOT00000102974 |
| Ankrd28-202 | 16 | 7.52 | 3.71 | 3.30E-07 | ENSRNOT00000094533 |
| Mmrn2-201 | 16 | 7.49 | 2.12 | 4.44E-04 | ENSRNOT00000084422 |
| Isyna1-201 | 16 | 894.52 | 2.04 | 9.23E-26 | ENSRNOT00000026821 |
| LOC102551257-202 | 16 | 9.07 | -2.01 | 1.79E-10 | ENSRNOT00000120042 |
| Ddx60-203 | 16 | 5.88 | -3.08 | 3.37E-09 | ENSRNOT00000088707 |
| Hpgd-201 | 16 | 48.16 | 2.86 | 5.45E-21 | ENSRNOT00000014229 |
| Tenm3-201 | 16 | 8.54 | 2.35 | 4.07E-14 | ENSRNOT00000017624 |
| Enpp6-201 | 16 | 5.52 | 4.18 | 0.01 | ENSRNOT00000013005 |
| Tlr3-202 | 16 | 5.75 | -2.36 | 0.01 | ENSRNOT00000087986 |
| Fat1-201 | 16 | 47.09 | 2.52 | 3.76E-27 | ENSRNOT00000067486 |
| Pcm1-201 | 16 | 11.35 | 3.24 | 4.67E-08 | ENSRNOT00000014202 |
| Pcm1-202 | 16 | 6.23 | -4.21 | 0.02 | ENSRNOT00000085435 |
| Zdhhc2-201 | 16 | 16.25 | 2.31 | 9.74E-03 | ENSRNOT00000036092 |
| Nek3-201 | 16 | 7.46 | -2.27 | 0.04 | ENSRNOT00000017106 |
| Myom2-202 | 16 | 11.49 | 9.59 | 1.06E-08 | ENSRNOT00000097431 |
| Cntnap3-201 | 17 | 14.24 | 2.2 | 2.59E-10 | ENSRNOT00000046525 |
| Fbp2-201 | 17 | 12.32 | -3.64 | 6.99E-04 | ENSRNOT00000023865 |
| LOC102552000-203 | 17 | 6.5 | -2.75 | 1.11E-07 | ENSRNOT00000118981 |

| | | | | | |
|---|---|---|---|---|---|
| LOC102552000-205 | 17 | 15.58 | -2.67 | 3.65E-20 | ENSRNOT00000103626 |
| ENSRNOT00000109753 | 17 | 8.77 | -2.06 | 2.09E-04 | ENSRNOT00000109753 |
| Phf2-202 | 17 | 20.15 | 2.1 | 1.15E-03 | ENSRNOT00000087692 |
| Phf2-201 | 17 | 5.13 | -2.81 | 0.02 | ENSRNOT00000022669 |
| Phactr1-203 | 17 | 5.07 | 2.35 | 0.03 | ENSRNOT00000107421 |
| Sycp2l-201 | 17 | 22.15 | -4.36 | 1.06E-22 | ENSRNOT00000038180 |
| Fars2-205 | 17 | 12.21 | 2.98 | 1.94E-03 | ENSRNOT00000113971 |
| Prpf4b-201 | 17 | 25.39 | -2.2 | 1.76E-03 | ENSRNOT00000022902 |
| Serpinb9-204 | 17 | 11.89 | -1,833.40 | 4.26E-03 | ENSRNOT00000115382 |
| Serpinb9-203 | 17 | 5.39 | 123.85 | 2.19E-04 | ENSRNOT00000111970 |
| Serpinb1a-201 | 17 | 44.05 | -4.03 | 5.84E-18 | ENSRNOT00000059854 |
| Dusp22-201 | 17 | 6.11 | -3.7 | 0.02 | ENSRNOT00000090946 |
| Carmil1-205 | 17 | 7 | -2.41 | 1.70E-08 | ENSRNOT00000117749 |
| Carmil1-204 | 17 | 6.79 | 34.5 | 2.25E-13 | ENSRNOT00000108358 |
| Btn1a1-202 | 17 | 13.5 | -4.07 | 4.59E-15 | ENSRNOT00000105103 |
| Btn1a1-201 | 17 | 10.24 | -2.36 | 4.81E-05 | ENSRNOT00000023622 |
| ENSRNOT00000097598 | 17 | 5.36 | -2.07 | 0.04 | ENSRNOT00000097598 |
| Elmo1-203 | 17 | 9.47 | 2.95 | 9.16E-09 | ENSRNOT00000090663 |
| Inhba-201 | 17 | 391.29 | 4.64 | 4.23E-85 | ENSRNOT00000019272 |
| RGD1308147-201 | 17 | 14.54 | -2.05 | 1.34E-03 | ENSRNOT00000020794 |
| Crem-210 | 17 | 6.1 | -1,609.51 | 5.85E-03 | ENSRNOT00000115950 |
| Mkx-201 | 17 | 11.53 | 2.06 | 1.19E-04 | ENSRNOT00000025623 |
| Pfkp-201 | 17 | 47.07 | 2.98 | 8.00E-28 | ENSRNOT00000023252 |
| Asb13-202 | 17 | 8.1 | 3.85 | 6.00E-03 | ENSRNOT00000103759 |
| Ccdc3-201 | 17 | 23.36 | 2.93 | 8.19E-20 | ENSRNOT00000024110 |
| Nmt2-205 | 17 | 12.65 | -3.47 | 2.24E-03 | ENSRNOT00000119939 |
| Nmt2-204 | 17 | 7.33 | -2.45 | 7.11E-05 | ENSRNOT00000112511 |
| Etl4-202 | 17 | 15.81 | 3.6 | 2.66E-25 | ENSRNOT00000096341 |
| Cmc4-201 | 18 | 49.13 | -2.15 | 5.24E-05 | ENSRNOT00000084138 |
| Npc1-201 | 18 | 11.5 | 3.3 | 4.91E-06 | ENSRNOT00000016167 |
| Cdh2-204 | 18 | 5.07 | 322.52 | 6.57E-08 | ENSRNOT00000118601 |
| Cdh2-201 | 18 | 14.89 | 2.43 | 7.13E-08 | ENSRNOT00000021170 |
| Dtna-202 | 18 | 17.04 | -4.62 | 2.59E-25 | ENSRNOT00000097179 |
| Fhod3-204 | 18 | 17.76 | 3.48 | 6.46E-15 | ENSRNOT00000093285 |
| Egr1-201 | 18 | 52.87 | 2.33 | 9.78E-05 | ENSRNOT00000026303 |
| Hspa9-202 | 18 | 47.32 | -10.33 | 1.41E-06 | ENSRNOT00000079460 |
| Cystm1-203 | 18 | 16.04 | -2.49 | 0.02 | ENSRNOT00000094556 |
| Cystm1-201 | 18 | 20.7 | -2.13 | 0.03 | ENSRNOT00000025495 |
| Zmat2-205 | 18 | 9.54 | 2.08 | 0.04 | ENSRNOT00000104252 |
| Zmat2-203 | 18 | 6.71 | -150.5 | 1.40E-09 | ENSRNOT00000095646 |
| Pcdha5-201 | 18 | 12.04 | 2.8 | 1.64E-17 | ENSRNOT00000027340 |
| Tnfaip8-201 | 18 | 11.66 | -2.07 | 0.01 | ENSRNOT00000021972 |
| Prdm6-201 | 18 | 7.52 | 2.85 | 3.72E-03 | ENSRNOT00000047755 |
| Fbn2-201 | 18 | 66.6 | 2.09 | 8.92E-33 | ENSRNOT00000066548 |
| Adamts19-201 | 18 | 42.33 | -3.06 | 4.23E-40 | ENSRNOT00000026543 |
| Synpo-201 | 18 | 13.82 | 2.26 | 3.36E-06 | ENSRNOT00000025934 |
| Tcof1-201 | 18 | 19.76 | 5.65 | 2.12E-15 | ENSRNOT00000045041 |
| Pdgfrb-202 | 18 | 5.95 | -30.4 | 1.04E-06 | ENSRNOT00000078764 |
| Grpel2-204 | 18 | 7.17 | -2.06 | 2.30E-06 | ENSRNOT00000117207 |
| Txnl1-204 | 18 | 19.99 | -1,405.49 | 7.00E-04 | ENSRNOT00000100860 |
| Ccbe1-201 | 18 | 18 | 2.4 | 3.62E-19 | ENSRNOT00000116155 |

| | | | | | |
|---|---|---|---|---|---|
| Lipg-201 | 18 | 14.34 | 2 | 2.06E-03 | ENSRNOT00000025257 |
| Terb1-201 | 19 | 32.18 | -2.41 | 5.46E-28 | ENSRNOT00000087469 |
| Cdh5-202 | 19 | 42.36 | 2.33 | 6.62E-31 | ENSRNOT00000086995 |
| Cdh5-201 | 19 | 8.84 | 2.08 | 0.03 | ENSRNOT00000017983 |
| Ces1d-201 | 19 | 5.48 | -10.28 | 1.49E-04 | ENSRNOT00000024187 |
| Irx5-202 | 19 | 7.32 | -2.46 | 6.02E-03 | ENSRNOT00000089975 |
| Irx3-201 | 19 | 23.73 | -4.03 | 1.44E-26 | ENSRNOT00000015583 |
| Chd9-202 | 19 | 5.79 | 2.35 | 6.73E-07 | ENSRNOT00000076138 |
| Cbln1-201 | 19 | 6.11 | 3.02 | 0.01 | ENSRNOT00000000011 |
| Gpt2-205 | 19 | 9.33 | 2.06 | 0.05 | ENSRNOT00000115000 |
| ENSRNOT00000049860 | 19 | 351.06 | -2.55 | 2.67E-52 | ENSRNOT00000049860 |
| LOC685989-202 | 19 | 186.94 | -4.67 | 6.67E-104 | ENSRNOT00000100796 |
| LOC685989-203 | 19 | 142.79 | -4.04 | 1.42E-73 | ENSRNOT00000115454 |
| LOC102554391-201 | 19 | 8.37 | -3.86 | 0.01 | ENSRNOT00000119060 |
| RGD1562660-201 | 19 | 146.96 | -2.84 | 3.65E-31 | ENSRNOT00000044160 |
| ENSRNOT00000112343 | 19 | 76.33 | -4.99 | 9.30E-40 | ENSRNOT00000112343 |
| ENSRNOT00000114003 | 19 | 104.77 | -3.16 | 6.43E-29 | ENSRNOT00000114003 |
| ENSRNOT00000108310 | 19 | 191.29 | -4.2 | 2.15E-86 | ENSRNOT00000108310 |
| ENSRNOT00000040545 | 19 | 25.11 | -5.02 | 2.24E-07 | ENSRNOT00000040545 |
| Farsa-202 | 19 | 13.77 | -14.62 | 5.52E-06 | ENSRNOT00000004404 |
| Podnl1-201 | 19 | 6.72 | -2.99 | 7.55E-03 | ENSRNOT00000037453 |
| Adgrl1-206 | 19 | 5.61 | -2.63 | 3.56E-06 | ENSRNOT00000103294 |
| Hhip-201 | 19 | 32.81 | 3.5 | 5.67E-29 | ENSRNOT00000024616 |
| Arhgap10-201 | 19 | 7.21 | 2.79 | 0.01 | ENSRNOT00000017867 |
| Smpd3-201 | 19 | 5.17 | 19.45 | 8.12E-10 | ENSRNOT00000031977 |
| LOC100360573-201 | 19 | 31.83 | -3.68 | 3.50E-03 | ENSRNOT00000027225 |
| Zfp612-202 | 19 | 13.81 | -2.48 | 3.56E-27 | ENSRNOT00000112604 |
| Calb2-201 | 19 | 9.68 | -3.11 | 0.02 | ENSRNOT00000022943 |
| Cmip-203 | 19 | 8.05 | 2.63 | 2.69E-03 | ENSRNOT00000112862 |
| Hsd17b2-201 | 19 | 17.58 | -4.12 | 2.21E-06 | ENSRNOT00000018795 |
| Osgin1-201 | 19 | 7.68 | 2.99 | 0.05 | ENSRNOT00000020225 |
| Adad2-201 | 19 | 10.41 | -2.24 | 0.01 | ENSRNOT00000021072 |
| Crispld2-202 | 19 | 48.33 | 2.49 | 4.14E-28 | ENSRNOT00000087857 |
| RGD1304884-202 | 19 | 11.83 | 2.05 | 1.00E-10 | ENSRNOT00000099058 |
| Slc7a5-201 | 19 | 53.88 | 2.31 | 7.62E-25 | ENSRNOT00000025784 |
| Cbfa2t3-201 | 19 | 7.24 | 3.33 | 1.58E-03 | ENSRNOT00000019820 |
| Tubb3-201 | 19 | 12.24 | 3.53 | 1.26E-04 | ENSRNOT00000023452 |
| Nrp1-201 | 19 | 12.57 | 2.86 | 3.79E-10 | ENSRNOT00000014492 |
| ENSRNOT00000118146 | 20 | 19.73 | -2.02 | 1.81E-07 | ENSRNOT00000118146 |
| RT1-CE16-201 | 20 | 22.22 | -4.1 | 1.77E-22 | ENSRNOT00000049310 |
| RT1-A2-201 | 20 | 23.58 | -2.39 | 7.37E-10 | ENSRNOT00000082497 |
| Ly6g6e-202 | 20 | 31.82 | -2.01 | 7.65E-05 | ENSRNOT00000113254 |
| Neu1-201 | 20 | 17.16 | 2.07 | 1.21E-04 | ENSRNOT00000049344 |
| Slc44a4-201 | 20 | 6.07 | 2.51 | 0.04 | ENSRNOT00000001174 |
| Cfb-201 | 20 | 10.28 | -2.6 | 3.23E-04 | ENSRNOT00000000477 |
| C4a-202 | 20 | 16.62 | -2.11 | 1.66E-11 | ENSRNOT00000086027 |
| Ppt2-203 | 20 | 11.6 | 3.14 | 3.63E-03 | ENSRNOT00000083489 |
| Snrpc-203 | 20 | 69.58 | 2.05 | 3.04E-05 | ENSRNOT00000115940 |
| Tcp11-201 | 20 | 7.9 | -3.64 | 5.14E-03 | ENSRNOT00000045044 |
| Scube3-202 | 20 | 10.57 | 2.22 | 1.37E-04 | ENSRNOT00000100675 |
| Kctd20-203 | 20 | 5.38 | -4.4 | 2.07E-03 | ENSRNOT00000105827 |

| | | | | | |
|---|---|---|---|---|---|
| Col6a2-202 | 20 | 67.24 | 2.59 | 3.85E-42 | ENSRNOT00000097191 |
| Col6a2-201 | 20 | 328.12 | 2.08 | 1.89E-33 | ENSRNOT00000001695 |
| Dip2a-202 | 20 | 7.21 | 2.98 | 1.89E-03 | ENSRNOT00000083878 |
| Chchd10-201 | 20 | 88.56 | 2.35 | 3.44E-15 | ENSRNOT00000031472 |
| Ube2d1-201 | 20 | 19.86 | 2.23 | 0.04 | ENSRNOT00000000750 |
| Fam13c-203 | 20 | 14.53 | 2.19 | 5.55E-07 | ENSRNOT00000118972 |
| Oit3-202 | 20 | 9.13 | -3.26 | 3.19E-03 | ENSRNOT00000112351 |
| Unc5b-203 | 20 | 8.89 | 2.16 | 1.77E-05 | ENSRNOT00000109345 |
| Gja1-202 | 20 | 67.41 | 3.33 | 9.26E-59 | ENSRNOT00000100494 |
| Gja1-201 | 20 | 511.79 | 2.82 | 1.93E-64 | ENSRNOT00000001054 |
| Atg5-201 | 20 | 8.24 | 3.04 | 9.71E-03 | ENSRNOT00000057078 |
| LOC100363372-203 | X | 10.33 | -2.03 | 5.65E-04 | ENSRNOT00000119696 |
| Kdm6a-205 | X | 8.03 | 3.42 | 2.25E-04 | ENSRNOT00000109873 |
| Bcor-203 | X | 5.33 | 3.17 | 7.22E-05 | ENSRNOT00000116761 |
| Mid1ip1-201 | X | 193.06 | 2.09 | 7.86E-21 | ENSRNOT00000004305 |
| Mid1ip1-202 | X | 90.13 | 2.3 | 4.50E-13 | ENSRNOT00000095850 |
| RGD1561661-201 | X | 10.52 | -6.26 | 2.15E-08 | ENSRNOT00000030170 |
| Cybb-201 | X | 12.38 | 2.18 | 3.73E-05 | ENSRNOT00000038994 |
| Hdac6-203 | X | 13.11 | 2.92 | 3.41E-05 | ENSRNOT00000116471 |
| Ccnb3-202 | X | 10.19 | -8.53 | 6.07E-38 | ENSRNOT00000094271 |
| Ezhip-201 | X | 20.29 | 2.66 | 1.73E-07 | ENSRNOT00000060213 |
| Fam104b-202 | X | 22.13 | -2.01 | 2.23E-07 | ENSRNOT00000093387 |
| Gpr143-201 | X | 5.22 | -4.19 | 7.69E-03 | ENSRNOT00000004758 |
| Egfl6-202 | X | 31.84 | -3.79 | 1.90E-26 | ENSRNOT00000087621 |
| Sat1-202 | X | 192.34 | -3.22 | 9.17E-45 | ENSRNOT00000086311 |
| Sat1-201 | X | 172.09 | -3.26 | 1.67E-44 | ENSRNOT00000068078 |
| Mageb2-201 | X | 8.68 | -3.59 | 5.06E-05 | ENSRNOT00000050661 |
| AABR07038554.1-201 | X | 7.29 | -4.61 | 0.02 | ENSRNOT00000029387 |
| Msn-201 | X | 33.79 | 2.08 | 1.37E-07 | ENSRNOT00000076181 |
| Ogt-202 | X | 200.37 | -2.17 | 1.02E-54 | ENSRNOT00000082967 |
| Cxhxorf49-202 | X | 7.39 | -2.7 | 0.04 | ENSRNOT00000097163 |
| Ankra2-201 | X | 5.56 | 77.31 | 2.60E-04 | ENSRNOT00000022525 |
| Zfp711-202 | X | 5.48 | -2.92 | 0.03 | ENSRNOT00000095292 |
| Gprasp1-201 | X | 6.93 | -2.42 | 2.32E-08 | ENSRNOT00000073661 |
| Tceal8-201 | X | 7.95 | -176.89 | 3.13E-04 | ENSRNOT00000017138 |
| Morf4l2-207 | X | 5.74 | -2 | 5.83E-06 | ENSRNOT00000116026 |
| LOC108349244-201 | X | 17.31 | -2.21 | 6.48E-09 | ENSRNOT00000075912 |
| Agtr2-201 | X | 16.15 | -2.07 | 1.11E-04 | ENSRNOT00000074269 |
| Dock11-202 | X | 8.87 | -3.23 | 6.91E-03 | ENSRNOT00000080517 |
| Il13ra1-ps1-202 | X | 8.41 | -110.42 | 5.27E-16 | ENSRNOT00000112175 |
| Upf3b-202 | X | 11.33 | -2.63 | 3.46E-04 | ENSRNOT00000092574 |
| Rhox5-201 | X | 40.94 | 2.17 | 5.72E-04 | ENSRNOT00000092622 |
| Tmem255a-201 | X | 9.06 | 3.09 | 3.97E-04 | ENSRNOT00000036472 |
| Smarca1-202 | X | 7.92 | -2.18 | 0.04 | ENSRNOT00000114801 |
| Fam9b-201 | X | 15.92 | -3.19 | 4.77E-05 | ENSRNOT00000031927 |
| RGD1564845-201 | X | 13.09 | -3.76 | 2.40E-04 | ENSRNOT00000056700 |
| Map7d3-201 | X | 5.77 | -2.22 | 0.02 | ENSRNOT00000030445 |
| Arhgef6-201 | X | 11.55 | 2.74 | 9.79E-07 | ENSRNOT00000001158 |
| Rbmx-201 | X | 14.39 | 1,910.64 | 2.02E-03 | ENSRNOT00000001154 |
| Fam9c-201 | X | 14.16 | -4.41 | 8.30E-04 | ENSRNOT00000103374 |
| Aff2-201 | X | 29.97 | 2.41 | 3.76E-28 | ENSRNOT00000087990 |

| | | | | | |
|---|---|---|---|---|---|
| LOC103690322-201 | X | 21.45 | -5.9 | 7.88E-06 | ENSRNOT00000091365 |
| LOC100911047-201 | X | 10.49 | -2.24 | 0.02 | ENSRNOT00000075736 |
| Xlr3a-201 | X | 16.42 | -2.2 | 2.45E-04 | ENSRNOT00000074398 |
| LOC100911002-201 | X | 222.71 | -3.81 | 3.40E-96 | ENSRNOT00000071895 |
| Irak1-202 | X | 12.73 | 2.66 | 6.73E-03 | ENSRNOT00000102227 |
| LOC120099253-201 | X | 8.19 | -7.98 | 6.98E-03 | ENSRNOT00000088637 |

**Supplementary Table A3-3.** Differentially expressed downstream genes in PD 6.5 Erβ$^{KO}$ ovaries compared to PD 4.5 Erβ$^{KO}$ ovaries

| Name | Chrom | Max group mean | Fold change | FDR p-value | ENSEMBL |
|---|---|---|---|---|---|
| Nup43-201 | 1 | 7.69 | -546.85 | 4.77E-03 | ENSRNOT00000073528 |
| Utrn-205 | 1 | 9.12 | -5.74 | 3.07E-06 | ENSRNOT00000110616 |
| Map3k5-202 | 1 | 7.47 | -2.1 | 0.01 | ENSRNOT00000067070 |
| Slc2a12-201 | 1 | 16.71 | 2.01 | 3.08E-05 | ENSRNOT00000015566 |
| Fndc1-205 | 1 | 5.34 | -4.19 | 7.63E-03 | ENSRNOT00000099286 |
| Lnpep-202 | 1 | 9.05 | -3.35 | 4.69E-07 | ENSRNOT00000080325 |
| Lnpep-201 | 1 | 24.95 | 2.44 | 5.47E-09 | ENSRNOT00000017718 |
| ENSRNOT00000096430 | 1 | 5.03 | 2.15 | 0.05 | ENSRNOT00000096430 |
| Prkcg-202 | 1 | 6.85 | -2.35 | 0.05 | ENSRNOT00000104999 |
| Nlrp4f-201 | 1 | 6.24 | 3.19 | 0.03 | ENSRNOT00000078556 |
| Peg3-202 | 1 | 14.68 | 2.26 | 2.21E-12 | ENSRNOT00000106399 |
| Nlrp9-201 | 1 | 25.2 | 2.57 | 2.01E-07 | ENSRNOT00000029591 |
| Tnnt1-201 | 1 | 6.49 | -68.21 | 2.62E-03 | ENSRNOT00000034957 |
| Nlrp2-201 | 1 | 9.11 | 5.62 | 4.28E-07 | ENSRNOT00000073464 |
| Nectin2-203 | 1 | 25.14 | -3,508.81 | 1.24E-03 | ENSRNOT00000117594 |
| Zfp94-201 | 1 | 12.92 | -38.6 | 4.48E-07 | ENSRNOT00000026252 |
| Tex101-201 | 1 | 13.53 | -3.88 | 9.75E-03 | ENSRNOT00000027202 |
| Lypd10-204 | 1 | 23.85 | -2.05 | 0.04 | ENSRNOT00000072555 |
| Ltbp4-201 | 1 | 53.74 | -2.47 | 7.59E-09 | ENSRNOT00000028322 |
| Spint2-201 | 1 | 29.06 | -1,824.93 | 3.76E-03 | ENSRNOT00000028006 |
| Zfp260-202 | 1 | 6.97 | -136.07 | 2.47E-09 | ENSRNOT00000074301 |
| Zfp260-204 | 1 | 33.81 | 2.39 | 9.99E-15 | ENSRNOT00000113539 |
| Clip3-202 | 1 | 14.42 | -2.41 | 0.02 | ENSRNOT00000072819 |
| Lsr-204 | 1 | 11.47 | -2.63 | 0.02 | ENSRNOT00000116572 |
| Gramd1a-204 | 1 | 37.38 | -10.95 | 5.77E-24 | ENSRNOT00000116007 |
| Uevld-201 | 1 | 11.8 | 2.07 | 0.02 | ENSRNOT00000045719 |
| Nav2-201 | 1 | 5.44 | -4.33 | 3.55E-07 | ENSRNOT00000046529 |
| Mtmr10-203 | 1 | 6.83 | 2.49 | 5.10E-03 | ENSRNOT00000114828 |
| Tjp1-204 | 1 | 18.53 | 2.11 | 1.60E-08 | ENSRNOT00000112594 |
| Ttc23-204 | 1 | 6.75 | -3.37 | 0.05 | ENSRNOT00000078707 |
| St8sia2-201 | 1 | 9.33 | -2.4 | 3.43E-03 | ENSRNOT00000113519 |
| LOC103691165-203 | 1 | 26.07 | 2.38 | 1.19E-21 | ENSRNOT00000116764 |
| Il16-201 | 1 | 10.93 | -2.17 | 1.38E-04 | ENSRNOT00000016289 |
| Pcf11-201 | 1 | 5.03 | 294.31 | 2.75E-05 | ENSRNOT00000013552 |
| Tenm4-205 | 1 | 7.93 | 2.57 | 0.04 | ENSRNOT00000114854 |
| Tenm4-203 | 1 | 17.74 | 2.23 | 7.90E-12 | ENSRNOT00000111362 |
| Nars2-201 | 1 | 5.74 | 4.32 | 0.04 | ENSRNOT00000015784 |
| Fam168a-203 | 1 | 6.27 | 1,625.53 | 5.90E-03 | ENSRNOT00000097710 |
| Fchsd2-204 | 1 | 7.14 | -7.55 | 1.42E-03 | ENSRNOT00000118008 |
| Numa1-203 | 1 | 6.22 | 118.56 | 1.30E-17 | ENSRNOT00000094333 |
| Dennd2b-207 | 1 | 60.02 | -2.95 | 7.11E-20 | ENSRNOT00000114792 |
| Dennd2b-201 | 1 | 5.38 | 6.33 | 4.82E-05 | ENSRNOT00000064902 |
| Mical2-203 | 1 | 69.47 | 2.26 | 1.01E-24 | ENSRNOT00000021858 |
| Parva-201 | 1 | 24.96 | 2.38 | 0.03 | ENSRNOT00000021671 |
| Parva-203 | 1 | 8.01 | -91.34 | 1.28E-05 | ENSRNOT00000099634 |
| Tead1-203 | 1 | 6.48 | 3.62 | 0.02 | ENSRNOT00000096192 |
| Smg1-201 | 1 | 16.24 | 2.5 | 6.91E-05 | ENSRNOT00000089399 |

| | | | | | |
|---|---|---|---|---|---|
| Tnrc6a-203 | 1 | 9.27 | 2.2 | 2.40E-04 | ENSRNOT00000096111 |
| Aqp8-201 | 1 | 9.38 | 10.98 | 3.51E-04 | ENSRNOT00000019939 |
| Il4r-201 | 1 | 70.02 | 2.58 | 1.90E-16 | ENSRNOT00000020994 |
| Cln3-201 | 1 | 15.97 | -2.06 | 0.04 | ENSRNOT00000025898 |
| Rnf40-203 | 1 | 5.6 | -11.74 | 7.04E-03 | ENSRNOT00000098677 |
| Fgfr2-202 | 1 | 12.47 | -2.14 | 4.43E-03 | ENSRNOT00000022331 |
| Igf2-201 | 1 | 57.34 | -2.2 | 1.61E-30 | ENSRNOT00000080246 |
| Cdkn1c-202 | 1 | 11.63 | -2.64 | 0.02 | ENSRNOT00000078944 |
| Cttn-203 | 1 | 11.88 | 655.54 | 4.97E-10 | ENSRNOT00000054859 |
| Ighmbp2-203 | 1 | 8.91 | -2.55 | 4.95E-03 | ENSRNOT00000118525 |
| Tmem151a-201 | 1 | 15.12 | -2.64 | 7.54E-03 | ENSRNOT00000075365 |
| Cnih2-202 | 1 | 8.93 | -7.39 | 0.01 | ENSRNOT00000105446 |
| Sf1-204 | 1 | 18.26 | 2.22 | 4.66E-03 | ENSRNOT00000110004 |
| Stip1-202 | 1 | 8.72 | -1,026.62 | 1.12E-03 | ENSRNOT00000106195 |
| Hnrnpr-ps2-201 | 1 | 6.64 | 7.68 | 3.87E-03 | ENSRNOT00000071413 |
| Ahnak-204 | 1 | 7.36 | 2.43 | 3.32E-10 | ENSRNOT00000116008 |
| Ahnak-202 | 1 | 12.46 | 2.51 | 1.89E-18 | ENSRNOT00000088387 |
| Myrf-204 | 1 | 15.57 | -2.81 | 9.51E-09 | ENSRNOT00000116485 |
| Anxa1-204 | 1 | 18.84 | -3.15 | 0.02 | ENSRNOT00000096367 |
| Ptar1-201 | 1 | 13.54 | 3.1 | 1.89E-03 | ENSRNOT00000054775 |
| Kank1-202 | 1 | 28.12 | -2.8 | 3.71E-05 | ENSRNOT00000080689 |
| Vldlr-201 | 1 | 10.36 | 2.79 | 3.58E-05 | ENSRNOT00000035814 |
| Vldlr-202 | 1 | 12.88 | 2.67 | 1.30E-05 | ENSRNOT00000106818 |
| Kif20b-202 | 1 | 9.89 | 2.2 | 9.17E-05 | ENSRNOT00000085880 |
| Btaf1-205 | 1 | 5.18 | 4.62 | 5.35E-06 | ENSRNOT00000113933 |
| Lcor-202 | 1 | 8.11 | 2.74 | 4.05E-05 | ENSRNOT00000076842 |
| Hif1an-201 | 1 | 6.54 | 72.03 | 1.95E-03 | ENSRNOT00000019070 |
| Wbp1l-202 | 1 | 14.29 | -22.56 | 8.98E-11 | ENSRNOT00000100128 |
| Cyp17a1-201 | 1 | 120.52 | 4.3 | 1.98E-13 | ENSRNOT00000027160 |
| Sh3pxd2a-201 | 1 | 12.79 | 2.82 | 8.31E-04 | ENSRNOT00000027574 |
| Smc3-201 | 1 | 5.96 | 730.98 | 2.73E-03 | ENSRNOT00000019560 |
| Vwa2-201 | 1 | 16.03 | -2.4 | 1.35E-06 | ENSRNOT00000051834 |
| AABR07007032.1-202 | 1 | 14.89 | 2.15 | 0.03 | ENSRNOT00000078416 |
| Vcan-202 | 2 | 24.52 | -4.07 | 2.73E-07 | ENSRNOT00000063821 |
| Vcan-201 | 2 | 19.17 | 2.42 | 4.87E-12 | ENSRNOT00000045532 |
| Serinc5-202 | 2 | 14.3 | 2.17 | 4.35E-05 | ENSRNOT00000098981 |
| Zbed3-201 | 2 | 32.37 | -3.29 | 2.42E-04 | ENSRNOT00000040983 |
| Enc1-202 | 2 | 23.58 | -2.37 | 2.24E-03 | ENSRNOT00000106862 |
| Arhgef28-202 | 2 | 8.15 | 5.92 | 3.59E-14 | ENSRNOT00000115432 |
| Map1b-203 | 2 | 12.1 | -22.35 | 9.83E-07 | ENSRNOT00000113536 |
| Srek1-201 | 2 | 9.38 | 2.48 | 0.03 | ENSRNOT00000040533 |
| Pde4d-204 | 2 | 7.48 | 2.39 | 0.04 | ENSRNOT00000101684 |
| Mier3-201 | 2 | 6.78 | -346.27 | 1.12E-09 | ENSRNOT00000017571 |
| Il6st-202 | 2 | 32.37 | 2.47 | 5.71E-19 | ENSRNOT00000105192 |
| Mtrex-202 | 2 | 7.9 | 320.64 | 1.38E-07 | ENSRNOT00000095778 |
| Itga1-202 | 2 | 8.25 | 2.41 | 2.73E-03 | ENSRNOT00000106157 |
| Dab2-201 | 2 | 15.91 | 2.82 | 4.33E-05 | ENSRNOT00000050655 |
| Lifr-201 | 2 | 11.97 | 2.18 | 2.12E-03 | ENSRNOT00000016036 |
| Lmbrd2-201 | 2 | 9.02 | 2.1 | 5.76E-03 | ENSRNOT00000077646 |
| Mtmr12-203 | 2 | 5.59 | -256.49 | 5.36E-06 | ENSRNOT00000114002 |
| Pdzd2-203 | 2 | 6.64 | 2.92 | 0.03 | ENSRNOT00000105047 |

| | | | | | |
|---|---|---|---|---|---|
| Myo10-202 | 2 | 57.99 | 2.38 | 7.68E-12 | ENSRNOT00000065897 |
| Myo10-203 | 2 | 24.27 | -4.09 | 2.45E-04 | ENSRNOT00000102421 |
| Car3-202 | 2 | 21.58 | 2.31 | 2.33E-03 | ENSRNOT00000105296 |
| Lrrcc1-202 | 2 | 8.95 | 7.39 | 2.87E-06 | ENSRNOT00000097272 |
| Fabp4-201 | 2 | 46.8 | 2.21 | 0.02 | ENSRNOT00000014701 |
| Pag1-203 | 2 | 18.01 | 2.63 | 0.01 | ENSRNOT00000119112 |
| Zfp704-201 | 2 | 17.91 | 2.66 | 5.85E-10 | ENSRNOT00000080313 |
| Zc2hc1a-201 | 2 | 5.82 | 4.51 | 0.04 | ENSRNOT00000066966 |
| Zc2hc1a-206 | 2 | 11.11 | -6.66 | 2.90E-03 | ENSRNOT00000110271 |
| Pld1-201 | 2 | 16.15 | -2.94 | 0.02 | ENSRNOT00000039296 |
| Pld1-202 | 2 | 57.08 | 2.31 | 9.89E-27 | ENSRNOT00000039308 |
| Phc3-202 | 2 | 8.03 | 2.13 | 0.02 | ENSRNOT00000089477 |
| Phc3-201 | 2 | 8.39 | 2.37 | 0.01 | ENSRNOT00000066495 |
| Zfp639-203 | 2 | 5.49 | 4.83 | 0.02 | ENSRNOT00000100539 |
| Atp11b-204 | 2 | 20.64 | -2.18 | 4.97E-05 | ENSRNOT00000116170 |
| Trpc3-201 | 2 | 7.41 | 3.45 | 2.32E-04 | ENSRNOT00000046700 |
| Larp1b-202 | 2 | 18.94 | -3,469.11 | 1.17E-03 | ENSRNOT00000090861 |
| Naa15-203 | 2 | 15.26 | 2.22 | 2.06E-04 | ENSRNOT00000120176 |
| Naa15-202 | 2 | 16.06 | -6.69 | 1.31E-09 | ENSRNOT00000095909 |
| Spart-202 | 2 | 9.06 | -4.31 | 6.03E-03 | ENSRNOT00000116850 |
| Dclk1-204 | 2 | 5.22 | -27.41 | 2.94E-03 | ENSRNOT00000093284 |
| Mbnl1-205 | 2 | 7.95 | -2.47 | 0.05 | ENSRNOT00000106022 |
| Bche-201 | 2 | 6.93 | 2.21 | 0.05 | ENSRNOT00000013279 |
| Trim2-201 | 2 | 5.5 | -2,212.06 | 3.04E-03 | ENSRNOT00000067618 |
| Lrba-201 | 2 | 10.12 | -2.16 | 3.24E-03 | ENSRNOT00000023418 |
| Hdgf-202 | 2 | 32.36 | -8.18 | 9.22E-04 | ENSRNOT00000065211 |
| Ns5atp4-202 | 2 | 6.58 | -53.72 | 0.01 | ENSRNOT00000089801 |
| Them5-201 | 2 | 228.75 | -2.04 | 3.51E-15 | ENSRNOT00000066020 |
| Rorc-204 | 2 | 11.39 | -2.08 | 2.29E-03 | ENSRNOT00000118722 |
| Cgn-201 | 2 | 9.95 | -2.79 | 2.61E-04 | ENSRNOT00000028440 |
| Pogz-201 | 2 | 5.61 | 360.48 | 1.16E-07 | ENSRNOT00000082849 |
| Arnt-202 | 2 | 12.58 | 2.12 | 0.02 | ENSRNOT00000064442 |
| Notch2-201 | 2 | 22.78 | 2.48 | 1.27E-03 | ENSRNOT00000025718 |
| Hsd3b3-201 | 2 | 193.87 | 3.78 | 9.19E-40 | ENSRNOT00000056172 |
| Hsd3b1-201 | 2 | 615.01 | 3.62 | 4.06E-30 | ENSRNOT00000086835 |
| Ptgfrn-202 | 2 | 30.15 | -17.67 | 8.56E-51 | ENSRNOT00000096621 |
| Col11a1-201 | 2 | 72.46 | 2.12 | 1.48E-18 | ENSRNOT00000068413 |
| Arhgap29-202 | 2 | 18.41 | 4.8 | 7.59E-09 | ENSRNOT00000066486 |
| Lef1-201 | 2 | 8.29 | -3.2 | 4.49E-03 | ENSRNOT00000013694 |
| Aimp1-203 | 2 | 16.12 | -776.67 | 2.09E-03 | ENSRNOT00000112244 |
| Dnajb14-201 | 2 | 16.51 | 2.02 | 0.04 | ENSRNOT00000081989 |
| Prkacb-202 | 2 | 22.51 | -5,311.88 | 4.87E-04 | ENSRNOT00000109432 |
| Fubp1-203 | 2 | 17.37 | 2.13 | 5.12E-10 | ENSRNOT00000110491 |
| Pigk-204 | 2 | 9.85 | -123.32 | 7.10E-06 | ENSRNOT00000113062 |
| ENSRNOT00000108800 | 3 | 14,344.94 | 4.14 | 1.34E-27 | ENSRNOT00000108800 |
| ENSRNOT00000112407 | 3 | 7,081.59 | 3.95 | 6.84E-24 | ENSRNOT00000112407 |
| Spopl-201 | 3 | 15.43 | 2.33 | 0.04 | ENSRNOT00000033653 |
| Nsmf-202 | 3 | 11.92 | -482.99 | 7.40E-03 | ENSRNOT00000042100 |
| Gpsm1-202 | 3 | 20.25 | -2.18 | 2.68E-04 | ENSRNOT00000084567 |
| Sec16a-201 | 3 | 10.86 | 608.14 | 2.64E-18 | ENSRNOT00000064083 |
| Adamtsl2-201 | 3 | 18.13 | 2.26 | 1.22E-07 | ENSRNOT00000036995 |

| | | | | | |
|---|---|---|---|---|---|
| Col5a1-202 | 3 | 21.99 | 2.11 | 1.01E-10 | ENSRNOT00000086352 |
| Usp20-203 | 3 | 6.23 | -1,082.47 | 0.01 | ENSRNOT00000114450 |
| Hmcn2-201 | 3 | 7.81 | 3.81 | 1.98E-10 | ENSRNOT00000111278 |
| Ass1-202 | 3 | 23.53 | 2.4 | 5.39E-05 | ENSRNOT00000093365 |
| Prrc2b-202 | 3 | 25.75 | 2.05 | 1.55E-04 | ENSRNOT00000066368 |
| Eng-204 | 3 | 9.79 | -2,323.79 | 1.24E-04 | ENSRNOT00000114060 |
| Dab2ip-204 | 3 | 13.45 | 2.1 | 1.52E-05 | ENSRNOT00000094494 |
| Dab2ip-205 | 3 | 12.98 | 2.29 | 3.80E-06 | ENSRNOT00000103632 |
| Dab2ip-207 | 3 | 7.04 | -101.18 | 2.52E-17 | ENSRNOT00000108040 |
| Strbp-202 | 3 | 13.53 | 1,660.51 | 5.38E-03 | ENSRNOT00000087611 |
| Dennd1a-201 | 3 | 8.87 | -2.33 | 0.01 | ENSRNOT00000025443 |
| Scai-203 | 3 | 9.01 | 2.07 | 2.23E-03 | ENSRNOT00000104956 |
| Zeb2-201 | 3 | 9.53 | -2.66 | 0.01 | ENSRNOT00000006350 |
| Zeb2-204 | 3 | 13.01 | 2.1 | 1.76E-04 | ENSRNOT00000096459 |
| Zeb2-203 | 3 | 12.72 | 6.16 | 1.18E-03 | ENSRNOT00000085801 |
| Rif1-203 | 3 | 5.62 | 432.86 | 2.62E-16 | ENSRNOT00000097817 |
| Rif1-202 | 3 | 10.52 | 4.77 | 2.99E-13 | ENSRNOT00000094327 |
| Marchf7-202 | 3 | 11.41 | 4.02 | 0.01 | ENSRNOT00000085563 |
| Hnrnpa3-204 | 3 | 18.24 | -44.85 | 5.94E-08 | ENSRNOT00000114122 |
| Nfe2l2-203 | 3 | 33.48 | -2.64 | 5.85E-05 | ENSRNOT00000110389 |
| Itprid2-202 | 3 | 5.35 | -1,513.67 | 5.23E-03 | ENSRNOT00000105305 |
| Pde1a-203 | 3 | 12.61 | 126.01 | 1.14E-12 | ENSRNOT00000097071 |
| Zc3h15-202 | 3 | 7.28 | -839.89 | 0.02 | ENSRNOT00000094114 |
| Fam171b-201 | 3 | 5.62 | 3.3 | 9.24E-04 | ENSRNOT00000006504 |
| Arhgap1-203 | 3 | 8.92 | -2.94 | 3.24E-03 | ENSRNOT00000098688 |
| Ambra1-201 | 3 | 6.9 | 4.37 | 0.02 | ENSRNOT00000045644 |
| Mapk8ip1-203 | 3 | 8.92 | -2.87 | 0.04 | ENSRNOT00000117436 |
| Accsl-201 | 3 | 15.45 | 2.54 | 5.61E-04 | ENSRNOT00000068585 |
| Fmn1-205 | 3 | 6.81 | 2.93 | 8.13E-03 | ENSRNOT00000114830 |
| Fmn1-204 | 3 | 8.92 | 3.01 | 2.27E-05 | ENSRNOT00000098989 |
| Aqr-202 | 3 | 8.27 | -11.3 | 9.12E-06 | ENSRNOT00000105477 |
| Nusap1-201 | 3 | 8.81 | 331.03 | 6.48E-08 | ENSRNOT00000052312 |
| Rtf1-201 | 3 | 30.57 | -2.03 | 1.65E-05 | ENSRNOT00000056392 |
| Snap23-201 | 3 | 5.8 | 39 | 4.20E-04 | ENSRNOT00000074392 |
| Stard9-202 | 3 | 34.34 | 3.64 | 7.67E-55 | ENSRNOT00000079109 |
| Stard9-201 | 3 | 5.49 | -4,877.36 | 5.82E-04 | ENSRNOT00000048141 |
| Ttbk2-202 | 3 | 11.24 | 2.35 | 1.83E-04 | ENSRNOT00000090680 |
| Tp53bp1-203 | 3 | 9.84 | -14.81 | 2.85E-09 | ENSRNOT00000107143 |
| Tp53bp1-202 | 3 | 8.25 | -126.52 | 3.93E-16 | ENSRNOT00000106843 |
| Tp53bp1-201 | 3 | 30.36 | 2.19 | 1.57E-03 | ENSRNOT00000019025 |
| Anapc1-202 | 3 | 13.12 | 2.29 | 1.02E-05 | ENSRNOT00000099712 |
| Dnaaf9-203 | 3 | 8.08 | -2.47 | 6.82E-03 | ENSRNOT00000112794 |
| Dnaaf9-201 | 3 | 13.12 | 2.47 | 1.43E-06 | ENSRNOT00000028842 |
| ENSRNOT00000028881 | 3 | 123.52 | -2.53 | 1.44E-07 | ENSRNOT00000028881 |
| Lrrn4-201 | 3 | 25.38 | -2.2 | 2.22E-08 | ENSRNOT00000040802 |
| Nrsn2-201 | 3 | 14.38 | -2.55 | 0.03 | ENSRNOT00000032881 |
| Phf20-202 | 3 | 6.23 | 3.41 | 4.70E-03 | ENSRNOT00000095858 |
| Soga1-202 | 3 | 23.15 | 2.07 | 3.91E-09 | ENSRNOT00000102663 |
| Rpn2-201 | 3 | 39.72 | -2.51 | 1.05E-14 | ENSRNOT00000068135 |
| Nnat-202 | 3 | 198.72 | -2.01 | 4.51E-08 | ENSRNOT00000072502 |
| Zhx3-201 | 3 | 7.11 | 2.04 | 0.03 | ENSRNOT00000032588 |

| | | | | | |
|---|---|---|---|---|---|
| Chd6-205 | 3 | 13.81 | -10.94 | 1.92E-06 | ENSRNOT00000114568 |
| Chd6-204 | 3 | 5.18 | -2.22 | 0.03 | ENSRNOT00000107966 |
| Sdc4-201 | 3 | 67.09 | -2.01 | 3.44E-12 | ENSRNOT00000019386 |
| B4galt5-201 | 3 | 5.81 | -622.66 | 3.64E-03 | ENSRNOT00000064849 |
| Gnas-207 | 3 | 213.26 | -2.11 | 3.54E-10 | ENSRNOT00000115047 |
| LOC103690425-201 | 3 | 9.45 | 2.74 | 5.54E-03 | ENSRNOT00000109993 |
| Sycp2-201 | 3 | 6.05 | -4.57 | 6.75E-06 | ENSRNOT00000078047 |
| Arfgap1-202 | 3 | 9.22 | -1,651.40 | 6.80E-03 | ENSRNOT00000055038 |
| Kmt2c-203 | 4 | 5.63 | 20.73 | 5.40E-26 | ENSRNOT00000103015 |
| Prkag2-203 | 4 | 9.92 | -2.21 | 0.02 | ENSRNOT00000106083 |
| Fastk-204 | 4 | 24.37 | -575.4 | 1.52E-06 | ENSRNOT00000118434 |
| Slc4a2-202 | 4 | 12.76 | -9.92 | 4.57E-05 | ENSRNOT00000108170 |
| LOC103692025-202 | 4 | 12.34 | 6.24 | 6.55E-04 | ENSRNOT00000104480 |
| Pmpcb-203 | 4 | 13.48 | -2.42 | 0.05 | ENSRNOT00000101407 |
| RGD1565355-204 | 4 | 12.15 | 4.04 | 6.45E-03 | ENSRNOT00000075962 |
| RGD1565355-201 | 4 | 5.63 | 460.98 | 6.85E-03 | ENSRNOT00000008319 |
| Cacna2d1-201 | 4 | 18.05 | 2.25 | 8.33E-08 | ENSRNOT00000042914 |
| Elapor2-202 | 4 | 8.41 | 2.79 | 5.00E-06 | ENSRNOT00000100645 |
| Cdk14-202 | 4 | 16.35 | -3.12 | 1.75E-08 | ENSRNOT00000094279 |
| Akap9-205 | 4 | 7.87 | 2.56 | 5.45E-05 | ENSRNOT00000111753 |
| Cdk6-201 | 4 | 13.78 | 2.12 | 0.05 | ENSRNOT00000012597 |
| Peg10-202 | 4 | 6.89 | 4.41 | 5.34E-03 | ENSRNOT00000104707 |
| Nrf1-205 | 4 | 5.69 | 2.12 | 0.05 | ENSRNOT00000105926 |
| Podxl-201 | 4 | 21.02 | -177.45 | 2.00E-43 | ENSRNOT00000016991 |
| Ubn2-203 | 4 | 8.28 | 3.24 | 7.42E-03 | ENSRNOT00000096540 |
| Zfp786-201 | 4 | 5.5 | 890.59 | 0.01 | ENSRNOT00000038550 |
| Hnrnpa2b1-202 | 4 | 29.01 | 2.19 | 8.71E-03 | ENSRNOT00000015325 |
| Adcyap1r1-205 | 4 | 6.48 | -2.85 | 0.03 | ENSRNOT00000103561 |
| Krcc1-202 | 4 | 22.3 | 2.43 | 0.02 | ENSRNOT00000094782 |
| St3gal5-203 | 4 | 25.87 | -2.74 | 0.02 | ENSRNOT00000115948 |
| Vamp5-201 | 4 | 33.96 | 2.12 | 4.33E-04 | ENSRNOT00000016984 |
| Capg-201 | 4 | 12.24 | -8.99 | 9.06E-04 | ENSRNOT00000018562 |
| Tet3-201 | 4 | 12.07 | 3.26 | 6.31E-10 | ENSRNOT00000015296 |
| Fbln2-203 | 4 | 104.86 | -2.41 | 5.79E-39 | ENSRNOT00000115833 |
| Nr2c2-201 | 4 | 24.29 | 2.38 | 1.26E-04 | ENSRNOT00000014353 |
| Itpr1-202 | 4 | 12.12 | 2.33 | 1.59E-03 | ENSRNOT00000043646 |
| Itpr1-204 | 4 | 5.98 | -4.45 | 1.14E-07 | ENSRNOT00000082723 |
| Arpc4-203 | 4 | 9.21 | -4.57 | 0.04 | ENSRNOT00000105231 |
| Ttll3-202 | 4 | 6.74 | -3.51 | 0.02 | ENSRNOT00000102169 |
| Fancd2-203 | 4 | 7.47 | -85.31 | 9.23E-13 | ENSRNOT00000094549 |
| Slc6a1-201 | 4 | 14.52 | -2.73 | 2.65E-03 | ENSRNOT00000009705 |
| H1f8-201 | 4 | 56.24 | 2.26 | 1.47E-06 | ENSRNOT00000042673 |
| Erc1-202 | 4 | 10.15 | 2.08 | 5.89E-03 | ENSRNOT00000091473 |
| Wnk1-204 | 4 | 16.66 | 2.36 | 2.67E-10 | ENSRNOT00000080788 |
| Wnk1-203 | 4 | 7.02 | -3.29 | 3.86E-07 | ENSRNOT00000079513 |
| Atp6v1e1-202 | 4 | 33.15 | -21.95 | 4.81E-09 | ENSRNOT00000098217 |
| Zfp384-203 | 4 | 5.92 | -1,132.33 | 8.76E-03 | ENSRNOT00000036654 |
| Atf7ip-201 | 4 | 10.82 | 2.1 | 7.02E-04 | ENSRNOT00000011803 |
| Plbd1-201 | 4 | 15.04 | 2.45 | 9.88E-03 | ENSRNOT00000011930 |
| Dera-203 | 4 | 5.09 | 288.72 | 0.02 | ENSRNOT00000116905 |
| Ppfibp1-201 | 4 | 12.2 | 3.04 | 5.14E-06 | ENSRNOT00000044147 |

| | | | | | |
|---|---|---|---|---|---|
| Dennd5b-203 | 4 | 12.97 | 2.09 | 7.70E-05 | ENSRNOT00000105569 |
| Pi15-201 | 5 | 10 | 3.11 | 0.04 | ENSRNOT00000023899 |
| Prex2-204 | 5 | 8.11 | 2.13 | 4.90E-04 | ENSRNOT00000108865 |
| Rb1cc1-203 | 5 | 9.55 | 3.41 | 4.23E-06 | ENSRNOT00000118113 |
| ENSRNOT00000113061 | 5 | 295.09 | 2.28 | 6.23E-05 | ENSRNOT00000113061 |
| Slc26a7-202 | 5 | 6.95 | 3.28 | 0.04 | ENSRNOT00000093186 |
| Nfx1-201 | 5 | 14.33 | 9.2 | 3.34E-09 | ENSRNOT00000060691 |
| Ncbp1-202 | 5 | 13.1 | -4.91 | 0.01 | ENSRNOT00000094126 |
| Akap2-205 | 5 | 6.7 | 2.31 | 8.16E-03 | ENSRNOT00000098662 |
| Susd1-201 | 5 | 8.62 | 2.21 | 0.03 | ENSRNOT00000085109 |
| Pappa-202 | 5 | 5.53 | 5.31 | 2.31E-11 | ENSRNOT00000076467 |
| Ptprd-204 | 5 | 13.97 | 2.57 | 1.51E-09 | ENSRNOT00000113220 |
| Ptprd-203 | 5 | 7.17 | 4.31 | 1.44E-07 | ENSRNOT00000110800 |
| Ptprd-205 | 5 | 7.12 | -2.19 | 2.04E-03 | ENSRNOT00000115119 |
| Zfp353-201 | 5 | 26.09 | 2.96 | 1.10E-06 | ENSRNOT00000072732 |
| Elavl2-201 | 5 | 12.9 | 5.64 | 2.77E-12 | ENSRNOT00000009034 |
| Elavl2-202 | 5 | 12.5 | 2.68 | 0.03 | ENSRNOT00000092743 |
| Mysm1-201 | 5 | 13.61 | 2.78 | 2.34E-05 | ENSRNOT00000039554 |
| Fyb2-204 | 5 | 6.97 | 2.32 | 0.04 | ENSRNOT00000095104 |
| Lrp8-205 | 5 | 33.9 | 3.22 | 1.14E-22 | ENSRNOT00000112745 |
| Lrp8-201 | 5 | 10.54 | 2.98 | 0.01 | ENSRNOT00000017575 |
| Scp2-202 | 5 | 5.13 | -185.96 | 4.72E-05 | ENSRNOT00000015466 |
| Osbpl9-204 | 5 | 8.2 | -69.98 | 3.32E-04 | ENSRNOT00000105115 |
| Nasp-201 | 5 | 53.05 | 2.17 | 7.37E-11 | ENSRNOT00000022170 |
| Ptprf-203 | 5 | 9.38 | -2.95 | 4.22E-03 | ENSRNOT00000102268 |
| Cited4-201 | 5 | 26.96 | 2.56 | 8.45E-04 | ENSRNOT00000072427 |
| Macf1-202 | 5 | 5.04 | 7,047.30 | 4.14E-06 | ENSRNOT00000081482 |
| LOC100294508-201 | 5 | 18.34 | 2.6 | 7.61E-11 | ENSRNOT00000016493 |
| LOC100294508-202 | 5 | 9.57 | -524.02 | 3.68E-08 | ENSRNOT00000100704 |
| RGD1561149-203 | 5 | 16.18 | -3.22 | 1.92E-11 | ENSRNOT00000108263 |
| Fabp3-201 | 5 | 206.46 | 2 | 2.21E-12 | ENSRNOT00000017325 |
| Pla2g2a-201 | 5 | 48.1 | -2.31 | 3.77E-05 | ENSRNOT00000022827 |
| Oog3-201 | 5 | 68.58 | 2.06 | 3.01E-08 | ENSRNOT00000055741 |
| Oog1-201 | 5 | 9.45 | 2.91 | 0.05 | ENSRNOT00000046115 |
| RGD1306186-202 | 5 | 72.61 | 2.14 | 1.49E-10 | ENSRNOT00000091144 |
| Ube4b-202 | 5 | 15.09 | -4,021.84 | 9.37E-04 | ENSRNOT00000105982 |
| Pik3cd-202 | 5 | 38.15 | 2.02 | 1.39E-19 | ENSRNOT00000095643 |
| Rere-203 | 5 | 35.77 | -2.12 | 1.46E-12 | ENSRNOT00000103362 |
| Nadk-202 | 5 | 18.88 | 2.03 | 7.37E-05 | ENSRNOT00000094609 |
| Lhcgr-201 | 6 | 15.58 | 4.02 | 2.84E-10 | ENSRNOT00000022481 |
| Srbd1-202 | 6 | 8.91 | -144.67 | 7.22E-04 | ENSRNOT00000096560 |
| Eml4-203 | 6 | 31.21 | -2.06 | 1.29E-11 | ENSRNOT00000111306 |
| Eml4-204 | 6 | 7.84 | 2.9 | 1.03E-03 | ENSRNOT00000116346 |
| Pkdcc-202 | 6 | 36.41 | 2.07 | 2.91E-07 | ENSRNOT00000104247 |
| Map4k3-203 | 6 | 8.04 | 3.33 | 0.03 | ENSRNOT00000084602 |
| Birc6-202 | 6 | 12.41 | 3.25 | 9.28E-22 | ENSRNOT00000083128 |
| Dpysl5-201 | 6 | 19.77 | -2.12 | 0.04 | ENSRNOT00000012273 |
| Atad2b-202 | 6 | 7.66 | 2.12 | 1.48E-03 | ENSRNOT00000084560 |
| Greb1-201 | 6 | 19.64 | 6.75 | 1.38E-56 | ENSRNOT00000032417 |
| Itgb1bp1-203 | 6 | 9.54 | -8.86 | 0.02 | ENSRNOT00000116410 |
| Mboat2-201 | 6 | 19.2 | 2.24 | 2.76E-05 | ENSRNOT00000082657 |

| | | | | | |
|---|---|---|---|---|---|
| Mboat2-202 | 6 | 8.79 | -4.07 | 9.67E-04 | ENSRNOT00000093903 |
| Lamb1-204 | 6 | 95.82 | -3.33 | 3.20E-32 | ENSRNOT00000114806 |
| Gpr22-201 | 6 | 6.07 | 2.02 | 0.05 | ENSRNOT00000011659 |
| Prkar2b-201 | 6 | 213.42 | 2.04 | 3.77E-20 | ENSRNOT00000012415 |
| Ralgapa1-205 | 6 | 19.53 | 2.66 | 1.11E-20 | ENSRNOT00000107390 |
| Ralgapa1-202 | 6 | 5.4 | 3.59 | 4.75E-03 | ENSRNOT00000065230 |
| Nin-203 | 6 | 8.57 | 2.57 | 6.40E-05 | ENSRNOT00000103711 |
| Rhoj-201 | 6 | 12.88 | 2.4 | 2.34E-03 | ENSRNOT00000031979 |
| Esr2-203 | 6 | 10.76 | 9.39 | 9.75E-07 | ENSRNOT00000042682 |
| Rdh11-203 | 6 | 6.54 | 572.1 | 4.69E-03 | ENSRNOT00000105155 |
| Smoc1-203 | 6 | 75.93 | 2.04 | 5.07E-13 | ENSRNOT00000114143 |
| Ylpm1-201 | 6 | 14.13 | -2.01 | 4.64E-05 | ENSRNOT00000006492 |
| Ston2-201 | 6 | 14.65 | 2.23 | 1.08E-03 | ENSRNOT00000005881 |
| Spata7-208 | 6 | 7.14 | -681.89 | 0.02 | ENSRNOT00000118680 |
| Ttc7b-201 | 6 | 15.98 | -2.55 | 1.17E-03 | ENSRNOT00000037902 |
| Ppp4r3a-204 | 6 | 8.3 | -277.56 | 2.04E-12 | ENSRNOT00000114004 |
| Begain-204 | 6 | 6.1 | -240.34 | 0.03 | ENSRNOT00000117298 |
| Eif5-202 | 6 | 8.44 | 3.66 | 0.03 | ENSRNOT00000102717 |
| Klc1-204 | 6 | 20.56 | 2.09 | 0.03 | ENSRNOT00000093106 |
| Klc1-201 | 6 | 13.81 | -2.6 | 5.96E-05 | ENSRNOT00000015935 |
| ENSRNOT00000109415 | 6 | 17.83 | -12.77 | 0.01 | ENSRNOT00000109415 |
| Jag2-202 | 6 | 6.5 | -2.29 | 0.02 | ENSRNOT00000100725 |
| Dync2i1-201 | 6 | 9.37 | -236.92 | 1.89E-11 | ENSRNOT00000006144 |
| Dync2i1-205 | 6 | 10.61 | 3.1 | 1.33E-04 | ENSRNOT00000109958 |
| Rab5b-203 | 7 | 26.19 | -11.68 | 1.12E-17 | ENSRNOT00000120020 |
| Nfic-203 | 7 | 11.89 | 2.51 | 0.02 | ENSRNOT00000097025 |
| Mex3d-201 | 7 | 8.83 | 1,263.67 | 8.32E-03 | ENSRNOT00000040170 |
| Rps15-202 | 7 | 23.07 | 6.21 | 0.02 | ENSRNOT00000113730 |
| Tmem259-202 | 7 | 37.03 | -2.02 | 5.59E-04 | ENSRNOT00000080292 |
| Tmem259-201 | 7 | 40.12 | 2.08 | 3.50E-05 | ENSRNOT00000017245 |
| Palm-203 | 7 | 11.31 | -6.67 | 0.01 | ENSRNOT00000114530 |
| Chst11-201 | 7 | 29.13 | 2.07 | 4.49E-06 | ENSRNOT00000012017 |
| Igf1-204 | 7 | 140.76 | 2.32 | 2.36E-35 | ENSRNOT00000118307 |
| Tmpo-209 | 7 | 7.61 | 4.19 | 0.01 | ENSRNOT00000113551 |
| Eea1-206 | 7 | 11.8 | 2.92 | 9.20E-06 | ENSRNOT00000117665 |
| Cep290-203 | 7 | 6.04 | 2.58 | 0.02 | ENSRNOT00000101678 |
| Tmtc2-201 | 7 | 14.7 | 2.45 | 3.77E-06 | ENSRNOT00000006086 |
| Csrp2-202 | 7 | 42.73 | 2.28 | 8.99E-07 | ENSRNOT00000080598 |
| Csrp2-203 | 7 | 6.37 | 2.12 | 3.84E-04 | ENSRNOT00000097951 |
| Osbpl8-201 | 7 | 24.05 | 2.12 | 2.25E-09 | ENSRNOT00000091910 |
| Osbpl8-205 | 7 | 7.04 | -201.78 | 1.06E-19 | ENSRNOT00000112719 |
| Frs2-202 | 7 | 6.27 | 2.22 | 0.02 | ENSRNOT00000108286 |
| Mdm2-201 | 7 | 16.85 | -2.83 | 7.08E-03 | ENSRNOT00000066767 |
| Slc16a7-201 | 7 | 9.28 | 2.38 | 0.02 | ENSRNOT00000058036 |
| Vps13b-203 | 7 | 8.5 | -11.96 | 3.03E-24 | ENSRNOT00000114232 |
| Vps13b-202 | 7 | 16.34 | 2.76 | 1.60E-15 | ENSRNOT00000105415 |
| Ubr5-201 | 7 | 25.59 | -3.53 | 5.65E-24 | ENSRNOT00000009115 |
| Angpt1-201 | 7 | 5.35 | 4.51 | 2.41E-04 | ENSRNOT00000007979 |
| Mtss1-204 | 7 | 5.82 | -3.49 | 0.03 | ENSRNOT00000104332 |
| Mtss1-201 | 7 | 14.26 | -11.31 | 2.69E-09 | ENSRNOT00000011984 |
| Ago2-201 | 7 | 19.22 | 2.74 | 1.50E-06 | ENSRNOT00000011898 |

| | | | | | |
|---|---|---|---|---|---|
| Ptk2-208 | 7 | 10.06 | -4.29 | 1.54E-09 | ENSRNOT00000113790 |
| Gga1-201 | 7 | 32.96 | -2.99 | 9.58E-09 | ENSRNOT00000063876 |
| Ddx17-203 | 7 | 174 | -2.72 | 6.08E-07 | ENSRNOT00000076164 |
| Tnrc6b-204 | 7 | 6.7 | 3.25 | 6.55E-06 | ENSRNOT00000106327 |
| Cyb5r3-202 | 7 | 19.74 | -3.08 | 2.45E-04 | ENSRNOT00000102247 |
| Parvb-202 | 7 | 23.11 | -2.21 | 3.53E-08 | ENSRNOT00000082805 |
| Zbed4-201 | 7 | 8.31 | 3.15 | 0.01 | ENSRNOT00000006050 |
| Alg10-201 | 7 | 27.94 | 2.07 | 7.30E-06 | ENSRNOT00000019672 |
| Rapgef3-202 | 7 | 8.8 | -2.28 | 0.03 | ENSRNOT00000083182 |
| Cacnb3-201 | 7 | 28.1 | -2.05 | 5.32E-05 | ENSRNOT00000081206 |
| Prkag1-201 | 7 | 8.24 | 3.91 | 7.93E-08 | ENSRNOT00000077502 |
| Prph-201 | 7 | 11.64 | -2.47 | 0.04 | ENSRNOT00000089211 |
| Nr4a1-201 | 7 | 15.75 | 3.86 | 2.04E-06 | ENSRNOT00000010171 |
| Krt7-201 | 7 | 24.04 | -2.25 | 3.92E-04 | ENSRNOT00000010660 |
| Sesn3-203 | 8 | 31.76 | 2.13 | 4.48E-07 | ENSRNOT00000110335 |
| Olfm2-203 | 8 | 9.18 | -5.52 | 0.02 | ENSRNOT00000117797 |
| Ldlr-201 | 8 | 21.94 | 2.07 | 4.10E-10 | ENSRNOT00000013496 |
| Ldlr-202 | 8 | 7.58 | -16.98 | 2.53E-06 | ENSRNOT00000108195 |
| Arhgap32-203 | 8 | 6.93 | 2.22 | 0.02 | ENSRNOT00000100925 |
| Ei24-202 | 8 | 6.28 | -709.88 | 2.87E-03 | ENSRNOT00000110156 |
| RGD1311744-202 | 8 | 8.68 | 5.3 | 9.37E-04 | ENSRNOT00000110068 |
| Gramd1b-204 | 8 | 10.22 | 3.08 | 3.94E-06 | ENSRNOT00000099621 |
| Clmp-202 | 8 | 8.96 | -2.37 | 0.02 | ENSRNOT00000108273 |
| Cbl-202 | 8 | 9.41 | 2.91 | 1.38E-03 | ENSRNOT00000101607 |
| Bco2-203 | 8 | 14.21 | 2.23 | 8.35E-04 | ENSRNOT00000059192 |
| Sik2-203 | 8 | 8.41 | -3.79 | 2.80E-05 | ENSRNOT00000119371 |
| Sik2-202 | 8 | 7.94 | 1,059.07 | 3.61E-13 | ENSRNOT00000091732 |
| Btg4-202 | 8 | 10.09 | 3.84 | 0.03 | ENSRNOT00000088320 |
| Cul5-201 | 8 | 24.78 | 2.01 | 4.42E-05 | ENSRNOT00000010956 |
| Cyp19a1-201 | 8 | 66.93 | 3.01 | 1.01E-14 | ENSRNOT00000000212 |
| Peak1-201 | 8 | 15.27 | 2.22 | 7.68E-09 | ENSRNOT00000068658 |
| Lingo1-202 | 8 | 5.69 | -4.6 | 0.04 | ENSRNOT00000094295 |
| Stra6-201 | 8 | 10.85 | -2.2 | 0.02 | ENSRNOT00000011074 |
| Arih1-203 | 8 | 6.32 | 2.29 | 0.03 | ENSRNOT00000098010 |
| Arih1-202 | 8 | 5.88 | 472.21 | 6.94E-03 | ENSRNOT00000084758 |
| Kif23-201 | 8 | 10.49 | 6.28 | 7.04E-03 | ENSRNOT00000037028 |
| Cilp-201 | 8 | 21.11 | -2.78 | 4.47E-15 | ENSRNOT00000044887 |
| Clpx-204 | 8 | 10.78 | -17.23 | 3.60E-04 | ENSRNOT00000110101 |
| Plekho2-202 | 8 | 5.05 | -168.4 | 1.18E-04 | ENSRNOT00000096404 |
| Tln2-201 | 8 | 19.97 | -2.31 | 4.51E-13 | ENSRNOT00000012026 |
| Tln2-202 | 8 | 5.09 | 99.17 | 1.21E-17 | ENSRNOT00000024885 |
| Ccnb2-203 | 8 | 31.12 | -10.34 | 9.24E-06 | ENSRNOT00000118940 |
| Tcf12-201 | 8 | 45.59 | 2.26 | 4.23E-10 | ENSRNOT00000081185 |
| Khdc3-201 | 8 | 16.93 | 3.57 | 0.05 | ENSRNOT00000079458 |
| Eef1a1-202 | 8 | 15.55 | -2.11 | 0.05 | ENSRNOT00000112753 |
| Phip-201 | 8 | 16.25 | 2.11 | 3.97E-13 | ENSRNOT00000011864 |
| Tent5a-201 | 8 | 25.4 | 2.23 | 9.22E-08 | ENSRNOT00000056937 |
| Cep162-202 | 8 | 5.57 | -145.87 | 5.17E-11 | ENSRNOT00000083934 |
| Syncrip-202 | 8 | 45.9 | 2.56 | 5.90E-05 | ENSRNOT00000074515 |
| Spsb4-201 | 8 | 19.75 | 2.17 | 2.27E-04 | ENSRNOT00000017363 |
| Amotl2-202 | 8 | 7.07 | -1,463.14 | 8.61E-03 | ENSRNOT00000090221 |

| | | | | | |
|---|---|---|---|---|---|
| Tf-202 | 8 | 55.81 | -3.54 | 4.52E-27 | ENSRNOT00000045628 |
| Tmem108-203 | 8 | 12.29 | -3.4 | 5.71E-08 | ENSRNOT00000104880 |
| Atp2c1-202 | 8 | 12.32 | 66.46 | 3.34E-13 | ENSRNOT00000045087 |
| Rbm15b-202 | 8 | 6.36 | 2.26 | 1.52E-03 | ENSRNOT00000097719 |
| Ifrd2-203 | 8 | 9.69 | -974.01 | 0.01 | ENSRNOT00000112128 |
| Sema3f-202 | 8 | 17.73 | -2.02 | 5.65E-03 | ENSRNOT00000085915 |
| Smarcc1-201 | 8 | 10.74 | 2.08 | 5.93E-03 | ENSRNOT00000064140 |
| Setd2-201 | 8 | 11.56 | -628.64 | 3.07E-20 | ENSRNOT00000028409 |
| Clasp2-205 | 8 | 5.67 | 2,104.65 | 2.90E-03 | ENSRNOT00000107377 |
| Clasp2-201 | 8 | 6.09 | -68.52 | 1.17E-04 | ENSRNOT00000012545 |
| Vill-201 | 8 | 14.78 | -2.2 | 1.12E-04 | ENSRNOT00000015888 |
| Plcd1-202 | 8 | 20.5 | -2.65 | 5.72E-05 | ENSRNOT00000110954 |
| Safb2-202 | 9 | 5.17 | 3.9 | 9.86E-07 | ENSRNOT00000097383 |
| Trerf1-203 | 9 | 15.19 | 2.07 | 1.18E-04 | ENSRNOT00000100821 |
| Ptk7-204 | 9 | 8.75 | 125.18 | 1.74E-15 | ENSRNOT00000118183 |
| Ptk7-201 | 9 | 6.02 | -5.62 | 9.57E-03 | ENSRNOT00000061432 |
| Cd2ap-202 | 9 | 21.29 | 2.01 | 1.64E-05 | ENSRNOT00000107057 |
| Dst-205 | 9 | 19 | 2.75 | 3.10E-15 | ENSRNOT00000101194 |
| Dst-203 | 9 | 11.87 | -11.23 | 3.58E-63 | ENSRNOT00000095842 |
| Dst-208 | 9 | 14.85 | 6.54 | 1.53E-70 | ENSRNOT00000112664 |
| Plekhb2-202 | 9 | 23.33 | -2.39 | 1.59E-06 | ENSRNOT00000097171 |
| Lman2l-204 | 9 | 11.97 | -3.25 | 5.03E-03 | ENSRNOT00000119365 |
| Inpp4a-203 | 9 | 14.71 | -3.39 | 4.59E-08 | ENSRNOT00000096431 |
| Eif5b-201 | 9 | 5.85 | -156.12 | 2.03E-05 | ENSRNOT00000035785 |
| Lonrf2-201 | 9 | 7.57 | 3.19 | 2.58E-03 | ENSRNOT00000033964 |
| Spats2l-203 | 9 | 10.17 | -1,363.54 | 4.60E-04 | ENSRNOT00000100381 |
| Spats2l-202 | 9 | 27.96 | -3.3 | 2.08E-06 | ENSRNOT00000081497 |
| Orc2-202 | 9 | 6.58 | -905.07 | 2.39E-03 | ENSRNOT00000099033 |
| Nop58-202 | 9 | 13.37 | 2.67 | 0.01 | ENSRNOT00000091283 |
| Bmpr2-202 | 9 | 10.3 | 2.96 | 2.06E-09 | ENSRNOT00000100349 |
| Bmpr2-201 | 9 | 36.26 | 2.16 | 9.94E-16 | ENSRNOT00000035238 |
| Ino80d-201 | 9 | 8.86 | 2.9 | 4.38E-05 | ENSRNOT00000035879 |
| Ndufs1-202 | 9 | 15.99 | -2.21 | 9.66E-03 | ENSRNOT00000098172 |
| Pikfyve-201 | 9 | 5.93 | 2.16 | 1.76E-05 | ENSRNOT00000020447 |
| Atic-202 | 9 | 17.35 | 12.17 | 1.82E-03 | ENSRNOT00000098789 |
| Cnot9-203 | 9 | 8.85 | -8.28 | 5.54E-03 | ENSRNOT00000107071 |
| Col4a4-203 | 9 | 22.5 | -2.49 | 3.72E-06 | ENSRNOT00000111693 |
| Pid1-202 | 9 | 44.73 | 2.49 | 3.27E-11 | ENSRNOT00000084446 |
| Cab39-203 | 9 | 17.04 | -4.4 | 4.64E-10 | ENSRNOT00000102022 |
| Col6a3-203 | 9 | 57.18 | 3.79 | 1.04E-39 | ENSRNOT00000087595 |
| Col6a3-201 | 9 | 5.38 | 18.08 | 1.19E-31 | ENSRNOT00000026719 |
| Ndufa10-203 | 9 | 8.09 | 3.8 | 0.02 | ENSRNOT00000110509 |
| Kif1a-203 | 9 | 7.93 | -40.33 | 2.40E-03 | ENSRNOT00000120091 |
| Kif1a-202 | 9 | 6.9 | -3.69 | 8.12E-04 | ENSRNOT00000106149 |
| Farp2-202 | 9 | 10.83 | 2.9 | 8.84E-10 | ENSRNOT00000110498 |
| ENSRNOT00000110318 | 9 | 16.74 | 3.74 | 2.31E-11 | ENSRNOT00000110318 |
| Ppip5k2-202 | 9 | 5.4 | 2.3 | 0.04 | ENSRNOT00000076669 |
| Abcc1-203 | 10 | 10.83 | -346.91 | 7.99E-13 | ENSRNOT00000115220 |
| Rpl39l-201 | 10 | 13.04 | -774.95 | 0.01 | ENSRNOT00000051564 |
| Glyr1-204 | 10 | 5.43 | 7.84 | 7.83E-03 | ENSRNOT00000106608 |
| Crebbp-202 | 10 | 5.37 | -6.86 | 0.03 | ENSRNOT00000117260 |

| | | | | | |
|---|---|---|---|---|---|
| Srrm2-201 | 10 | 38.7 | 18,234.56 | 1.65E-07 | ENSRNOT00000091287 |
| Mslnl-201 | 10 | 15.2 | -2.02 | 9.32E-04 | ENSRNOT00000060286 |
| Msln-201 | 10 | 17.83 | -3.11 | 1.96E-06 | ENSRNOT00000026395 |
| Rab11fip3-201 | 10 | 12.59 | -3.24 | 1.80E-03 | ENSRNOT00000027439 |
| Hba-a1-201 | 10 | 1,979.97 | -2.27 | 8.50E-10 | ENSRNOT00000051483 |
| Hba-a1-202 | 10 | 1,783.91 | -2.08 | 1.28E-08 | ENSRNOT00000052292 |
| Hba-a1-206 | 10 | 1,511.76 | -2.37 | 1.59E-17 | ENSRNOT00000119443 |
| Fbxw11-203 | 10 | 10.7 | -37.07 | 2.71E-06 | ENSRNOT00000098062 |
| Havcr1-201 | 10 | 15.91 | 4.81 | 1.48E-03 | ENSRNOT00000009573 |
| Larp1-202 | 10 | 24.23 | 2.03 | 7.38E-07 | ENSRNOT00000113107 |
| Rasd1-201 | 10 | 39.15 | 2.22 | 5.64E-07 | ENSRNOT00000004475 |
| Tom1l2-207 | 10 | 8.06 | -3.84 | 0.03 | ENSRNOT00000111837 |
| Rnf112-201 | 10 | 15.19 | -2.08 | 7.84E-03 | ENSRNOT00000003228 |
| Ndel1-202 | 10 | 21.38 | -2.33 | 3.29E-04 | ENSRNOT00000065505 |
| Chd3-204 | 10 | 5.8 | 2.27 | 1.43E-03 | ENSRNOT00000114886 |
| Kdm6b-202 | 10 | 16.42 | 2.12 | 2.45E-07 | ENSRNOT00000116399 |
| Alox15-201 | 10 | 63.32 | -2.38 | 1.18E-10 | ENSRNOT00000026038 |
| P2rx1-202 | 10 | 39.02 | -2.54 | 8.03E-14 | ENSRNOT00000088245 |
| Rap1gap2-205 | 10 | 11.84 | 3.03 | 1.29E-03 | ENSRNOT00000103095 |
| Hic1-201 | 10 | 7.99 | 2.48 | 0.01 | ENSRNOT00000004143 |
| Nufip2-202 | 10 | 36.42 | 2.02 | 1.87E-11 | ENSRNOT00000101390 |
| ENSRNOT00000115958 | 10 | 22.48 | 2.02 | 3.73E-04 | ENSRNOT00000115958 |
| Ap2b1-203 | 10 | 24.8 | -5.97 | 3.37E-04 | ENSRNOT00000090446 |
| Ypel2-202 | 10 | 14.42 | -14.01 | 6.19E-28 | ENSRNOT00000114367 |
| Mtmr4-202 | 10 | 6.49 | 3.05 | 0.01 | ENSRNOT00000099688 |
| Mtmr4-203 | 10 | 5.35 | 2.49 | 6.23E-03 | ENSRNOT00000109434 |
| Mmd-201 | 10 | 28.94 | -14.24 | 4.09E-15 | ENSRNOT00000003308 |
| Utp18-201 | 10 | 17.24 | 2.18 | 0.02 | ENSRNOT00000003582 |
| Spag9-204 | 10 | 22.46 | 2.31 | 5.04E-08 | ENSRNOT00000094632 |
| Spag9-208 | 10 | 7.39 | 12.9 | 5.64E-22 | ENSRNOT00000117102 |
| Luc7l3-202 | 10 | 18.59 | 2.39 | 7.43E-03 | ENSRNOT00000101307 |
| Col1a1-203 | 10 | 163.55 | -53,077.70 | 2.87E-06 | ENSRNOT00000115096 |
| Col1a1-202 | 10 | 98.61 | -12.18 | 1.42E-25 | ENSRNOT00000114204 |
| Ngfr-202 | 10 | 12.77 | -411.69 | 5.85E-10 | ENSRNOT00000107122 |
| Srcin1-201 | 10 | 13.35 | 2.81 | 6.77E-08 | ENSRNOT00000016070 |
| Med1-202 | 10 | 7 | 3.57 | 1.22E-03 | ENSRNOT00000101134 |
| Cdk12-202 | 10 | 7.84 | 2.01 | 0.02 | ENSRNOT00000082668 |
| Cdk12-205 | 10 | 11.48 | -3.7 | 3.38E-03 | ENSRNOT00000038939 |
| Med24-202 | 10 | 40.26 | -2.01 | 2.62E-16 | ENSRNOT00000100598 |
| Krt16-203 | 10 | 321.2 | -2.7 | 7.78E-24 | ENSRNOT00000019133 |
| Klhl11-201 | 10 | 61.53 | 2.11 | 1.65E-12 | ENSRNOT00000022695 |
| Hsd17b1-201 | 10 | 26.14 | 3.78 | 2.64E-07 | ENSRNOT00000026878 |
| Mlx-202 | 10 | 6.56 | 713.4 | 0.02 | ENSRNOT00000099833 |
| Mapt-209 | 10 | 9 | -4.12 | 0.02 | ENSRNOT00000115998 |
| Ace-202 | 10 | 10.88 | -5.88 | 1.15E-04 | ENSRNOT00000092961 |
| Bptf-204 | 10 | 5.91 | 2.28 | 1.68E-04 | ENSRNOT00000104873 |
| Abca8a-202 | 10 | 11.86 | 2.05 | 2.26E-04 | ENSRNOT00000065947 |
| Cog1-204 | 10 | 9.03 | -2.57 | 3.04E-04 | ENSRNOT00000113704 |
| St6galnac2-203 | 10 | 12.31 | -2.09 | 0.02 | ENSRNOT00000093828 |
| Tnrc6c-202 | 10 | 12.05 | 2.73 | 3.67E-07 | ENSRNOT00000106642 |
| Tepsin-202 | 10 | 5.45 | -8.26 | 3.44E-03 | ENSRNOT00000098173 |

| | | | | | |
|---|---|---|---|---|---|
| Nploc4-201 | 10 | 30.56 | -12.34 | 3.85E-21 | ENSRNOT00000054973 |
| Nploc4-202 | 10 | 43.41 | 2.17 | 1.10E-11 | ENSRNOT00000080188 |
| Foxk2-203 | 10 | 17.69 | -3.81 | 3.62E-05 | ENSRNOT00000099287 |
| LOC102550944-202 | 10 | 6.1 | 962.23 | 0.01 | ENSRNOT00000107856 |
| Robo1-203 | 11 | 9.21 | 2.47 | 5.09E-05 | ENSRNOT00000104951 |
| Nrip1-201 | 11 | 6.78 | 2.39 | 0.02 | ENSRNOT00000002152 |
| App-202 | 11 | 37.24 | -2.58 | 1.28E-04 | ENSRNOT00000048854 |
| Tiam1-202 | 11 | 6.7 | -2.67 | 4.01E-03 | ENSRNOT00000109463 |
| Son-201 | 11 | 34.05 | 6.68 | 3.58E-06 | ENSRNOT00000002769 |
| Crybg3-202 | 11 | 11.19 | 2.2 | 2.80E-12 | ENSRNOT00000074653 |
| Crybg3-201 | 11 | 11.36 | -2.68 | 6.36E-08 | ENSRNOT00000002292 |
| Dzip3-201 | 11 | 16.55 | 3.01 | 1.72E-08 | ENSRNOT00000002678 |
| Usf3-201 | 11 | 8.15 | 2.01 | 5.26E-06 | ENSRNOT00000046208 |
| Naa50-203 | 11 | 5.56 | -123 | 7.47E-13 | ENSRNOT00000098332 |
| Golgb1-203 | 11 | 8.01 | 3.45 | 4.58E-25 | ENSRNOT00000106429 |
| Golgb1-202 | 11 | 9.73 | 4.78 | 6.95E-19 | ENSRNOT00000083152 |
| Acap2-201 | 11 | 7.36 | 2.64 | 0.01 | ENSRNOT00000029549 |
| Atp13a3-201 | 11 | 5.1 | 2.35 | 5.44E-04 | ENSRNOT00000098527 |
| Lpp-203 | 11 | 10.17 | 3.63 | 1.12E-09 | ENSRNOT00000107426 |
| Lpp-201 | 11 | 6.61 | 4.07 | 0.02 | ENSRNOT00000044279 |
| Clcn2-204 | 11 | 6.85 | -814.63 | 1.83E-03 | ENSRNOT00000073435 |
| Dgcr2-203 | 11 | 24.67 | -6.84 | 8.57E-06 | ENSRNOT00000107429 |
| Smpd4-202 | 11 | 7.13 | 7.15 | 3.31E-04 | ENSRNOT00000102059 |
| Dnm1l-201 | 11 | 12.36 | 2.94 | 8.33E-04 | ENSRNOT00000002477 |
| Insr-203 | 12 | 15.67 | 2.32 | 2.26E-10 | ENSRNOT00000113217 |
| Arhgef18-203 | 12 | 5.19 | -370.4 | 4.31E-08 | ENSRNOT00000110862 |
| Stxbp2-203 | 12 | 6.54 | -3.27 | 0.04 | ENSRNOT00000115749 |
| Map2k7-201 | 12 | 5.2 | 82.17 | 8.29E-04 | ENSRNOT00000061821 |
| Kpna7-201 | 12 | 8.17 | 4.75 | 0.01 | ENSRNOT00000075986 |
| Fscn1-203 | 12 | 35.54 | -2.19 | 5.96E-03 | ENSRNOT00000102873 |
| Stag3-201 | 12 | 26.94 | -2.37 | 4.98E-14 | ENSRNOT00000042006 |
| Col26a1-201 | 12 | 25.35 | -2.83 | 2.03E-10 | ENSRNOT00000080068 |
| Cux1-204 | 12 | 7.52 | 4.07 | 1.84E-09 | ENSRNOT00000112152 |
| Upk3b-201 | 12 | 34.46 | -2.5 | 2.53E-05 | ENSRNOT00000037639 |
| Ncor2-205 | 12 | 32.93 | -2.08 | 4.54E-07 | ENSRNOT00000104620 |
| Ppp1cc-203 | 12 | 5.62 | -405.04 | 0.01 | ENSRNOT00000083296 |
| Git2-201 | 12 | 8.6 | -3.88 | 4.21E-03 | ENSRNOT00000045946 |
| Ube3b-204 | 12 | 8.63 | -2,514.75 | 2.33E-03 | ENSRNOT00000116293 |
| Acacb-204 | 12 | 5.15 | 2.13 | 0.02 | ENSRNOT00000080557 |
| RGD1306556-202 | 12 | 12.15 | -8.15 | 8.19E-08 | ENSRNOT00000095507 |
| Ttc28-201 | 12 | 6.31 | 2.3 | 2.04E-04 | ENSRNOT00000046920 |
| Ep400-204 | 12 | 8.46 | 11.48 | 3.25E-13 | ENSRNOT00000118850 |
| Ptpn4-202 | 13 | 12.96 | 2.75 | 2.25E-07 | ENSRNOT00000095537 |
| Ptpn4-201 | 13 | 19.74 | 2.31 | 1.24E-13 | ENSRNOT00000003568 |
| Steap3-202 | 13 | 11.78 | -380.77 | 3.56E-08 | ENSRNOT00000101486 |
| Actr3-201 | 13 | 19.82 | 205.11 | 1.41E-16 | ENSRNOT00000004520 |
| Rab3gap1-201 | 13 | 9.72 | 10.74 | 2.54E-07 | ENSRNOT00000005289 |
| Nucks1-204 | 13 | 76.26 | 2.34 | 2.06E-14 | ENSRNOT00000112895 |
| Mdm4-201 | 13 | 60.17 | 2.16 | 5.56E-14 | ENSRNOT00000012984 |
| Zbed6-202 | 13 | 34.35 | 2.4 | 8.09E-13 | ENSRNOT00000115014 |
| Nr5a2-202 | 13 | 69.93 | 2.05 | 2.90E-15 | ENSRNOT00000097279 |

| | | | | | |
|---|---|---|---|---|---|
| Nek7-204 | 13 | 6.49 | -195.16 | 6.90E-06 | ENSRNOT00000098011 |
| Dennd1b-202 | 13 | 7.26 | 2.42 | 5.30E-04 | ENSRNOT00000043907 |
| Ro60-201 | 13 | 11.35 | 2.16 | 6.61E-04 | ENSRNOT00000004701 |
| Edem3-201 | 13 | 13.63 | 2.7 | 7.55E-08 | ENSRNOT00000017858 |
| Arpc5-204 | 13 | 14.18 | -1,362.30 | 6.05E-03 | ENSRNOT00000107757 |
| Smg7-202 | 13 | 6.08 | -2,014.27 | 1.41E-04 | ENSRNOT00000092269 |
| Smg7-204 | 13 | 9.07 | 2.93 | 1.07E-07 | ENSRNOT00000092397 |
| Mrps14-201 | 13 | 7.25 | 522.99 | 0.03 | ENSRNOT00000003460 |
| Rc3h1-201 | 13 | 18.28 | 2.41 | 3.47E-08 | ENSRNOT00000003689 |
| Mpzl1-202 | 13 | 29.77 | -3.74 | 0.04 | ENSRNOT00000093821 |
| Ddr2-201 | 13 | 17.32 | 2.54 | 4.74E-12 | ENSRNOT00000003901 |
| Fcrla-201 | 13 | 22.86 | -2.21 | 0.02 | ENSRNOT00000004183 |
| Cdc42bpa-206 | 13 | 23.58 | 3,425.58 | 1.18E-18 | ENSRNOT00000116713 |
| Ephx1-203 | 13 | 17.87 | 1,826.48 | 4.13E-03 | ENSRNOT00000085279 |
| Tlr5-202 | 13 | 52.59 | 2.36 | 3.26E-06 | ENSRNOT00000094732 |
| Mark1-201 | 13 | 28.33 | 2.5 | 3.27E-13 | ENSRNOT00000003198 |
| Lrrc8d-205 | 14 | 7.03 | 1,494.84 | 5.28E-03 | ENSRNOT00000109682 |
| Lrrc8d-203 | 14 | 21.33 | -2.45 | 6.33E-08 | ENSRNOT00000105738 |
| Ptpn13-201 | 14 | 13.6 | 2.45 | 2.26E-08 | ENSRNOT00000061162 |
| Bmp2k-202 | 14 | 15.55 | 2.11 | 4.71E-05 | ENSRNOT00000101160 |
| Septin11-205 | 14 | 12.7 | 5.19 | 1.16E-03 | ENSRNOT00000111632 |
| Septin11-204 | 14 | 11.87 | 3.44 | 9.66E-03 | ENSRNOT00000103937 |
| Rchy1-202 | 14 | 5.41 | -74.01 | 1.80E-03 | ENSRNOT00000104798 |
| Ociad1-202 | 14 | 15.44 | -681.83 | 0.02 | ENSRNOT00000094276 |
| Fbxl5-204 | 14 | 21.01 | -3.24 | 8.86E-05 | ENSRNOT00000103550 |
| Tbc1d14-201 | 14 | 6.18 | 2.76 | 0.03 | ENSRNOT00000058414 |
| Eif4enif1-205 | 14 | 35.62 | -2.43 | 1.03E-06 | ENSRNOT00000102028 |
| Pik3ip1-201 | 14 | 11.13 | -2.09 | 5.63E-03 | ENSRNOT00000057738 |
| Morc2-201 | 14 | 12.8 | -5.98 | 6.84E-24 | ENSRNOT00000026569 |
| Thoc5-202 | 14 | 24.77 | -2.15 | 1.89E-05 | ENSRNOT00000096419 |
| Sertad2-209 | 14 | 5.83 | 2.41 | 1.07E-03 | ENSRNOT00000115608 |
| Xpo1-202 | 14 | 27.63 | -2.08 | 2.27E-05 | ENSRNOT00000101915 |
| Usp34-202 | 14 | 23.43 | 2.76 | 2.55E-15 | ENSRNOT00000094041 |
| Efemp1-203 | 14 | 6.2 | -209.71 | 3.02E-04 | ENSRNOT00000101622 |
| Ccdc88a-201 | 14 | 5.99 | 3.17 | 1.81E-04 | ENSRNOT00000067591 |
| Sptbn1-202 | 14 | 11.73 | 2.26 | 1.12E-07 | ENSRNOT00000084595 |
| Dlg5-204 | 15 | 8.1 | 2.12 | 2.75E-03 | ENSRNOT00000112381 |
| Nr1d2-203 | 15 | 18.92 | -4.6 | 0.02 | ENSRNOT00000106171 |
| Slc4a7-206 | 15 | 5.09 | 3.17 | 7.52E-07 | ENSRNOT00000118797 |
| Ptprg-203 | 15 | 7.95 | -307.88 | 1.64E-06 | ENSRNOT00000099333 |
| Ktn1-204 | 15 | 7.13 | -6.27 | 9.98E-03 | ENSRNOT00000106317 |
| Tep1-202 | 15 | 11.89 | -13.27 | 3.21E-07 | ENSRNOT00000088981 |
| Tep1-201 | 15 | 18.31 | 2.31 | 5.41E-15 | ENSRNOT00000079506 |
| Arhgef40-204 | 15 | 11.58 | 3.49 | 3.97E-04 | ENSRNOT00000115526 |
| Supt16h-201 | 15 | 29.65 | 11.95 | 5.80E-07 | ENSRNOT00000016288 |
| Acin1-201 | 15 | 17.2 | 2.38 | 7.98E-05 | ENSRNOT00000018982 |
| Slc22a17-202 | 15 | 22.79 | -2,381.63 | 2.63E-03 | ENSRNOT00000102200 |
| Msra-203 | 15 | 5.6 | -47.25 | 2.69E-03 | ENSRNOT00000095387 |
| Hmbox1-202 | 15 | 12.86 | 2.08 | 1.14E-10 | ENSRNOT00000109493 |
| Trim35-202 | 15 | 10.82 | -2.6 | 1.96E-04 | ENSRNOT00000113248 |
| Dpysl2-202 | 15 | 69.67 | -2.32 | 0.04 | ENSRNOT00000103229 |

| | | | | | |
|---|---|---|---|---|---|
| Ppp3cc-202 | 15 | 7.82 | 166.31 | 2.61E-04 | ENSRNOT00000082505 |
| Fndc3a-205 | 15 | 21.04 | 3.87 | 8.88E-13 | ENSRNOT00000111875 |
| Mycbp2-205 | 15 | 6.97 | -3.48 | 4.64E-10 | ENSRNOT00000109278 |
| Mycbp2-206 | 15 | 11.82 | 3.05 | 3.12E-05 | ENSRNOT00000112777 |
| Ednrb-201 | 15 | 38.59 | 2.66 | 2.22E-11 | ENSRNOT00000014747 |
| Dzip1-201 | 15 | 22.78 | -2.77 | 0.04 | ENSRNOT00000038596 |
| Dock9-206 | 15 | 5.64 | 5.14 | 8.70E-05 | ENSRNOT00000111187 |
| Itgbl1-201 | 15 | 15.5 | 2.66 | 6.94E-05 | ENSRNOT00000006264 |
| Tasor-201 | 16 | 15.58 | 2.81 | 1.18E-07 | ENSRNOT00000074069 |
| Dcp1a-202 | 16 | 6.84 | -2,175.40 | 2.85E-03 | ENSRNOT00000111266 |
| Nisch-203 | 16 | 19.54 | -2.27 | 0.02 | ENSRNOT00000113930 |
| Mettl6-204 | 16 | 21.83 | -3.14 | 2.84E-03 | ENSRNOT00000103715 |
| Ankrd28-204 | 16 | 10.38 | 2.99 | 1.38E-05 | ENSRNOT00000112368 |
| Mapk8-203 | 16 | 8.07 | -2.04 | 0.01 | ENSRNOT00000099742 |
| Mast3-203 | 16 | 6.52 | -2.31 | 9.92E-03 | ENSRNOT00000115023 |
| Psd3-204 | 16 | 12.48 | 2.33 | 8.90E-15 | ENSRNOT00000093876 |
| Psd3-207 | 16 | 7.89 | 2.33 | 4.06E-04 | ENSRNOT00000113025 |
| Psd3-201 | 16 | 5.04 | -2.96 | 2.68E-03 | ENSRNOT00000041994 |
| Hpgd-201 | 16 | 15.12 | 5.31 | 2.22E-06 | ENSRNOT00000014229 |
| Gpm6a-201 | 16 | 17.46 | -1,040.72 | 1.56E-03 | ENSRNOT00000014312 |
| Fat1-203 | 16 | 19.18 | -3.16 | 9.27E-16 | ENSRNOT00000095529 |
| Fat1-201 | 16 | 18.9 | 15.48 | 8.35E-08 | ENSRNOT00000067486 |
| Pcm1-201 | 16 | 8.19 | 3.78 | 4.23E-03 | ENSRNOT00000014202 |
| Pcm1-202 | 16 | 9.88 | 2.18 | 1.08E-03 | ENSRNOT00000085435 |
| Tex15-202 | 16 | 5.58 | -236.96 | 1.31E-09 | ENSRNOT00000107080 |
| AABR07026291.1-205 | 16 | 10.83 | 2.31 | 2.23E-03 | ENSRNOT00000117895 |
| Arhgef7-202 | 16 | 12.25 | -2.77 | 1.33E-04 | ENSRNOT00000043449 |
| Naa35-204 | 17 | 17.55 | 2.06 | 0.02 | ENSRNOT00000108214 |
| Fam193b-205 | 17 | 5.98 | 2.03 | 5.78E-03 | ENSRNOT00000114459 |
| Nsd1-205 | 17 | 21.12 | 3.14 | 7.28E-18 | ENSRNOT00000118233 |
| Nsd1-203 | 17 | 50.52 | 2.1 | 3.09E-21 | ENSRNOT00000107527 |
| Faf2-203 | 17 | 10.09 | -2.84 | 1.89E-03 | ENSRNOT00000095037 |
| Simc1-201 | 17 | 8.64 | -254.27 | 3.18E-05 | ENSRNOT00000022980 |
| Ogn-201 | 17 | 322.92 | 2.08 | 2.21E-18 | ENSRNOT00000020532 |
| Ogn-202 | 17 | 126.89 | 2.31 | 1.02E-07 | ENSRNOT00000093561 |
| Omd-201 | 17 | 35.68 | 3.43 | 4.82E-14 | ENSRNOT00000020648 |
| Phf2-204 | 17 | 8.32 | 2.93 | 0.03 | ENSRNOT00000110404 |
| Prpf4b-201 | 17 | 45.52 | -2.42 | 7.43E-09 | ENSRNOT00000022902 |
| Serpinb9-204 | 17 | 11.02 | 331.68 | 6.13E-05 | ENSRNOT00000115382 |
| Exoc2-205 | 17 | 5.8 | -1,426.00 | 8.76E-03 | ENSRNOT00000092101 |
| Carmil1-201 | 17 | 10.66 | -3,016.88 | 1.58E-03 | ENSRNOT00000059510 |
| Sfrp4-202 | 17 | 8.49 | 2.79 | 0.02 | ENSRNOT00000079368 |
| Inhba-201 | 17 | 66.87 | 5.34 | 4.37E-38 | ENSRNOT00000019272 |
| Mkx-201 | 17 | 6.51 | 2.29 | 0.05 | ENSRNOT00000025623 |
| Gtpbp4-202 | 17 | 11.44 | 2.23 | 0.02 | ENSRNOT00000095348 |
| Tasor2-203 | 17 | 5.53 | 2.4 | 1.76E-03 | ENSRNOT00000118354 |
| Usp6nl-202 | 17 | 7.46 | -18.26 | 1.19E-08 | ENSRNOT00000103406 |
| Prpf18-203 | 17 | 13.02 | -3.21 | 1.02E-04 | ENSRNOT00000110607 |
| Nmt2-205 | 17 | 25.25 | -2.39 | 9.86E-04 | ENSRNOT00000119939 |
| Greb1l-201 | 18 | 13.25 | 2.47 | 8.90E-16 | ENSRNOT00000033468 |
| Esco1-204 | 18 | 5.76 | -10.39 | 0.01 | ENSRNOT00000118344 |

| | | | | | |
|---|---|---|---|---|---|
| Map3k2-201 | 18 | 10.97 | 2.65 | 5.42E-03 | ENSRNOT00000060996 |
| Epb41l4a-202 | 18 | 7.1 | -3.47 | 0.04 | ENSRNOT00000086729 |
| Apc-203 | 18 | 11.07 | -2.39 | 9.74E-06 | ENSRNOT00000102176 |
| Brd8-207 | 18 | 19.83 | -2.38 | 0.03 | ENSRNOT00000118939 |
| Kdm3b-203 | 18 | 15.74 | -3.66 | 2.04E-12 | ENSRNOT00000111068 |
| Eif4ebp3-202 | 18 | 21.68 | -3.57 | 0.03 | ENSRNOT00000032316 |
| Pcdha5-201 | 18 | 7.7 | 2.23 | 2.00E-04 | ENSRNOT00000027340 |
| Hdac3-201 | 18 | 11.4 | 6.18 | 0.02 | ENSRNOT00000060417 |
| Dpysl3-202 | 18 | 38.32 | -2.81 | 7.23E-05 | ENSRNOT00000087876 |
| Dpysl3-201 | 18 | 12.11 | 2.99 | 6.58E-05 | ENSRNOT00000025880 |
| LOC103690064-204 | 18 | 25.67 | 2.13 | 7.98E-06 | ENSRNOT00000076060 |
| Mcc-203 | 18 | 17.47 | 2.45 | 3.34E-06 | ENSRNOT00000085714 |
| Sncaip-202 | 18 | 12.96 | 2.14 | 6.78E-03 | ENSRNOT00000082760 |
| Pdgfrb-202 | 18 | 24.18 | -5.95 | 5.89E-05 | ENSRNOT00000078764 |
| Csnk1a1-203 | 18 | 15.82 | 3.41 | 1.26E-03 | ENSRNOT00000103047 |
| Txnl1-204 | 18 | 20.53 | -292.54 | 1.11E-05 | ENSRNOT00000100860 |
| Ccbe1-201 | 18 | 11.67 | 2.06 | 9.21E-06 | ENSRNOT00000116155 |
| Ctif-203 | 18 | 8.7 | 2.05 | 3.96E-03 | ENSRNOT00000098958 |
| Zbtb7c-202 | 18 | 6 | -14.12 | 0.01 | ENSRNOT00000101836 |
| Epg5-202 | 18 | 11.03 | -79.77 | 5.72E-51 | ENSRNOT00000108039 |
| Cnot1-203 | 19 | 22.82 | 3.01 | 9.37E-12 | ENSRNOT00000118032 |
| Hmgxb4-204 | 19 | 9.93 | -6.06 | 7.08E-05 | ENSRNOT00000110845 |
| Chd9-203 | 19 | 11.28 | 2.38 | 1.71E-12 | ENSRNOT00000078786 |
| Zfp423-203 | 19 | 6.39 | 2.83 | 0.02 | ENSRNOT00000101895 |
| Zfp423-201 | 19 | 11.18 | -3.45 | 5.44E-08 | ENSRNOT00000020028 |
| Gpt2-203 | 19 | 7.59 | -4.64 | 0.03 | ENSRNOT00000099582 |
| LOC685989-201 | 19 | 83.46 | -6.88 | 1.64E-24 | ENSRNOT00000049865 |
| Adgrl1-206 | 19 | 7.59 | 2.86 | 3.06E-07 | ENSRNOT00000103294 |
| Matcap1-202 | 19 | 7.59 | -2.73 | 0.05 | ENSRNOT00000107741 |
| Cyb5b-201 | 19 | 12.58 | 2.69 | 5.51E-04 | ENSRNOT00000015006 |
| Atxn1l-201 | 19 | 8.99 | -3.17 | 1.89E-05 | ENSRNOT00000059225 |
| Phlpp2-201 | 19 | 17.7 | 2.44 | 7.02E-13 | ENSRNOT00000021744 |
| Chst4-201 | 19 | 5.08 | -2.94 | 0.02 | ENSRNOT00000022749 |
| Cntnap4-201 | 19 | 5.72 | -2.27 | 0.03 | ENSRNOT00000066057 |
| Cdh13-202 | 19 | 12.09 | -2.15 | 0.02 | ENSRNOT00000077053 |
| Ankrd11-202 | 19 | 18.08 | 2.18 | 5.64E-22 | ENSRNOT00000103866 |
| Nrp1-201 | 19 | 10.22 | 108.26 | 3.51E-19 | ENSRNOT00000014492 |
| Nrp1-204 | 19 | 6.11 | -6.14 | 0.01 | ENSRNOT00000114861 |
| Gabbr1-206 | 20 | 16.64 | -218.05 | 1.19E-20 | ENSRNOT00000090936 |
| Zfp57-201 | 20 | 98.07 | 2.29 | 2.60E-20 | ENSRNOT00000011704 |
| Prrc2a-202 | 20 | 58.63 | 3.49 | 1.58E-38 | ENSRNOT00000092322 |
| Prrc2a-203 | 20 | 44.02 | -2.11 | 3.01E-13 | ENSRNOT00000111770 |
| Cfb-201 | 20 | 11.4 | -2.23 | 0.01 | ENSRNOT00000000477 |
| Col11a2-201 | 20 | 7.05 | -2.03 | 0.01 | ENSRNOT00000045533 |
| Nudt3-202 | 20 | 8.14 | -131 | 1.25E-04 | ENSRNOT00000096637 |
| Bltp3a-204 | 20 | 11.2 | 2.11 | 3.84E-04 | ENSRNOT00000104971 |
| Tead3-203 | 20 | 18.82 | -2.16 | 0.04 | ENSRNOT00000107495 |
| Kctd20-203 | 20 | 10.83 | -85.88 | 3.55E-17 | ENSRNOT00000105827 |
| Stk38-202 | 20 | 9.6 | -4.13 | 2.64E-04 | ENSRNOT00000092400 |
| Abcg1-202 | 20 | 8.3 | -511.24 | 4.94E-06 | ENSRNOT00000078031 |
| Pcbp3-203 | 20 | 22.38 | -2.18 | 2.65E-04 | ENSRNOT00000103785 |

| | | | | | |
|---|---|---|---|---|---|
| Col6a2-202 | 20 | 61.49 | 2.48 | 3.29E-26 | ENSRNOT00000097191 |
| Dip2a-201 | 20 | 8.28 | 2.14 | 3.04E-03 | ENSRNOT00000066017 |
| Jmjd1c-202 | 20 | 9.46 | 2.7 | 2.72E-06 | ENSRNOT00000099911 |
| Reep3-202 | 20 | 13.53 | 2.19 | 0.04 | ENSRNOT00000109024 |
| Tet1-201 | 20 | 5.2 | 2.74 | 5.59E-04 | ENSRNOT00000000302 |
| Dse-201 | 20 | 11.53 | 2.35 | 4.98E-04 | ENSRNOT00000001093 |
| Lims1-201 | 20 | 10.7 | 5.97 | 2.65E-03 | ENSRNOT00000064288 |
| Ranbp2-203 | 20 | 18.61 | 2.18 | 6.39E-07 | ENSRNOT00000114220 |
| Oit3-202 | 20 | 18.58 | -3.17 | 0.05 | ENSRNOT00000112351 |
| Slc9a7-201 | X | 7.85 | 2.53 | 0.04 | ENSRNOT00000005623 |
| Usp9x-201 | X | 29.3 | 3.93 | 4.13E-16 | ENSRNOT00000048980 |
| Tbc1d25-202 | X | 6.76 | -6.71 | 6.96E-03 | ENSRNOT00000099656 |
| Clcn5-202 | X | 11.46 | 2.53 | 4.23E-04 | ENSRNOT00000079054 |
| Ezhip-201 | X | 21.79 | 2.21 | 1.25E-03 | ENSRNOT00000060213 |
| Spin2b-201 | X | 49.37 | 2.14 | 1.42E-06 | ENSRNOT00000042670 |
| Spin2b-203 | X | 485.88 | 2.61 | 1.66E-50 | ENSRNOT00000114926 |
| Spin2b-202 | X | 87.92 | 2.3 | 1.79E-21 | ENSRNOT00000113757 |
| Huwe1-203 | X | 29.47 | 3.03 | 2.02E-19 | ENSRNOT00000098599 |
| Cdkl5-201 | X | 7.24 | 2.78 | 7.92E-03 | ENSRNOT00000005061 |
| Sh3kbp1-205 | X | 17.57 | -2.02 | 4.22E-03 | ENSRNOT00000096669 |
| Mageb6b1-201 | X | 5.86 | 523.94 | 0.03 | ENSRNOT00000043817 |
| Ar-201 | X | 9.59 | 3.53 | 4.02E-07 | ENSRNOT00000009129 |
| Ogt-203 | X | 100.89 | -94.73 | 6.74E-33 | ENSRNOT00000100805 |
| Cxhxorf49-202 | X | 14.62 | -2.11 | 0.04 | ENSRNOT00000097163 |
| Chm-203 | X | 6.54 | 107.16 | 1.90E-16 | ENSRNOT00000102082 |
| Gprasp1-205 | X | 7.05 | 2.87 | 1.49E-03 | ENSRNOT00000109064 |
| Fam199x-201 | X | 13.71 | 2.04 | 1.82E-03 | ENSRNOT00000071973 |
| Pak3-201 | X | 17.84 | 2.71 | 4.28E-05 | ENSRNOT00000006459 |
| Il13ra2-201 | X | 48.73 | 2.12 | 1.80E-08 | ENSRNOT00000043338 |
| Il13ra1-ps1-201 | X | 70.21 | 2.37 | 1.07E-16 | ENSRNOT00000017746 |
| Il13ra1-ps1-202 | X | 16.58 | -51.75 | 8.58E-15 | ENSRNOT00000112175 |
| Smarca1-202 | X | 6.98 | 4.32 | 2.66E-03 | ENSRNOT00000114801 |
| Bgn-203 | X | 18.3 | -2.41 | 8.61E-04 | ENSRNOT00000104560 |
| Mecp2-201 | X | 11.65 | 2.76 | 0.05 | ENSRNOT00000085723 |

**Supplementary Table A3-4.** Differentially expressed downstream genes in PD 8.5 Erβ$^{KO}$ ovaries compared to PD 6.5 Erβ$^{KO}$ ovaries.

| Transcript Variant | Chrom | Max group mean | Fold change | FDR p-value | ENSEMBL |
|---|---|---|---|---|---|
| Map3k5-202 | 1 | 6.55 | 2.13 | 0.03 | ENSRNOT00000067070 |
| Sgk1-202 | 1 | 35.23 | 2.01 | 1.70E-04 | ENSRNOT00000040736 |
| Lnpep-203 | 1 | 5.22 | -4.63 | 0.02 | ENSRNOT00000088586 |
| Nlrp4b-201 | 1 | 10.71 | 3.05 | 2.46E-03 | ENSRNOT00000035435 |
| Nlrp4b-202 | 1 | 9.88 | 3.42 | 1.30E-03 | ENSRNOT00000041128 |
| Nectin2-203 | 1 | 10.45 | 1,558.51 | 0.02 | ENSRNOT00000117594 |
| Akt2-201 | 1 | 12.45 | 13.65 | 8.47E-04 | ENSRNOT00000025303 |
| Akt2-202 | 1 | 15.53 | -1,503.78 | 0.02 | ENSRNOT00000100544 |
| Slco2b1-203 | 1 | 15.63 | 2.3 | 4.95E-03 | ENSRNOT00000095615 |
| Numa1-203 | 1 | 6.22 | -32.28 | 3.14E-12 | ENSRNOT00000094333 |
| LOC100302465-202 | 1 | 9.2 | -3.66 | 0.03 | ENSRNOT00000073836 |
| Osbpl5-202 | 1 | 7.61 | -109.73 | 9.77E-09 | ENSRNOT00000079454 |
| Ndufv1-201 | 1 | 8.4 | -215.51 | 8.18E-04 | ENSRNOT00000024517 |
| Brms1-202 | 1 | 8.24 | -661.13 | 0.04 | ENSRNOT00000072647 |
| Rab1b-204 | 1 | 36.6 | 2.91 | 0.04 | ENSRNOT00000100410 |
| Anxa1-204 | 1 | 22 | 3.83 | 0.02 | ENSRNOT00000096367 |
| Zfp518a-204 | 1 | 7.71 | -2.38 | 0.03 | ENSRNOT00000105629 |
| Scd-201 | 1 | 11.4 | 2.49 | 1.76E-05 | ENSRNOT00000018447 |
| Cyp17a1-201 | 1 | 247.38 | 2.07 | 1.89E-05 | ENSRNOT00000027160 |
| Fam172a-203 | 2 | 7.34 | -8.02 | 9.46E-07 | ENSRNOT00000093741 |
| Zbed3-201 | 2 | 10.19 | -16.99 | 1.44E-03 | ENSRNOT00000040983 |
| Rps15al4-201 | 2 | 67.2 | 2.32 | 0.01 | ENSRNOT00000041483 |
| Rai14-201 | 2 | 12.3 | 2.24 | 2.70E-04 | ENSRNOT00000040348 |
| Pdzd2-203 | 2 | 6.64 | -2.52 | 0.02 | ENSRNOT00000105047 |
| Exosc9-203 | 2 | 8.84 | -808.62 | 0.03 | ENSRNOT00000108461 |
| Jade1-202 | 2 | 5.49 | -46.84 | 1.17E-10 | ENSRNOT00000098865 |
| Naa15-201 | 2 | 25.58 | -2.35 | 4.27E-05 | ENSRNOT00000017379 |
| Naa15-202 | 2 | 19.54 | 8.47 | 3.40E-10 | ENSRNOT00000095909 |
| Postn-204 | 2 | 23.97 | -12.54 | 1.16E-06 | ENSRNOT00000110085 |
| Postn-201 | 2 | 119.63 | -2.12 | 3.57E-15 | ENSRNOT00000017453 |
| Tm4sf1-201 | 2 | 29.09 | 2.07 | 6.02E-04 | ENSRNOT00000021535 |
| B3galnt1-202 | 2 | 10.46 | 2.62 | 0.03 | ENSRNOT00000095435 |
| Hsd3b3-201 | 2 | 436.49 | 2.25 | 2.51E-13 | ENSRNOT00000056172 |
| Hsd3b1-201 | 2 | 1,521.65 | 2.6 | 1.26E-21 | ENSRNOT00000086835 |
| Hao2-202 | 2 | 7.56 | 6 | 0.01 | ENSRNOT00000091444 |
| Hao2-201 | 2 | 14.1 | 376.32 | 3.87E-05 | ENSRNOT00000046942 |
| Ptgfrn-202 | 2 | 10.68 | 5.67 | 1.16E-13 | ENSRNOT00000096621 |
| Aimp1-203 | 2 | 5.77 | 364.3 | 0.04 | ENSRNOT00000112244 |
| Prkacb-202 | 2 | 10.02 | 2,519.69 | 8.45E-03 | ENSRNOT00000109432 |
| ENSRNOT00000108800 | 3 | 14,344.94 | -3.1 | 2.19E-08 | ENSRNOT00000108800 |
| ENSRNOT00000112407 | 3 | 7,081.59 | -3.02 | 1.92E-07 | ENSRNOT00000112407 |
| Ass1-202 | 3 | 50.03 | 2.24 | 2.48E-07 | ENSRNOT00000093365 |
| Rif1-202 | 3 | 10.52 | -2.26 | 1.44E-04 | ENSRNOT00000094327 |
| Fap-203 | 3 | 5.42 | -765.52 | 0.04 | ENSRNOT000096836 |
| Nfe2l2-203 | 3 | 11.16 | -21.56 | 1.42E-07 | ENSRNOT00000110389 |
| Ctnnd1-201 | 3 | 7.14 | -2.15 | 0.04 | ENSRNOT00000064009 |
| Fibin-201 | 3 | 10.55 | 2.9 | 6.48E-03 | ENSRNOT00000006203 |
| Snap23-201 | 3 | 5.8 | -45.5 | 0.01 | ENSRNOT00000074392 |

| | | | | | |
|---|---|---|---|---|---|
| Sema6d-203 | 3 | 6.33 | -8.26 | 1.16E-08 | ENSRNOT00000098848 |
| ENSRNOT00000028881 | 3 | 88.78 | 2.22 | 0.02 | ENSRNOT00000028881 |
| Chd6-202 | 3 | 10 | -2.39 | 4.20E-03 | ENSRNOT00000089958 |
| Ythdf1-201 | 3 | 12.8 | -2.58 | 0.01 | ENSRNOT00000037271 |
| Fastk-204 | 4 | 15.8 | 284.4 | 2.78E-05 | ENSRNOT00000118434 |
| RGD1565355-201 | 4 | 5.63 | -455.94 | 0.02 | ENSRNOT00000008319 |
| Capza2-201 | 4 | 58.91 | 2.02 | 2.18E-04 | ENSRNOT00000090451 |
| Podxl-201 | 4 | 8.43 | 68.83 | 9.99E-16 | ENSRNOT00000016991 |
| Akr1b10-204 | 4 | 20.44 | 2.55 | 0.04 | ENSRNOT00000116609 |
| Ttll3-203 | 4 | 6.95 | -2.95 | 0.01 | ENSRNOT00000107721 |
| Atp6v1e1-202 | 4 | 18.38 | 12.82 | 0.04 | ENSRNOT00000098217 |
| Plbd1-201 | 4 | 34.38 | 2.39 | 1.42E-07 | ENSRNOT00000011930 |
| Slc35a1-201 | 5 | 10.19 | 1,167.15 | 6.03E-03 | ENSRNOT00000011969 |
| Ncbp1-202 | 5 | 12.69 | 4.96 | 0.05 | ENSRNOT00000094126 |
| Ptprd-203 | 5 | 7.17 | -2.22 | 0.02 | ENSRNOT00000110800 |
| Faf1-203 | 5 | 5.38 | -24.21 | 0.02 | ENSRNOT00000108187 |
| LOC100294508-202 | 5 | 5.21 | 275.11 | 1.63E-05 | ENSRNOT00000100704 |
| Eif4g3-209 | 5 | 10.11 | -15.38 | 4.63E-03 | ENSRNOT00000115071 |
| LOC100359951-201 | 5 | 332.09 | 2.03 | 5.02E-05 | ENSRNOT00000040481 |
| Ccnl2-203 | 5 | 26.43 | -3.4 | 1.26E-05 | ENSRNOT00000089936 |
| Ints11-203 | 5 | 5.44 | 35.37 | 0.01 | ENSRNOT00000107036 |
| Xdh-201 | 6 | 7.76 | 2.4 | 4.09E-03 | ENSRNOT00000009634 |
| Tpo-201 | 6 | 13.16 | -2.01 | 0.02 | ENSRNOT00000006526 |
| Lamb1-201 | 6 | 22.22 | -3.01 | 7.83E-06 | ENSRNOT00000008321 |
| Rdh11-203 | 6 | 6.54 | -565.73 | 0.01 | ENSRNOT00000105155 |
| Zfp410-204 | 6 | 7.23 | 7.67 | 0.03 | ENSRNOT00000109998 |
| Eml5-201 | 6 | 5.64 | -3.52 | 1.00E-03 | ENSRNOT00000057601 |
| Papola-202 | 6 | 6.22 | 184.13 | 8.47E-04 | ENSRNOT00000083732 |
| Klc1-202 | 6 | 5.64 | 217.79 | 7.71E-04 | ENSRNOT00000065281 |
| Klc1-205 | 6 | 5.07 | -3.47 | 0.03 | ENSRNOT00000118156 |
| ENSRNOT00000109415 | 6 | 26.33 | 19.63 | 2.14E-03 | ENSRNOT00000109415 |
| Mta1-201 | 6 | 12.48 | 2.28 | 8.03E-03 | ENSRNOT00000006521 |
| Rab5b-203 | 7 | 15.84 | 6.97 | 4.82E-06 | ENSRNOT00000120020 |
| Eef2-202 | 7 | 7.6 | 2.62 | 0.05 | ENSRNOT00000096642 |
| Eea1-202 | 7 | 5.43 | 9.21 | 9.60E-03 | ENSRNOT00000087297 |
| Eea1-206 | 7 | 11.8 | -2.22 | 5.66E-03 | ENSRNOT00000117665 |
| Csrp2-202 | 7 | 86 | 2.11 | 9.70E-11 | ENSRNOT00000080598 |
| Lyz2-201 | 7 | 81.33 | 2.05 | 3.73E-07 | ENSRNOT00000007747 |
| Ubr5-201 | 7 | 14.12 | 2.02 | 3.04E-04 | ENSRNOT00000009115 |
| Angpt1-201 | 7 | 10.55 | 2.06 | 0.03 | ENSRNOT00000007979 |
| Eif3e-201 | 7 | 11.83 | 6.64 | 0.04 | ENSRNOT00000038340 |
| Mtss1-205 | 7 | 22.29 | -2.19 | 5.69E-07 | ENSRNOT00000118933 |
| Mtss1-201 | 7 | 6.43 | 5.38 | 8.33E-03 | ENSRNOT00000011984 |
| Cyb5r3-202 | 7 | 17.96 | 2.66 | 4.76E-03 | ENSRNOT00000102247 |
| Igfbp6-201 | 7 | 245.69 | 2.38 | 1.65E-15 | ENSRNOT00000014807 |
| Hoxc6-202 | 7 | 5.93 | -2.33 | 0.01 | ENSRNOT00000107142 |
| Ldlr-202 | 8 | 7.47 | 10.48 | 5.78E-03 | ENSRNOT00000108195 |
| Acp5-202 | 8 | 5.13 | 117.22 | 6.39E-03 | ENSRNOT00000092200 |
| LOC120094123-202 | 8 | 39.08 | -2.17 | 0.02 | ENSRNOT00000111823 |
| Btg4-202 | 8 | 21.66 | 2.24 | 0.04 | ENSRNOT00000088320 |
| Cyp11a1-201 | 8 | 25.37 | 3.12 | 1.32E-05 | ENSRNOT00000010831 |

| | | | | | |
|---|---|---|---|---|---|
| Cyp11a1-202 | 8 | 29.18 | 4.17 | 8.25E-06 | ENSRNOT00000106594 |
| Arih1-202 | 8 | 5.88 | -467.69 | 0.02 | ENSRNOT00000084758 |
| Clpx-204 | 8 | 7.07 | 10.14 | 0.05 | ENSRNOT00000110101 |
| Tln2-202 | 8 | 5.09 | -2.23 | 0.03 | ENSRNOT00000024885 |
| Tln2-204 | 8 | 5.27 | 2.02 | 0.04 | ENSRNOT00000064613 |
| Snx14-201 | 8 | 32.41 | -2.16 | 1.22E-03 | ENSRNOT00000056818 |
| Morf4l1-201 | 8 | 35.45 | 2,995.97 | 6.42E-03 | ENSRNOT00000080421 |
| Slc25a36-202 | 8 | 8.15 | -16.73 | 0.01 | ENSRNOT00000110158 |
| Ifrd2-203 | 8 | 9.71 | 1,044.36 | 0.03 | ENSRNOT00000112128 |
| AABR07071482.2-201 | 8 | 10.03 | -39.06 | 0.03 | ENSRNOT00000047247 |
| Gtpbp2-203 | 9 | 7.54 | -2.96 | 0.03 | ENSRNOT00000094924 |
| Map4k4-203 | 9 | 7.78 | -2.3 | 0.05 | ENSRNOT00000102137 |
| Myo1b-205 | 9 | 5.51 | 4.59 | 0.03 | ENSRNOT00000118845 |
| Cnot9-203 | 9 | 10.44 | 11.26 | 1.74E-03 | ENSRNOT00000107071 |
| Ptprn-201 | 9 | 10.28 | 2.13 | 0.03 | ENSRNOT00000026654 |
| Col6a3-201 | 9 | 5.38 | -2.73 | 0.02 | ENSRNOT00000026719 |
| Abcc1-203 | 10 | 7.53 | 250.72 | 3.35E-06 | ENSRNOT00000115220 |
| Myh11-201 | 10 | 10.16 | 2.3 | 7.41E-05 | ENSRNOT00000084608 |
| Srrm2-201 | 10 | 38.7 | -5.71 | 0.01 | ENSRNOT00000091287 |
| Hba-a3-201 | 10 | 200.6 | 2.03 | 1.21E-05 | ENSRNOT00000048977 |
| Myh10-205 | 10 | 12.41 | -4.79 | 1.19E-05 | ENSRNOT00000113616 |
| Alox15-201 | 10 | 51.46 | 2.08 | 1.24E-03 | ENSRNOT00000026038 |
| Camta2-203 | 10 | 5.24 | 6.51 | 0.01 | ENSRNOT00000109888 |
| P2rx1-202 | 10 | 15.38 | -4.8 | 1.31E-03 | ENSRNOT00000088245 |
| P2rx1-203 | 10 | 8.31 | -1,168.29 | 3.37E-03 | ENSRNOT00000118157 |
| P2rx1-201 | 10 | 7.2 | 28.26 | 3.67E-05 | ENSRNOT00000024068 |
| Arhgap23-203 | 10 | 10.85 | 2.23 | 5.80E-03 | ENSRNOT00000100892 |
| Hsd17b1-201 | 10 | 108.31 | 4.34 | 2.77E-37 | ENSRNOT00000026878 |
| Nsf-203 | 10 | 5.77 | 39.38 | 2.37E-05 | ENSRNOT00000099576 |
| Nploc4-201 | 10 | 17.65 | 6.54 | 1.38E-04 | ENSRNOT00000054973 |
| Abi3bp-205 | 11 | 11.86 | -3.61 | 8.82E-05 | ENSRNOT00000063864 |
| Cd200-203 | 11 | 23.76 | -4.34 | 8.46E-03 | ENSRNOT00000093295 |
| Dgcr2-203 | 11 | 15.85 | 4.62 | 0.02 | ENSRNOT00000107429 |
| Smpd4-202 | 11 | 7.13 | -3.87 | 0.02 | ENSRNOT00000102059 |
| Rilpl1-202 | 12 | 17.26 | 2.02 | 0.04 | ENSRNOT00000090831 |
| Rab35-202 | 12 | 16.49 | 4.49 | 2.45E-03 | ENSRNOT00000103324 |
| Rab35-203 | 12 | 6.74 | -798.79 | 0.03 | ENSRNOT00000106644 |
| Pxn-201 | 12 | 13.32 | -3.86 | 8.51E-05 | ENSRNOT00000047466 |
| Pxn-203 | 12 | 11.5 | 6.99 | 2.89E-04 | ENSRNOT00000081134 |
| Tnnt2-204 | 13 | 6.15 | 7.8 | 0.04 | ENSRNOT00000108522 |
| Arpc5-204 | 13 | 14.96 | 1,542.52 | 0.03 | ENSRNOT00000107757 |
| Dars2-201 | 13 | 5.16 | 355.34 | 0.04 | ENSRNOT00000003828 |
| Mael-201 | 13 | 66.36 | -2.02 | 1.09E-05 | ENSRNOT00000005060 |
| Copa-203 | 13 | 5.81 | -3.31 | 8.83E-03 | ENSRNOT00000109085 |
| Dcaf8-203 | 13 | 6.27 | 26.15 | 8.84E-03 | ENSRNOT00000101398 |
| Ephx1-201 | 13 | 80.11 | 2.03 | 1.21E-05 | ENSRNOT00000004780 |
| Cd46-203 | 13 | 29.87 | -2.36 | 0.01 | ENSRNOT00000079549 |
| Cd46-204 | 13 | 44.04 | 2.06 | 2.27E-05 | ENSRNOT00000084320 |
| Sec31a-201 | 14 | 10.9 | 2.08 | 0.02 | ENSRNOT00000003072 |
| Bmp3-202 | 14 | 23.4 | -2.06 | 2.95E-03 | ENSRNOT00000104172 |
| Cxcl9-201 | 14 | 39.02 | 2.07 | 6.98E-03 | ENSRNOT00000003082 |

| | | | | | |
|---|---|---|---|---|---|
| Gnrhr-201 | 14 | 20.34 | 3.39 | 8.45E-07 | ENSRNOT00000002755 |
| Ociad1-202 | 14 | 15.62 | 926.17 | 0.03 | ENSRNOT00000094276 |
| Limch1-202 | 14 | 5.71 | -2.7 | 0.03 | ENSRNOT00000095221 |
| Dlg5-204 | 15 | 8.1 | -2.22 | 4.87E-03 | ENSRNOT00000112381 |
| Lgals3-202 | 15 | 48.99 | 2.75 | 2.75E-06 | ENSRNOT00000082304 |
| Slc22a17-202 | 15 | 8.19 | 914.7 | 0.04 | ENSRNOT00000102200 |
| Mcpt1l1-201 | 15 | 25.88 | -2.83 | 0.02 | ENSRNOT00000090206 |
| Ints9-203 | 15 | 10.48 | 2.8 | 0.03 | ENSRNOT00000096924 |
| Ephx2-201 | 15 | 22.06 | 2 | 0.03 | ENSRNOT00000023385 |
| Plac9-201 | 16 | 103.51 | 2.01 | 1.59E-04 | ENSRNOT00000014964 |
| Hpgd-201 | 16 | 39.4 | 2.51 | 1.88E-05 | ENSRNOT00000014229 |
| Fat1-203 | 16 | 9.84 | 2.15 | 0.05 | ENSRNOT00000095529 |
| Ogn-202 | 17 | 268.51 | 2.1 | 1.86E-09 | ENSRNOT00000093561 |
| Sycp2l-201 | 17 | 23.45 | -2.08 | 4.78E-03 | ENSRNOT00000038180 |
| Exoc2-205 | 17 | 5.76 | 1,522.67 | 0.02 | ENSRNOT00000092101 |
| Sfrp4-202 | 17 | 29.51 | 3.62 | 5.72E-11 | ENSRNOT00000079368 |
| Inhba-201 | 17 | 140.94 | 2.08 | 1.65E-07 | ENSRNOT00000019272 |
| Ccdc3-201 | 17 | 19.83 | 2.01 | 3.50E-03 | ENSRNOT00000024110 |
| Npc1-202 | 18 | 10.93 | 2.55 | 0.04 | ENSRNOT00000104000 |
| Hdac3-201 | 18 | 11.4 | -5.56 | 4.78E-03 | ENSRNOT00000060417 |
| Txnl1-204 | 18 | 12.21 | 165.54 | 6.93E-04 | ENSRNOT00000100860 |
| Zbtb7c-202 | 18 | 9.55 | 37.86 | 1.84E-13 | ENSRNOT00000101836 |
| Gpt2-203 | 19 | 8.85 | 5.84 | 0.02 | ENSRNOT00000099582 |
| Adgrl1-206 | 19 | 7.59 | -6.17 | 1.93E-03 | ENSRNOT00000103294 |
| Adgrl1-204 | 19 | 11.3 | -2.5 | 5.64E-04 | ENSRNOT00000101340 |
| Dpep1-201 | 19 | 74.12 | 2.05 | 2.52E-08 | ENSRNOT00000021397 |
| Rab4a-202 | 19 | 6.09 | 498.21 | 0.02 | ENSRNOT00000098153 |
| Syngap1-206 | 20 | 7.43 | -2.3 | 0.03 | ENSRNOT00000120108 |
| Tead3-203 | 20 | 9.78 | -8.11 | 3.97E-03 | ENSRNOT00000107495 |
| Srpk1-204 | 20 | 7.21 | -55.13 | 3.97E-03 | ENSRNOT00000100033 |
| Lims1-201 | 20 | 10.7 | -8.84 | 5.15E-03 | ENSRNOT00000064288 |
| Mageb6b1-201 | X | 5.86 | -114.39 | 2.90E-03 | ENSRNOT00000043817 |
| Ogt-201 | X | 59.06 | -2.2 | 0.02 | ENSRNOT00000004692 |
| Ogt-203 | X | 10.8 | 11.02 | 0.04 | ENSRNOT00000100805 |
| Taf9b-201 | X | 5.9 | -1,069.45 | 6.43E-03 | ENSRNOT00000090833 |
| Cstf2-202 | X | 11.02 | -2.22 | 0.05 | ENSRNOT00000079572 |
| Armcx3-201 | X | 5.08 | -111.11 | 1.23E-03 | ENSRNOT00000029664 |
| Il13ra1-ps1-202 | X | 11.32 | 22.62 | 2.90E-03 | ENSRNOT00000112175 |
| Zcchc12-202 | X | 11.49 | 4.18 | 0.05 | ENSRNOT00000094904 |
| Xiap-203 | X | 11.18 | -3.04 | 5.88E-03 | ENSRNOT00000085110 |
